# The origin and molecular evolution of the mammalian liver cell architecture

**DOI:** 10.1101/2025.10.14.682049

**Authors:** Xuefei Yuan, Leticia Rodríguez-Montes, Bastienne Zaremba, Nils Trost, Céline Schneider, Julia Schmidt, Bianka Berki, David Ibberson, Evgeny Leushkin, Birgit Nickel, Miklós Palkovits, Richard W. Truman, Greg Barsh, John Lees, Amir Fallahshahroudi, Frank Grützner, Athanasia C. Tzika, Michel C. Milinkovitch, Svante Pääbo, Margarida Cardoso-Moreira, Henrik Kaessmann

**Author notes:** Present address: LOEWE Center for Translational Biodiversity Genomics, Frankfurt, Germany (Evgeny Leushkin). These authors contributed equally (Xuefei Yuan, Leticia Rodríguez-Montes). e-mail: X.Y.; L.R.-M.; H.K.

## Abstract

The liver is a central organ with essential roles in processes such as nutritional metabolism, detoxification, and immune defense^1–6^. It has been instrumental in the adaptation of mammalian species to diverse environments, as reflected by its rapid molecular evolution^7,8^. However, the origins and evolutionary dynamics of liver cell types and their structural organization remain largely unexplored. Here we report evolutionary analyses of transcriptome and chromatin accessibility data for liver cells from 17 species, spanning all major feeding strategies (herbivory, omnivory, carnivory, insectivory), great apes (including humans), placental clades (Afrotheria, Xenarthra, Laurasiatheria, Euarchontoglires), major mammalian lineages (placentals, marsupials, monotremes), and a bird as outgroup. Integrated with spatial transcriptomics, our data reveal that liver zonation—the compartmentalization of hepatocyte functions along the lobule, the liver’s fundamental anatomical and functional unit—is conserved across mammals but absent in other vertebrates. We find that zonation originated in the mammalian ancestor, driven by the emergence of WNT and R-spondin signaling from central vein endothelial cells, which activate central hepatocyte gene expression via the transcription factor TCF7L2. Despite this conserved architecture and signaling, genes with zonated expression exhibit rapid evolutionary turnover. Consistently, hepatocytes evolve fast, likely due to reduced selective constraints, enabling adaptive changes under positive selection. Alongside immune cells, hepatocytes are therefore key drivers of the liver’s rapid evolution and functional innovations. In great apes, we identify human-specific shifts in zonation and cell-type-specific expression linked to recent cis-regulatory changes, particularly in genes involved in lipid metabolism, likely contributing to human-specific metabolic traits. Our study uncovers the origins of a mammal-specific liver cell architecture, within which reduced constraints facilitated molecular changes underlying ecological adaptations.

---

Mammals have adapted to diverse and extreme environments, with the liver playing a central role through its wide range of functions, including nutrient metabolism, detoxification, pathogen clearance, regulation of homeostasis, coagulation, and thermogenesis^1–6^. Despite its ancient origin in the vertebrate stem lineage^9–11^ and essential roles, the liver expresses a high proportion of evolutionarily young genes^7,8^ and genes under positive selection^8^ compared to other mammalian organs, as assessed from transcriptomic investigations of bulk-tissue samples. This rapid molecular evolution suggests that the liver has been pivotal for mammalian adaptations; yet the molecular and gene regulatory changes shaping its evolution across different mammalian lineages remain unexplored at the cellular level.

Investigations of placental mammals revealed that hepatocytes, the liver’s primary cells, are organized into specialized polygonal units, typically 0.5–1 mm in diameter, called lobules^12,13^. Blood enters the lobule through the portal vein and hepatic artery and flows toward the central vein, establishing gradients of oxygen, nutrients, and signaling molecules. These gradients underlie liver zonation, the spatial compartmentalization of hepatocyte metabolic functions in distinct lobule regions^14^. The liver’s basic vascular organization is conserved across vertebrates^15–18^, suggesting that it arose in the common vertebrate ancestor. The origination and evolutionary dynamics of zonation have not been studied.

Here we present gene expression and chromatin accessibility measurements of liver cells from 16 representative mammals and a bird, complemented by spatial transcriptomics in key species (https://apps.kaessmannlab.org/liver_app/). Integrated with single-cell expression data from a vertebrate outgroup, these analyses unveiled the evolutionary origins of zonation, along with the ancestral and species-specific gene expression programs of mammalian liver cell types and their regulatory foundations.

## Cellular multiomic atlases of mammalian livers

To investigate the molecular, cellular, and regulatory mechanisms underlying the rapid evolution of mammalian livers, we generated single-nucleus RNA sequencing (snRNA-seq) atlases for 16 mammals, covering all major feeding strategies, great apes, placental clades, and major mammalian lineages (Fig. 1a). For 13 species, we also obtained matching chromatin accessibility data through separate single-nucleus ATAC-seq (snATAC-seq) or joint single-nucleus multiome profiling (Fig. 1a and Supplementary Table 1). In addition, we generated multiome data for chicken as an evolutionary outgroup. To optimize read mapping and reduce gene detection bias in cross-species comparisons, we refined and extended existing transcriptome annotations for non-model organisms using bulk or snRNA-seq liver data (Methods and Supplementary Table 2).

**Figure 1.**
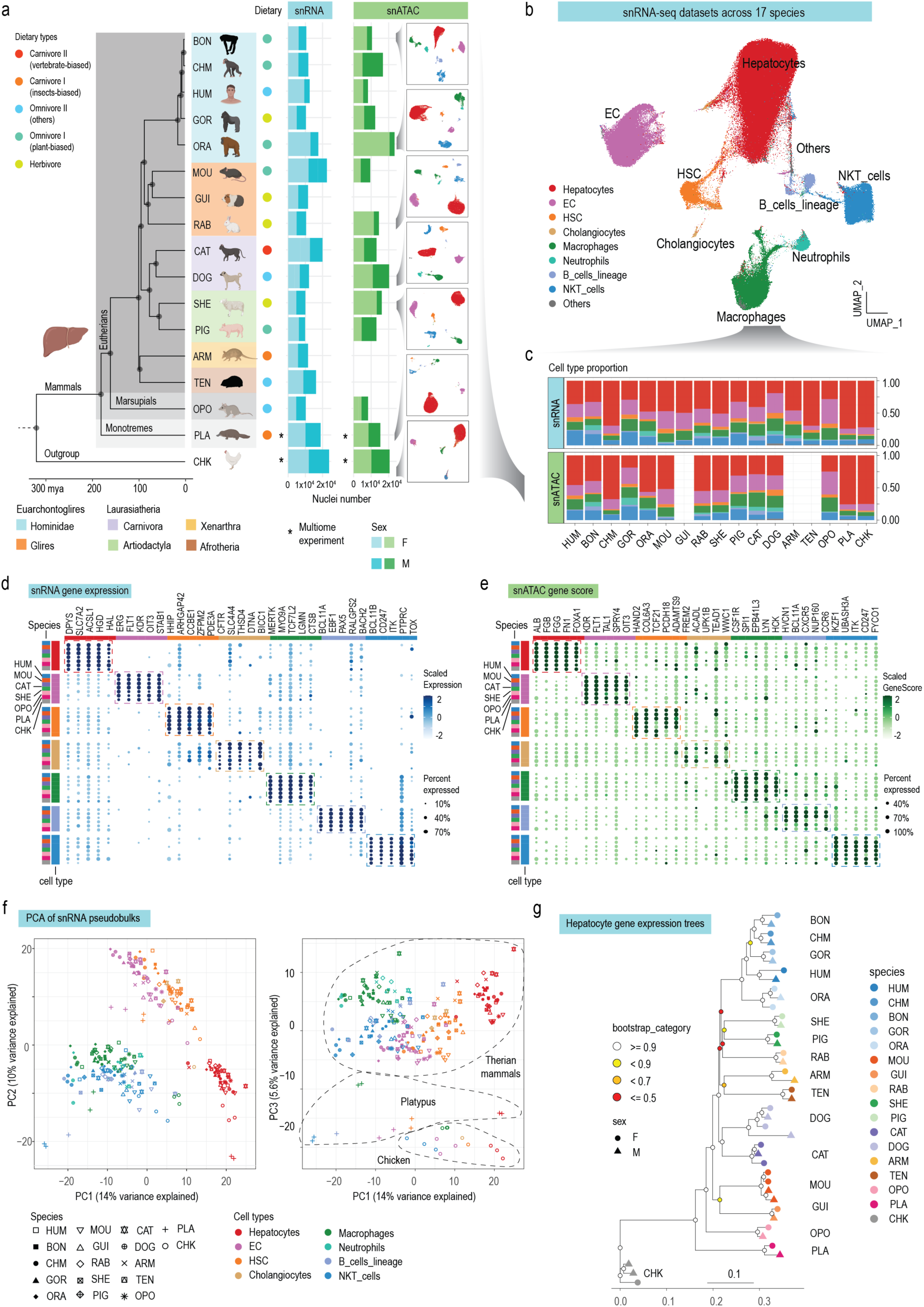
| Multiomic cellular atlases of mammalian livers. **a**, Species and number of nuclei sampled for snRNA-seq and snATAC-seq, and UMAP representation of the snATAC-seq data of 7 species, colored by cell types (left). The dietary labels are derived from the EltonTraits 1.0 dataset. Star signs indicate joint-profiling of RNA and ATAC from the same nuclei for platypus and chicken samples. mya: million years ago. **b**, UMAP visualization of integrated snRNA-seq data across all 17 species. Cells are colored by their original annotation in each species. **c**, Cell composition of the snRNA-seq and snATAC-seq datasets in each species. **d**, **e**, Dotplot showing the scaled gene expression from snRNA-seq (**d**) or scaled gene scores from snATAC-seq (**e**) of the top 5 conserved markers in 7 representative species. Only the gene expression/score that is detected in no less than 10% of the nuclei is plotted. **f**, PCA of pseudobulk profiles aggregated from snRNA-seq data across all 17 species based on variable and robustly expressed 1:1 orthologous genes. Each data point represents cell-type pseudobulks from one individual. Dotted lines encircle species or lineages. **g,** Gene expression tree based on pseudobulk transcriptomes for hepatocytes. Bootstrap values (3,399 1:1 orthologous genes were randomly sampled with replacement 1,000 times) are indicated by circles. The species icons were created in BioRender. Rodriguez, L. (2025) https://BioRender.com/gwk4qwp.

Our datasets include both sexes and 2–4 biological replicates per assay per species (Fig. 1a and Extended Fig. 1). After quality control, we obtained transcriptomic profiles for 239,512 nuclei and chromatin accessibility profiles for 201,147 nuclei. The transcriptomic dataset averaged 14,089 nuclei per species, with a median of 3,572 RNA molecules (UMIs) and 1,970 genes detected per nucleus (Extended Fig. 1 and Supplementary Table 3). Chromatin accessibility profiling averaged 14,368 nuclei per species, with a median of 10,392 fragments per nucleus (Extended Fig. 1 and Supplementary Table 4).

Using marker gene-based annotation (Methods), we identified all major liver cell types in the transcriptomic and chromatin accessibility datasets across all 17 species: hepatocytes, endothelial cells (ECs), resident macrophages, hepatic stellate cells (HSCs), and a mixed group of T cells and NK cells (collectively referred to as NKT cells) (Fig. 1b-e and Extended Data Fig. 2, 3). Smaller populations, including cholangiocytes, B cells, and neutrophils, were detected in most species (Fig. 1c and Extended Data Fig. 2, 3). Cross-species transcriptome integration confirmed these annotations and showed that liver cell types are conserved among amniotes (Fig. 1b, c).

To compare transcriptomes across cell types, species, and replicates, we generated pseudobulk profiles for each cell type and replicate and performed principal component analysis (PCA) using robustly expressed 1:1 orthologous genes. Gene expression variance is explained primarily by cell type (PC1 and PC2), followed by evolutionary lineage (PC3) (Fig. 1f). We also reconstructed gene expression trees from the pseudobulks (Fig. 1g and Extended Data Fig. 4; Methods), which broadly recapitulated the mammalian phylogeny (eutherians, marsupials, monotremes). Species from the same order or family mostly clustered together, with rodents as a notable exception (not clustering with their sister group), potentially reflecting lineage-specific biology. Bootstrap analysis showed weak support for the phylogeny of Afrotheria, Xenarthra, Euarchontoglires, and Laurasiatheria (Fig. 1g and Extended Data Fig. 4), likely due to the rapid early radiation of placental mammals that complicates phylogenetic reconstruction. Together, these analyses show that our data capture both ancestral liver gene expression programs as well as species– and lineage-specific divergences.

To assess the conservation of gene regulation across liver cell types, we analyzed chromatin accessibility profiles for each species, identifying accessible peaks as putative cis-regulatory elements (CREs). We detected a median of 322,091 putative CREs per species, with 37.6% being distal intergenic and 49.3% intronic (Extended Data Fig. 5a). To pinpoint the transcription factors (TFs) driving cell-type-specific programs, we focused on cell-type specific CREs and examined enriched TF binding motifs. Major cell types showed consistent motif enrichment across amniotes: HNF, RXR, and CEBP families in hepatocytes; ETV, ELF, and SOX in endothelial cells; SP1, ELF, and IRF in macrophages; and RUNX1, TBX, and EOMES in NKT cells (Extended Data Fig. 5b), indicating strong conservation of liver trans-regulatory programs across amniotes^19–21^. We detected small differences in platypus hepatocytes and chicken NKT cells, which displayed altered motif rankings (Extended Data Fig. 5b-d), suggesting some lineage-specific variation.

Together, the transcriptome and chromatin accessibility profiles of liver cells reveal a core hepatic gene expression program conserved across amniotes.

## Rates of cell type evolution

Previous studies showed that the liver evolves rapidly in mammals at the molecular level^7,8,22,23^, but it is unknown which cell types and evolutionary forces drive this rapid evolution. We thus compared transcriptome similarity (UMI– and cell number-controlled) between human and eight species across a range of evolutionary distances. As expected, similarity declined with increasing evolutionary time but saturated at ∼200 million years: differences between human and platypus were comparable to those between human and chicken, despite birds diverging ∼110 million years earlier (Fig. 2a). This saturation, also seen at the whole-organ level^22^, likely reflects a conserved core of liver functions under strong stabilizing selection. Notably, hepatocytes showed the lowest transcriptome similarity between human and most (6/8) other species (Fig. 2a), suggesting that they drive the liver’s rapid evolution.

**Figure 2.**
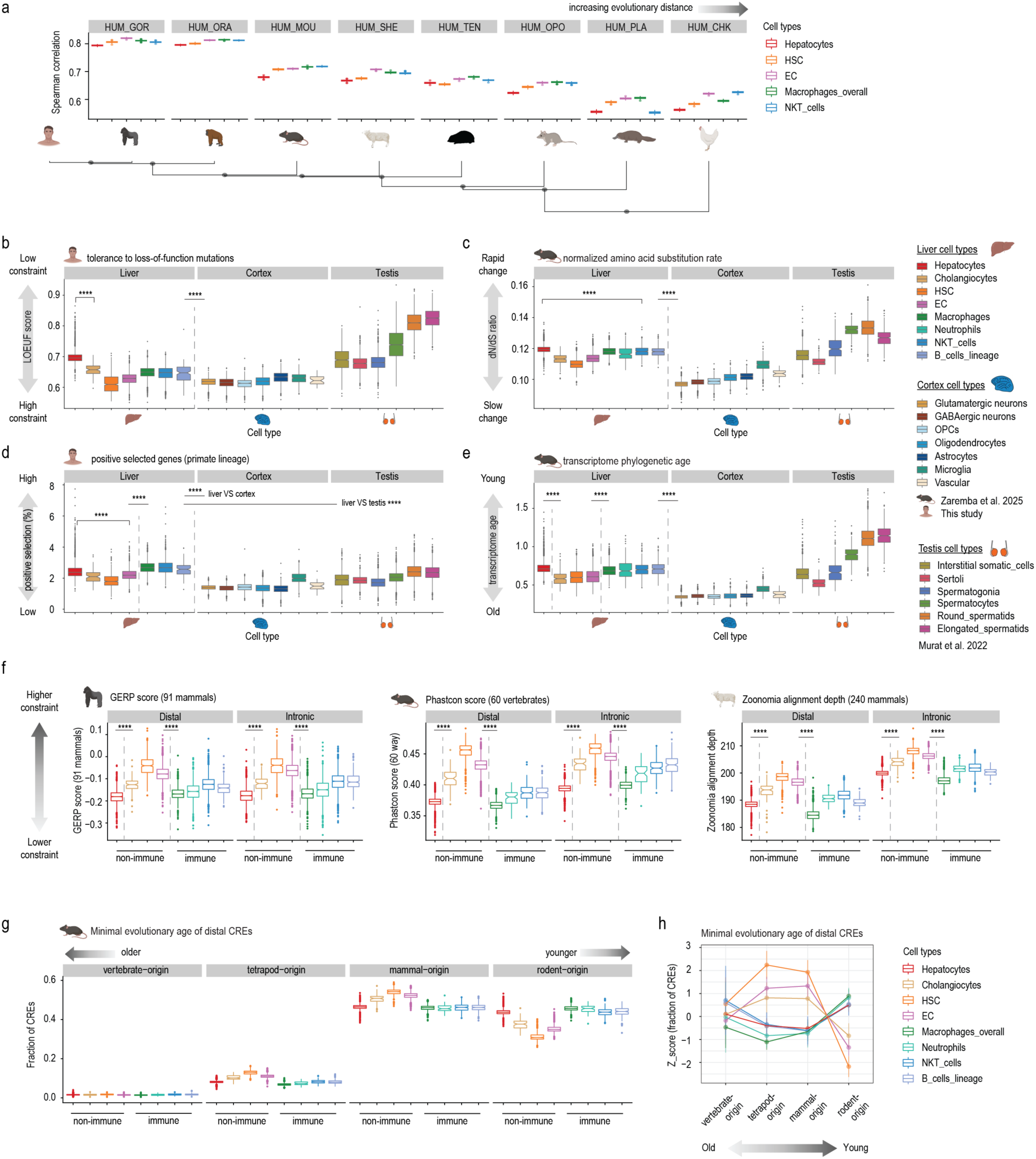
| Evolutionary rates of mammalian liver cell types. **a**, Pair-wise correlation between human and the other 8 species after UMI and cell-number controlled downsampling, with 100 times bootstrapping. **b**, LOEUF (loss-of-function observed/expected upper bound fraction) scores of genes expressed in each cell type in human. High LOEUF scores indicate high tolerance to loss-of-function mutations. **c**, d*N*/d*S* ratio (across 8 mammals) of genes expressed in each cell type in mouse. **d**, Percentage of expressed genes with signs of being positively selected in the primate lineage. **e**, Transcriptome phylogenetic ages of genes expressed in each cell type in mouse. **f**, GERP scores (left), Phastcon scores (middle), and Zoonomia alignment depths (right) of the putative distal or intronic CREs active in each cell type in gorilla (left), mouse (middle), and sheep (right). **g**, Fraction of distal CREs with different minimal evolutionary ages in mouse liver cells. **h**, Z-score of the fraction of distal CREs with different minimal evolutionary ages scaled across all mouse nuclei. **b-f**, Sources of the data and metrics used in the plots are described in Methods. Each data point represents a nucleus, with the average value of the genes or putative CREs detected in the nucleus plotted. **a-g**, Boxplots display the median (center value) and upper and lower quantiles (box limits) with whiskers at 1.5 times the interquartile ranges, ****: *P* < 0.0001.

To examine hepatocyte evolution in a broader cellular context, we incorporated previously available and newly generated snRNA-seq data from one of the slowest-evolving organs, the brain (cortex), and one of the fastest, the testis^7,8,22^. We assessed functional constraint in genes expressed in main cell types of the liver, cortex, and testis using three independent metrics: embryonic lethality phenotypes in mouse knockouts^24^, tolerance to loss-of-function mutations in human populations^25^, and coding sequence constraint across 240 mammals^26^ (Methods, Fig. 2b, Extended Data Fig. 6a-c, and Supplementary Table 5). All three measures indicated that hepatocyte-expressed genes are under the weakest constraint in the liver (Mann-Whitney *U* test, *P* < 10^−11^), followed by immune cell genes. Across the three organs, human liver cells are under lower constraint than cortical cells (Mann-Whitney *U* test, *P* < 10^−100^) but more constrained than spermatocytes and spermatids, which are known for their highly rapid evolution (Fig. 2b and Extended Data Fig. 6a).

To assess constraint patterns across different evolutionary lineages, we calculated the normalized rate of amino acid altering substitutions (d*N*/d*S* ratio) for protein-coding genes in mammals, primates, and great apes and compared liver cell transcriptomes in 11 species (Methods). Hepatocyte transcriptomes exhibited the highest substitution rates across all species and lineages examined (Fig. 2c and Extended Data Fig. 6d-f, Mann-Whitney *U* test, *P* < 10^−9^). Elevated rates may reflect weaker purifying selection, consistent with the lower functional constraints observed, but could also indicate stronger positive selection. To test for this possibility, we assessed the proportion of expressed genes in each liver cell type with signatures of positive selection in primates^27^ or mammals^28^. Liver cells overall expressed a higher proportion of positively selected genes than cortical cells, and surprisingly, even than testicular cells in most species (Fig. 2d and Extended Data Fig. 6g-j, Mann-Whitney *U* test, *P* < 10^−13^). Immune cells showed the strongest signals of positive selection (Fig. 2d and Extended Data Fig. 6g-j, Mann-Whitney *U* test *P* < 10^−15^), consistent with their known evolutionary dynamics^29,30^. Among non-immune cells, hepatocytes expressed the highest proportion of positively-selected genes in great apes, especially when considering genes under positive selection in the primate lineage (Fig. 2d and Extended Data Fig. 6g,h, Mann-Whitney *U* test *P* < 10^−5^), a pattern not observed in non-primate mammals (Extended Data Fig. 6i,j). These results suggest that liver-expressed genes, especially those in immune cells, are strong targets of positive selection across mammals.

New genes are key contributors to phenotypic innovations, so we examined the use of evolutionarily young genes in liver cell types (Methods). Hepatocytes and immune cells expressed a higher fraction of recently evolved genes than other liver cell types (Fig. 2e and Extended Data Fig. 6k, Mann-Whitney *U* test *P* < 10^−160^). Cellular transcriptomes of the liver are overall younger than those of the brain (Mann-Whitney *U* test *P* < 10^−200^), consistent with previous work^7,8^, but older than those of late spermatogenic cells.

We next focused on the evolution of regulatory regions. We found that putative distal and intronic CREs active in hepatocytes and immune cells are under lower sequence constraint than those in other liver cell types across all species (Fig. 2f and Extended Data Fig. 7a-r, *P* < 10^−57^). To assess how CREs of different evolutionary ages contribute to liver gene regulation, we inferred the minimal age of each CRE in mouse from syntenic sequence alignments and categorized them as originating in the common ancestor of vertebrates, tetrapods, mammals, or rodents (Methods, Extended Data Fig. 7s). All liver cell types contained a similarly small fraction of vertebrate-origin distal CREs (1.6%), but the proportions of more recently evolved CREs varied markedly across cell types (Fig. 2g and Extended Data Fig. 7t). Hepatic stellate cells and endothelial cells showed higher proportions of tetrapod– and mammal-origin CREs, whereas hepatocytes and immune cells use a higher percentage of rodent-specific CREs (Fig. 2h and Extended Data Fig. 7u, Mann-Whitney *U* test, *P* < 10^−213^), suggesting a substantial contribution of evolutionarily young CREs to the evolution of liver functions, consistent with previous bulk-tissue analyses^23^.

Overall differences in the breadth of gene expression (expression pleiotropy) between cell types could explain both decreased functional constraints and increased adaptive signals^8,31,32^, as observed in hepatocytes and immune cells. To assess expression pleiotropy, we used multiple specificity indices, providing complementary information at cellular, tissue, and temporal levels (Methods). Across most species, hepatocyte transcriptomes exhibited the highest spatiotemporal specificity among liver cell types, reflecting the lowest pleiotropy (Extended Data Fig. 6l-q, Mann-Whitney *U* test, *P* < 10^−7^). These data suggest that the hepatocytes’ lower pleiotropy underlies both their reduced functional constraint and elevated adaptation rates.

Our analyses show that hepatocytes and immune cells are the primary drivers of the liver’s rapid molecular evolution. Increased usage of evolutionarily young genes (especially in hepatocytes and immune cells) and the accumulation of positively selected genes (most pronounced in immune cells) have fueled mammalian liver evolution.

## Origination of liver zonation

Liver zonation (zonated gene expression) is well documented in placental mammals, including humans, cynomolgus monkeys, mice, rats, and pigs^12,33–35^. In non-eutherian vertebrates, findings are mixed: no zonated gene expression has been observed in hagfish, zebrafish, Texas tortoises, or Argentine tree frogs^15,36,37^, while there is limited evidence of zonation of certain enzymes in the trout liver^38^. However, studies in non-mammalian species typically examined only a few genes rather than genome-wide patterns, leaving the origin and distribution of zonation across vertebrates poorly understood.

Based on marker gene expression, we found clear hepatocyte zonation in snRNA-seq data across all 16 mammalian species (Fig. 3a-b, Extended Data Fig. 8a), but not in chicken. Unsupervised clustering of mammalian hepatocytes consistently revealed sub-clusters with high expression of known central or portal genes. In chicken, however, no such sub-clusters were detected, and zonation genes conserved across distant mammalian lineages (e.g., *GLUL*, *SLC1A2*, *GLDC*, etc.) showed no cell-population bias, in line with previous observations^36^ (Fig. 3b).

**Figure 3.**
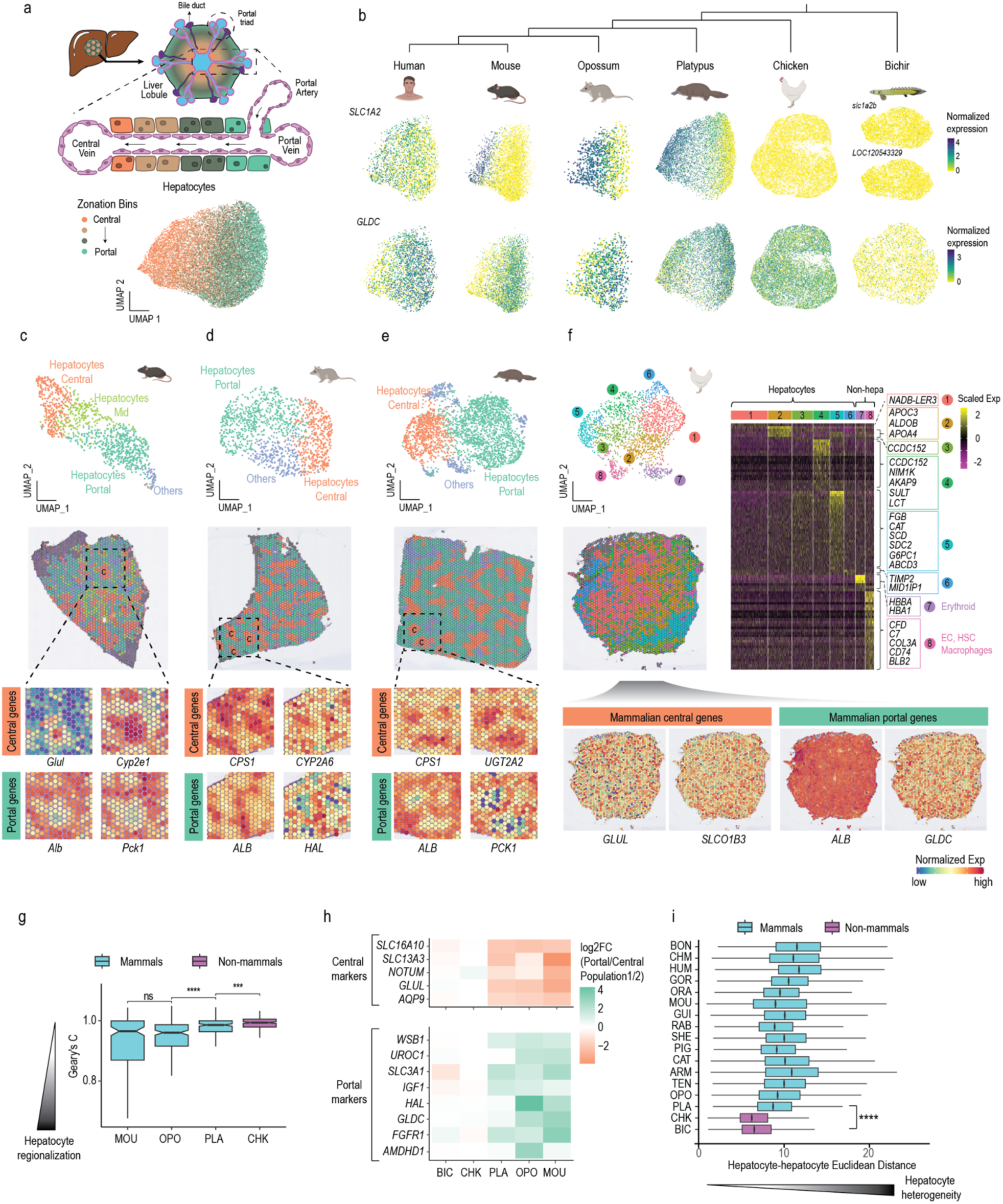
| Spatial organization of the liver in mammalian and non-mammalian species. **a**, Schematic representation of liver architecture (top) showing the liver organized into polygonal units called liver lobules, where blood from the portal vein and hepatic artery flows toward a central vein. UMAP (bottom) of integrated hepatocytes across human, mouse, opossum, and platypus with cells colored by zonation bin (4 bins, from most central to most portal). **b**, Feature plots showing the expression of *SLC1A2* and *GLDC* across hepatocyte species-specific embeddings for human, mouse, opossum, platypus, chicken, and bichir. **c-e**, UMAP (top) and spatial plot (middle) of the Visium data for mouse (**c**), opossum (**d**), platypus (**e**), with zoom-in views of the expression of two central and portal marker genes for each species (bottom). C: Spots represent mostly central hepatocytes. **f**, UMAP (top left), spatial plot (middle left), and heatmap of marker genes (right) of the Visium data for chicken. The expression of mammalian central and portal markers is shown within the spatial plots (bottom). **g**, Distributions of Geary’s C index of spatial autocorrelation for the top 200 highly variable genes in mouse, opossum, platypus and chicken. Pairwise differences were tested with two-sided Mann-Whitney *U* tests; *p*-values were adjusted for multiple comparisons using the Benjamini–Hochberg procedure (****: *P* < 0.0001, ***: *P* < 0.001, ns: not significant). **h**, Log_2_ fold changes of known portal and central hepatocyte marker genes across species. For mammalian species (mouse, opossum, platypus), fold changes represent portal versus central hepatocyte expression. For non-mammalian species (chicken, bichir), fold changes are shown between the two hepatocyte populations that are most different transcriptomically (Methods). **i**, Distribution of pairwise hepatocyte Euclidean distances across species. The closest mammalian and non-mammalian species were compared with a two-sided Mann-Whitney *U* test (****: *P* < 0.0001).

To validate this mammal-chicken difference, we generated spatial transcriptomic data (10x Visium) for chicken and three representative mammals (mouse, opossum, platypus) (Fig. 3c-f and Extended Data Fig. 9). In all three mammals, liver sections showed clusters of cells enriched in central gene expression surrounded by cells expressing portal marker genes, indicating zonation along the porto-central axis (Fig. 3c-e and Extended Data Fig. 9). In chicken, hepatocyte clusters 1–4 were intermixed, and clusters 5–6 were limited to peripheral regions, with no zonal expression of mammalian marker genes, consistent with the snRNA-seq-based results (Fig. 3f). Highly variable genes also showed greater spatial autocorrelation in mammals than chicken (lower Geary’s C values; Mann-Whitney *U* test, *P* < 10^−3^, Fig. 3g), indicating more structured gene expression patterns and regional specialization in mammals. These results demonstrate that chicken hepatocytes lack the zonal organization and transcriptome heterogeneity observed in mammals.

Two scenarios could explain the absence of zonation in chicken: it either emerged in the common mammalian ancestor, or was present in the amniote ancestor but subsequently lost in the avian lineage. To distinguish between these scenarios, we combined our dataset with publicly available snRNA-seq liver data from a fish species, the Senegal bichir^9^. We tested for zonation using two approaches: a targeted analysis of known zonated genes (from mouse and human studies^33,34,39,40^) and an untargeted analysis of hepatocyte heterogeneity (based on pairwise Euclidean distances, Methods). In bichir, as in chicken, we found no evidence of zonated gene expression (Fig. 3h). Hepatocyte heterogeneity was also lower in bichir and chicken than in mammals (Mann-Whitney *U* test, *P* < 10^−4^, Fig. 3i). While heterogeneity alone does not prove zonation, its absence supports a lack of spatial organization in these non-mammalian species.

Together, our findings suggest that zonation is mammal-specific, originating ∼180–320 million years ago in the stem lineage after its split from sauropsids (birds and non-avian reptiles).

## Conservation and innovation of zonated gene expression in mammals

To assess zonated gene expression across mammals, we ordered hepatocytes along the porto-central axis (summarized as a “zonation score”, Methods, Extended Data Fig. 8b and Supplementary Tables 6–21) and identified genes differentially expressed between central and portal hepatocytes in all mammalian species. We found between 224 (gorilla) and 1705 (mouse) zonated genes per species, representing ∼2–12% of hepatocyte-expressed genes (Fig. 4a and Supplementary Tables 22–37).

**Figure 4.**
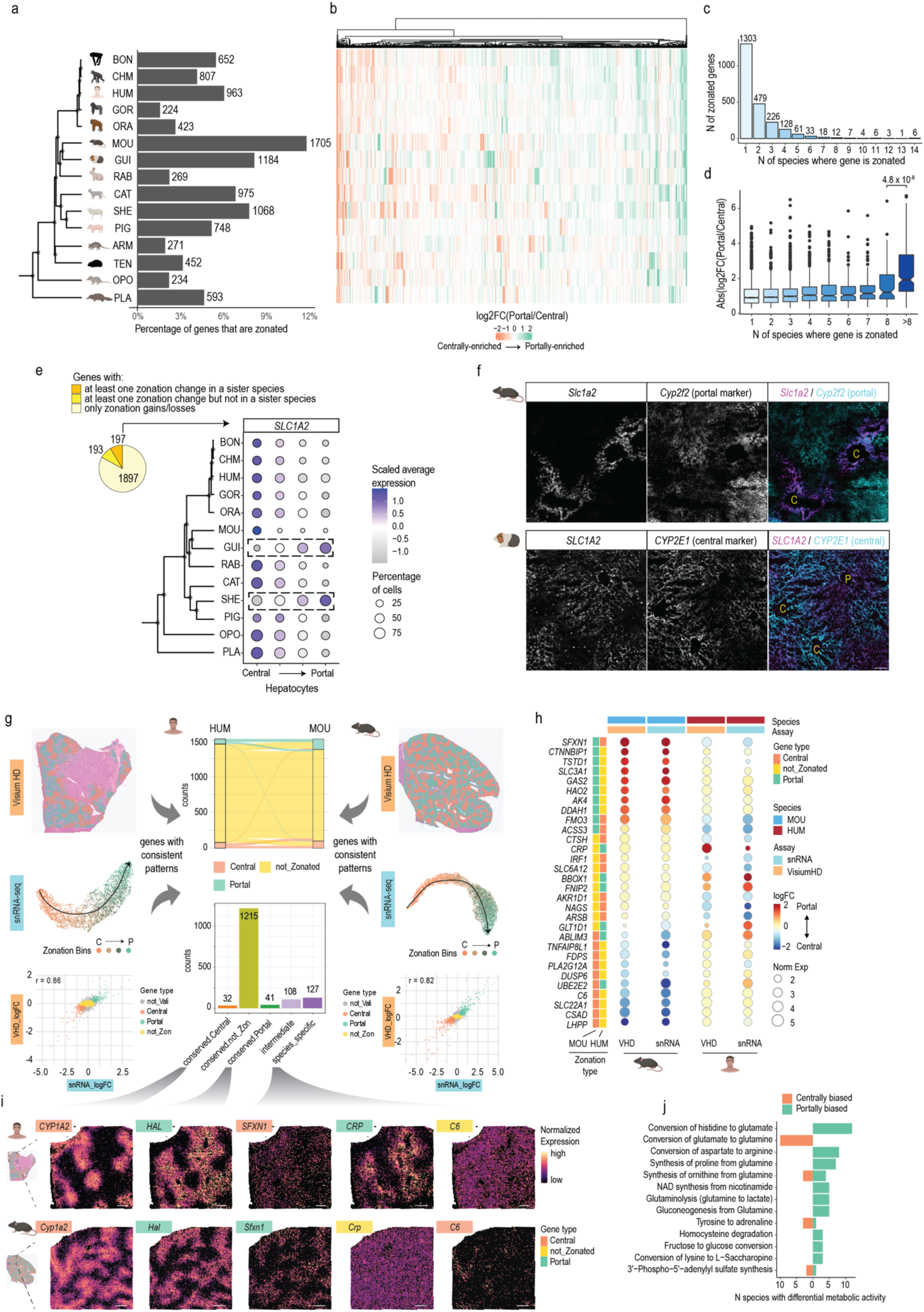
| Conservation and innovation of zonated gene expression in mammals. **a**, Number and percentage of zonated genes per species. **b**, Heatmap of zonated 1:1 orthologs across species. All genes that are zonated in at least 1 species are included (1046 genes). **c**, Barplot classifying genes by the number of species in which they are zonated. **d**, Distribution of log_2_FC(Portal/Central) by degree of conservation (from species-specific to highly-conserved). Statistical significance was assessed with a two-sided Mann-Whitney *U* test (****: *P* < 0.0001). **e**, Genes classified by the class of zonation change they show, and an example of an extreme zonation change (*SLC1A2*). **f**, HCR co-staining of mouse *Slc1a2* with a portal hepatocyte marker *Cyp2f2* on a mouse liver section (top) and HCR co-staining of guinea pig *Slc1a2* with a central hepatocyte marker *Cyp2e1* on a guinea pig liver section (bottom). C: Central vein, P: portal vein. **g**, Visium HD and snRNA-seq data used for calling robust and high-confidence zonation genes in human (left) and mouse (right). Only spots passing the UMI filtering are shown in the Visium HD spatial map, with orange and green denoting central and portal hepatocyte bins, respectively. The correlation between the Visium HD and snRNA-seq data is plotted for each species (bottom left and right). Sankey plot and bar plot showing the number of genes identified for each category (middle). **h**, Dotplot showing the expression of representative species-specific zonation genes in Visium HD and snRNA-seq data in both species. **i**, Spatial feature plots showing the expression of representative genes from different categories. The same areas of the human/mouse sections are selected to show all genes. All scale bars represent 1000 units in the Visium HD coordinate system. **j**, Barplot with the most conserved differentially active metabolic tasks between portal central hepatocytes.

In human and mouse, we detected a smaller fraction of zonated genes (∼6% and ∼12%, respectively) compared with earlier reports (40–50%)^33,34,41^. This difference likely reflects methodological factors: we used a conservative pseudobulk strategy, which better accounts for variations in biological replicates and reduces false discoveries compared to single-cell approaches^42^ (Methods). Variation in hepatocyte capture may also contribute, given that datasets with more captured hepatocytes tend to yield more zonated genes (Extended Data Fig. 8c).

We next traced the genes underlying conserved (ancestral) aspects of liver zonation by comparing the zonation status of expressed 1:1 orthologs across mammals (Methods). Although liver structure and zonation patterns are conserved (Fig. 3a-b), the underlying zonated genes show extensive evolutionary turnover (Fig. 4b). Most zonated 1:1 orthologs are only zonated in one or a few species (Fig. 4c). Genes zonated across multiple (≥9) species show significantly larger fold-change differences between central and portal hepatocytes than less conserved zonated genes (Mann-Whitney *U* test, *P* < 10^−3^, Fig. 4d). The small core of conserved zonated genes (27 genes) is enriched in amino acid metabolism pathways (Extended Data Fig. 8d).

This high turnover largely reflects gains and losses of zonation status (1,897 genes) rather than shifts in zonation direction (portal to central or vice versa; 197 genes, Fig. 4e and Supplementary Table 38). One exception is the glutamate transporter *SLC1A2*, which is centrally expressed in most mammals but independently shifted to portal expression in guinea pig and sheep, as was validated by fluorescence *in situ* hybridization (Fig. 4f).

To further validate the rapid turnover, we performed spatial transcriptomics (Visium HD) on human and mouse livers (Fig. 4g-i and Extended. Data Fig. 10a). Zonation patterns were highly consistent between the snRNA-seq and Visium HD data (Fig. 4g-h). Comparing the two species, we found 127 genes with divergent zonation patterns and 73 with conserved patterns (Methods, Fig. 4i, Extended. Data Fig. 10b and Supplementary Table 39). Most differences reflected gains or losses of zonation, though 17 genes switched directions between human and mouse (examples in Fig. 4h,i). Together, these results demonstrate rapid evolutionary turnover of zonated expression in mammalian livers.

To further investigate zonated metabolic pathways, we used scCellfie^43^ to quantify differential activity between central and portal hepatocytes across species (Methods). This analysis showed strong conservation of amino acid catabolism (e.g., histidine to glutamate, aspartate to arginine) in portal hepatocytes and ammonia detoxification (e.g., glutamate to glutamine) in central hepatocytes (Fig. 4j, Extended Data Fig. 8e). It thus revealed that the previously reported spatial pattern of these opposing pathways^14^ **–** amino acid breakdown generating nitrogenous waste in portal zones, and ammonia clearance via glutamine synthesis in central zones^44^ **–** represent deeply conserved features of mammalian liver zonation.

Taken together, our results show that despite a small conserved core of zonated genes, the overall zonation program evolves rapidly, with frequent gains and losses of the zonation status of genes even over short evolutionary timescales. These expression changes might have contributed to the liver’s adaptive evolution, in addition to selectively-driven coding sequence changes and newly emerged genes (see above).

## WNT and R-spondin secretion by central endothelial cells drives zonation across mammals

Liver zonation is primarily established through interactions between endothelial cells and hepatocytes. In mouse, WNT and R-spondin signaling from the central vein are essential for this process^45,46^, but it is unclear whether this mechanism is conserved across mammals. We thus annotated endothelial cell subtypes along the porto-central axis using marker genes^47–49^ (Extended Data Fig. 11a) and inferred ligand-receptor interactions between endothelial subtypes and hepatocytes in six species (human, mouse, cat, sheep, opossum and platypus) using CellPhoneDB^50^ (Methods and Supplementary Table 40).

In the portal region, Notch signaling is highly conserved (Fig. 5a). JAG1–NOTCH2 surface protein interactions between portal endothelial cells and portal hepatocytes were detected in all six mammals, consistent with their reported essential role in bile duct development in the portal area^51^.

**Figure 5.**
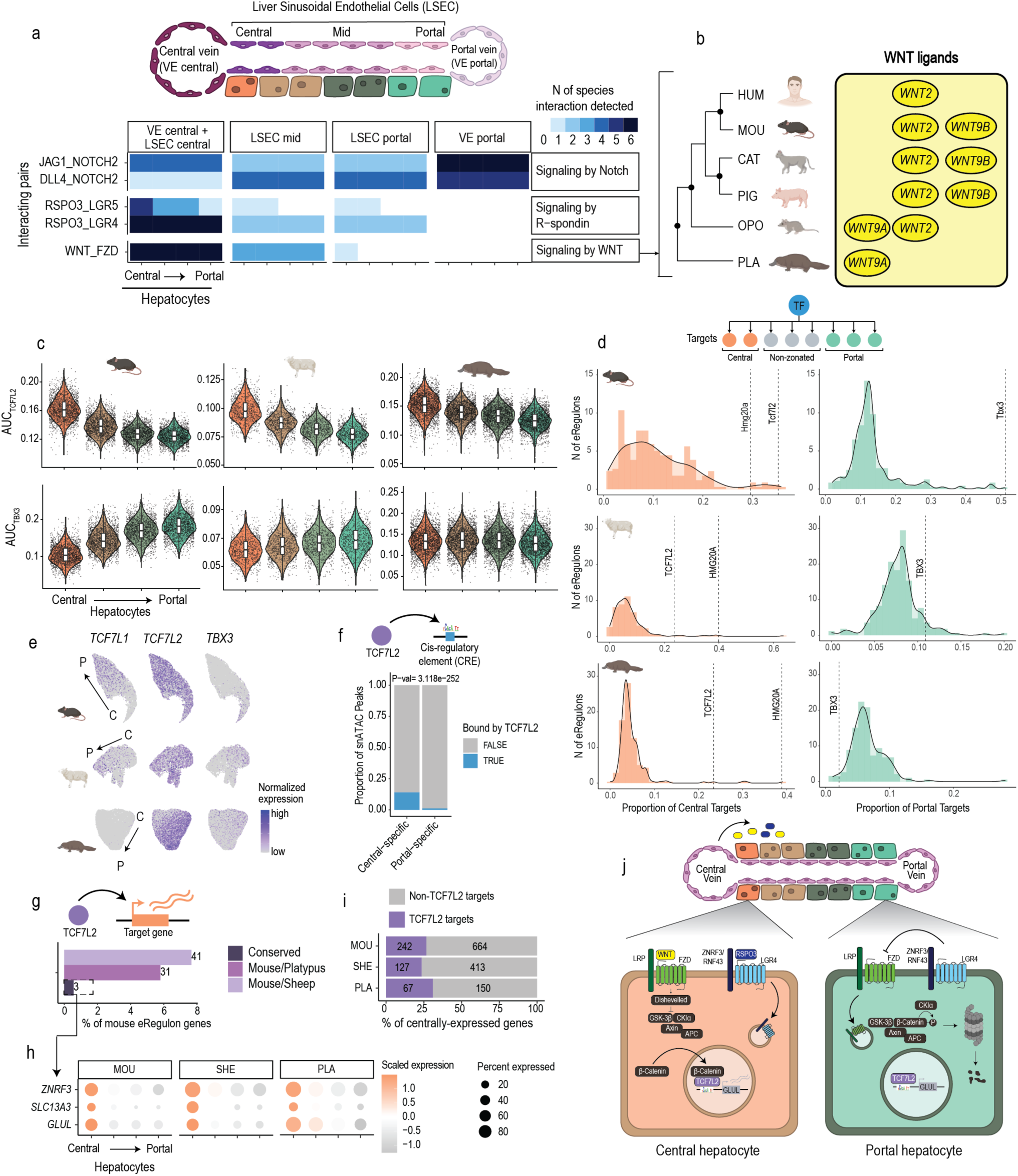
| Molecular mechanisms driving mammalian zonation. **a**, Heatmap with highly conserved ligand receptor interactions between endothelial cells and hepatocytes across human, mouse, cat, pig, opossum and platypus in either the portal or central area of the lobule (VE_central: central vein, LSEC_central: central liver sinusoidal endothelial cells, LSEC_mid: mid-lobular liver sinusoidal endothelial cells, LSEC_portal: portal liver sinusoidal endothelial cells, VE_portal: portal vein). **b**, Schematic representation of the turnover in WNT ligands across different mammalian lineages. **c**, Activity of TCF7L2 and TBX3 eRegulons (as AUC: area under the curve) along the different subtypes of hepatocytes ordered from central to portal in mouse, sheep and platypus. **d**, Histograms and density plots with the proportions of portal and central targets in the eRegulons of all expressed TFs in mouse, sheep and platypus. Relevant TFs are highlighted. **e**, Expression of *TCF7L1*, *TCF7L2,* and *TBX3* across the liver lobule in mouse, sheep, and platypus. **f**, Proportion of central-specific peaks and portal-specific peaks bound by TCF7L2 (based on TCF7L2 ChIPseq data). **g**, Number and percentage of target genes of the *TCF7L2* eRegulon that are conserved across mouse, sheep, and platypus. **h**, Expression profiles of genes that are conserved targets of TCF7L2 (*ZNRF3*, *GLUL*, *SLC13A3*) along the different subtypes of hepatocytes ordered from central to portal in mouse, sheep, and platypus. **i**, Proportion of centrally-expressed genes that are potentially targeted by TCF7L2 in mouse, sheep, and platypus. **j**, Proposed conserved mechanism for establishing central and portal hepatocyte identities across mammals.

In the central region, R-spondin signaling is also shared across mammals (Fig. 5a). RSPO3-LGR4-mediated interaction between endothelial cells and hepatocytes was observed in all six mammals, whereas RSPO3-LGR5-mediated interaction was present in therian (i.e., placental and marsupial) mammals but absent in platypus. *RSPO3* is specifically expressed by central endothelial cells, indicating this morphogen reaches high concentration in the central part of the lobule. *LGR4* is broadly expressed along the lobule, but *LGR5* is specifically expressed in central hepatocytes (Fig. 5a). These patterns suggest that all mammalian hepatocytes are potentially responsive to *RSPO3* through *LGR4*, but that central hepatocytes in therian mammals have enhanced sensitivity due to additional *LGR5* receptors.

WNT signaling is also strongly conserved in mammals but showed turnover in individual genes (Fig. 5b). Previous studies reported conserved *WNT9B* and *WNT2* expression by central endothelial cells in placental mammals^48^. We detected enriched *WNT9B* expression in central endothelial cells in mouse, cat, and pig, but not in human (potentially due to our limited recovery of central vein endothelial cells), or opossum (Extended Data Fig. 11b). Moreover, *WNT9B* has been lost from the platypus genome. We observed conserved expression of *WNT2* in placental mammals as well as in opossum, but no *WNT2* expression was detected in platypus. Instead, platypus central endothelial cells express *WNT9A*, a pattern also observed in opossum (Extended Data Fig. 11b). This suggests that *WNT9A* may have originally driven the establishment of zonation in the ancestor of all mammals, with *WNT2* later taking on this role in therian mammals.

In chicken and bichir, we identified portal and central vein, as well as sinusoidal endothelial cells (Extended Data Fig. 11c-d, 11f-g), confirming the conservation of vascular architecture in the vertebrate liver, as previously reported^17,52^. Unlike mammals, central endothelial cells in these species express almost none of the WNT or RSPO ligands detected in mammals (Methods). This absence is unlikely to reflect a detection artifact, as these genes are well annotated in the respective genomes, and we observed no read coverage beyond annotated 3’ UTRs (Extended Data Fig. 11e, 11h). The only exception is *WNT9A* in bichir, which is sparsely expressed across all endothelial subtypes without central enrichment. In both species, *JAG1* is specifically expressed in portal vein cells, suggesting that Notch signaling between endothelial cells and hepatocytes in the portal region may be conserved across bony vertebrates (Extended Data Fig. 11i).

The conserved endothelial cell expression patterns across mammals, chicken, and bichir suggests that the vascular liver architecture – with the parenchyma perfused by portal and central vessels having distinct properties and transcriptional identities – is conserved across bony vertebrates. However, only mammals have WNT/RSPO morphogen signaling from the central vein, essential for inducing metabolic zonation in hepatocytes^45^, further supporting our finding that liver zonation is a mammalian innovation.

## TCF7L2 regulates central hepatocyte identity across mammals

In mouse, hepatocyte heterogeneity along the lobule has been suggested to be driven by the zone-specific repressors TCF7L1 and TBX3^53^. *TCF7L2*, a paralog of *TCF7L1*, has also been proposed to be essential for maintaining liver zonation in this species^54,55^. In humans, TBX3 and HNF4A have been implicated in regulating zonated gene expression^56^. Whether these mechanisms are conserved across mammals remains unknown.

To identify candidate TFs underlying hepatocyte subtypes, we performed enhancer-driven gene regulatory networks (eGRNs) analyses using SCENIC+^57^ (Methods). We deliberately skipped the last filtering steps of the SCENIC+ pipeline, that retain only target genes that are highly correlated with TF expression (Methods), reasoning that TFs downstream of WNT signaling – though broadly expressed – could still drive zonated expression in response to localized WNT secretion. We then defined an enhancer regulon (eRegulon) as a TF with its predicted target enhancers and genes, requiring correlation between enhancer accessibility and target gene expression, but not between TF expression and target gene expression.

We observed higher TCF7L1/2 eRegulon activity in central than portal hepatocytes in mouse, sheep, and platypus (Fig. 5c). Given that *TCF7L1* is not expressed in platypus, this pattern is likely driven by *TCF7L2* (Fig. 5e). TCF7L2 targeted a higher proportion of centrally expressed genes than other TFs in all three species (ranking above the 90th percentile of the distribution, Fig. 5d), consistent with a role in specifying central hepatocyte identity. HMG20A eRegulon showed a similar pattern (Fig. 5d, Extended Data Fig. 12a); however, strong transcription factor binding site (TFBS) similarity and overlapping eRegulons suggest that this signal may reflect motif similarity with TCF7L2 rather than independent activity (Extended Data Fig. 12b).

We validated the SCENIC+ TCF7L2 eGRN using publicly available mouse liver TF chromatin immunoprecipitation followed by sequencing (ChIP–seq) data^58^. This analysis revealed that TCF7L2 preferentially binds centrally over portally accessible regions (*X*^2^-test, *P* < 10^−251^, Fig. 5f), indicating that TCF7L2 TFBSs present in open chromatin regions correspond to higher TCF7L2 occupancy.

Given TCF7L2’s preference for central genes, we assessed conservation of predicted targets across mouse, sheep, and platypus (Fig. 5g, Supplementary Tables 41–43). Only three targets were conserved, consistent with rapid turnover of zonated expression across mammals (Fig. 3). These three targets are among nine genes showing conserved central expression, including *GLUL*, essential for ammonia-to-glutamine conversion (Fig. 5h), in agreement with mouse knockout studies demonstrating that knocking out TCF7L2 in the mouse liver leads to disorganization of glutamine metabolism^54,55^. Most TCF7L2 targets are, however, species-specific. We found that ∼25% of centrally expressed genes in each species are potentially regulated by TCF7L2 (Fig. 5i).

We found higher TBX3 eRegulon activity in portal hepatocytes compared to central hepatocytes in mouse and – less strongly – in sheep. Although *TBX3* is centrally expressed in all three species (Fig. 5e), it only shows a high proportion of portally expressed targets in mouse and sheep (Fig. 5d), consistent with its proposed role in repressing portal expression in central hepatocytes^53^. Its activity did not vary among hepatocyte subtypes in platypus (Fig. 5c), indicating that this mechanism is not shared with monotremes. No other TF showed consistent targeting of portal genes across all three species (none ranked above the 90th percentile, Fig. 5d).

These results suggest a conserved mechanism for establishing zonation across mammals, mediated by TCF7L2 downstream of WNT to activate central genes in central hepatocytes (Fig. 5j). In species like mouse or human, additional mechanisms, involving TCF7L1 and TBX3 as previously reported^53^, may further refine central and portal hepatocyte identities.

## Molecular changes underlying human-specific metabolic traits

Humans differ from other great apes in several metabolic traits, including higher energy expenditure^59^, greater body fat percentage^59,60^, and unique pathways for metabolizing branched-chain fatty acids^61^. However, the molecular mechanisms underlying these human-specific adaptations remain unknown. We therefore leveraged our comprehensive single-nucleus datasets from great ape livers to gain cellular-level insights into human metabolism. To make use of the cell-type-resolved gene expression data while avoiding issues with absolute quantification in cross-species comparison, we focused on relative gene expression differences at the cell-type level.

We first identified genes with cell-type specific expression changes across liver cell types within the great ape phylogeny (Methods, Extended Data Fig. 13a and Supplementary Tables 44 and 45). Through stringent filtering, we identified 14 genes with human-specific expression shifts (Fig. 6a,b). For example, *CD36* and *ACSM3* are primarily expressed in hepatocytes in bonobo, chimpanzee, and gorilla, but in humans, they are strongly expressed in endothelial cells (Fig. 6b,c). Both genes play essential roles in fatty acid transport and metabolism^62,63^, suggesting that these human-specific cellular expression shifts may contribute to known differences in fat storage and metabolism between humans and other great apes^59–61^.

**Figure 6.**
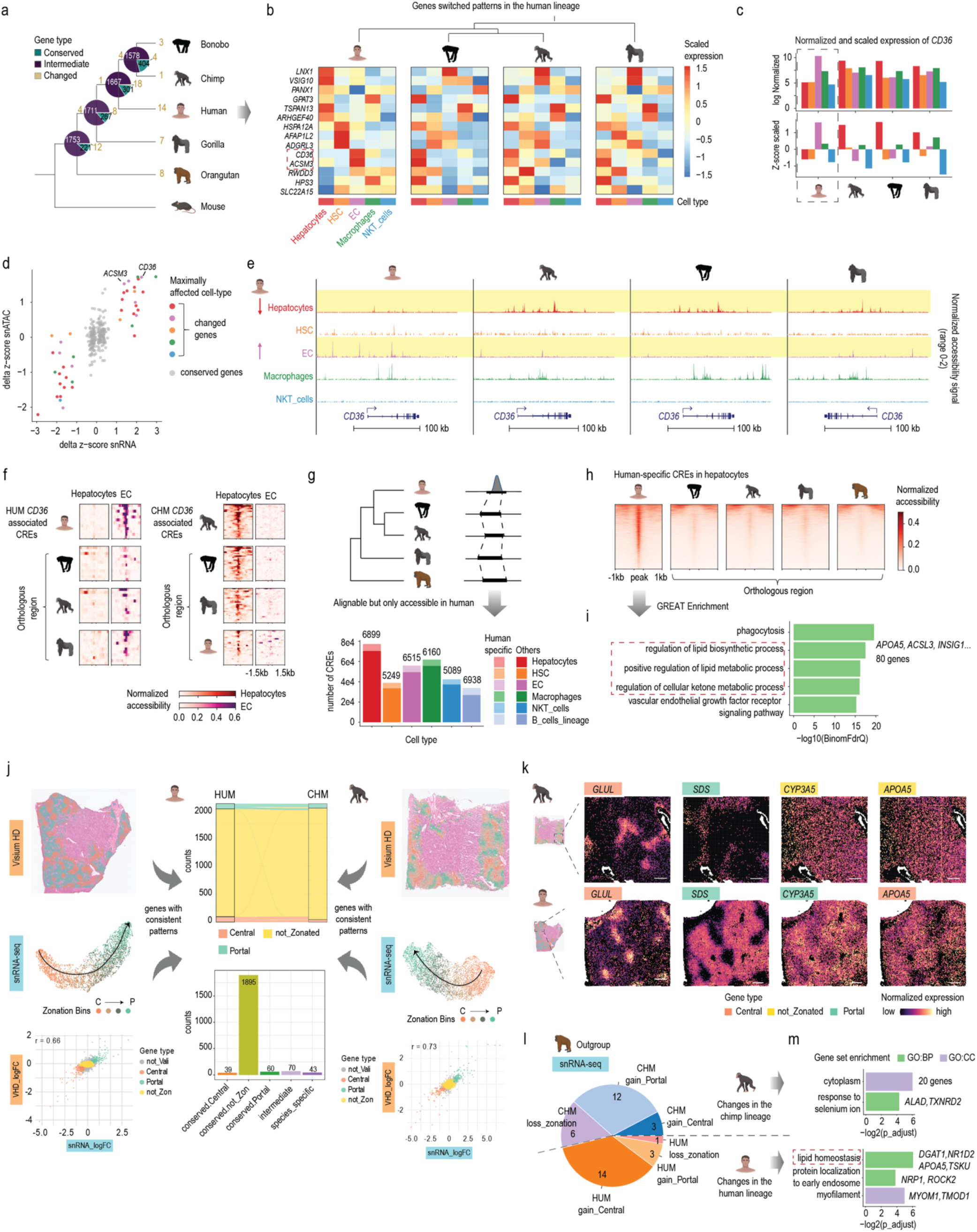
| Human-specific molecular changes in the liver. **a**, Number of genes with conserved, intermediate, and changed cell-type specificity patterns are indicated for each branch within the great ape lineages, with mouse as the outgroup. Branch lengths in the phylogenetic tree do not reflect divergent time. **b**, Heatmap showing the scaled expression (z-score) of the 14 genes that changed relative expression in the human lineage. Red rectangle highlights the genes involved in lipid metabolism. **c**, Dotplot showing the delta z-score (Δ z-score = z-score_changing_node_ – z-score_ancestral_node_) of the conserved (grey) and changed genes (colored by the maximally affected cell types), calculated based on snRNA-seq gene expressions (x axis) or snATAC-seq gene scores (y axis). **d**, Barplot showing the log-transformed, normalized expression (top) and z-score-scaled expression (bottom) of *CD36* in human, bonobo, chimpanzee, and gorilla data. **e**, Chromatin accessibility profiles around the *CD36* locus in each species. All cell-type pseudobulks were processed and normalized following the same pipeline and shown with the same range scale. **f**, Heatmap showing the chromatin accessibility signals centered around *CD36*-associated peaks (1.5 kb extended upstream and downstream) identified in the human liver (left) or in the chimp liver (right) based on correlation, together with the signals from orthologous regions in other great ape species. **g**, Schematics for identifying human-specific CREs and the numbers of human-specific CREs identified in each cell type. **h**, Heatmap showing the chromatin accessibility signals in hepatocytes centered around hepatocyte human-specific CREs (1 kb extended upstream and downstream) and the orthologous regions in other species. **i**, GREAT enrichment analysis of human-specific CREs from hepatocytes. Red rectangle highlights the pathways related to lipid metabolism. **j**, Visium HD and snRNA-seq data for calling robust and high-confidence zonation genes in human (left) and chimpanzee (right). Only spots passing the UMI filtering are shown in the Visium HD spatial map, with orange and green denoting central and portal hepatocytes, respectively. The correlation between the Visium HD and snRNA-seq data is plotted for each species (bottom left and right). Sankey plot and bar plot showing the number of genes identified for each category (middle). **k**, Spatial feature plots showing the expression of representative genes (conserved: *GLUL*, *SDS*; species-specific: *CYP3A5*, *APOA5*) from the human-chimpanzee comparison. The same areas of the Visium HD sections are selected to display all genes. All scale bars represent 1000 units in the Visium HD coordinate system. **m**, Gene set enrichment analysis of genes with changed zonated patterns in the chimpanzee (top) or human (bottom) lineage.

We next examined the regulatory landscapes of genes with lineage-or species-specific cell type switches. Chromatin accessibility-inferred gene scores confirmed most transcriptome-based specificity changes (31/41), indicating that shifts in chromatin accessibility underlie these transcriptional differences (Fig. 6d). The *CD36* locus exemplifies this: in humans, *CD36*-associated CREs are strongly accessible in endothelial cells and weakly in hepatocytes, whereas in bonobo (8/12), chimpanzee (6/12), and gorilla (5/12), orthologous regions remain largely inaccessible in endothelial cells. Instead, *CD36*-associated CREs in the non-human great apes show strong accessibility in hepatocytes, mirroring species-specific expression patterns (Fig. 6e,f).

At a genome-wide scale, we identified human-specific CREs in each major liver cell type. We found 5,089–6,899 CREs with orthologous sequences across all five great apes but accessible only in humans (Methods, Fig. 6g,h and Extended Data Fig. 13b). In hepatocytes, these human-specific CREs are strongly enriched for lipid biosynthesis and metabolic pathways (Fig. 6i), suggesting a potential role in uniquely maintaining human lipid homeostasis.

Zonated gene expression in hepatocytes is essential for metabolic homeostasis^14^. To identify genes with human-specific zonation patterns, we performed Visium-HD experiments on a chimpanzee liver sample and classified the zonation patterns of 3,954 robustly expressed genes (supported by both snRNA-seq and Visium HD data; Fig. 6j, Extended Data Fig. 13c and Supplementary Table 46). We identified 43 genes with zonation differences between human and chimpanzee, mostly representing gains or losses of zonation rather than shifts of direction (Methods, Fig. 6j,k and Extended Data Fig. 13d,e). Among these, 18 genes changed in the human lineage and 21 in chimpanzee (orangutan was used as an outgroup species, Fig. 6l and Extended Data Fig. 13e). The 18 genes that switched zonation patterns in the human lineage are enriched for lipid metabolic processes, whereas chimpanzee-specific genes showed no such enrichment (Fig. 6m). Notably, some human-specific zonated genes (*DGAT1*, *APOA5*) directly mediate triglyceride synthesis or hydrolysis^64,65^, while others (*PDK2*, *ABHD10*) indirectly regulate lipid metabolism or modifications^66,67^.

Together, our genome-wide and candidate-focused analyses highlight molecular and regulatory mechanisms likely underlying known human-specific innovations in lipid metabolism.

## Discussion

In this study, we present a multiomic cellular atlas of mammalian livers spanning representative evolutionary lineages and dietary types, thus capturing transcriptomes, chromatin accessibility, and the spatial organization of liver cell types across mammals. Beyond serving as an extensive resource for studying liver evolution and function, our analyses reveal the evolutionary origins of liver zonation, conserved molecular mechanisms underlying its establishment and maintenance, and the limited conservation of zonation gene modules among the overall fast-evolving zonated genes. Focusing on great apes, we also identified candidate genes and regulatory elements that may contribute to human-specific lipid metabolism.

Across vertebrates, the basic vascular organization of the liver is largely conserved^15–18,68^, and, consistently, we observe similar molecular conservation, as both mammals and other vertebrates exhibit endothelial cell specialization along the portal–central axis. By contrast, our work revealed a fundamental innovation of the organization of hepatocyte gene expression during early mammalian evolution. Mammals evolved well-defined hepatocyte zonation of gene expression across the lobule, whereas in representative bird and fish species, hepatocyte expression is more homogeneous, with no clear zonation, which likely reflects the ancestral architecture.

Hepatocyte spatial specialization in mammals is driven by differential WNT and R-spondin signaling from the central vein. Our results show that this signaling axis is highly conserved, in accord with previous studies highlighting its role in establishing and maintaining zonation^45,46^. Our eGRN analysis indicates that WNT interacts with TCF7L2 (via β-catenin) to activate central gene expression, thereby shaping central hepatocyte identity across mammals. In birds and fish, however, we detect either no expression or no central-enrichment of WNT or R-spondin ligand expression, revealing a fundamental difference in the signaling environment governing liver organization between mammals and other vertebrates.

The evolutionary emergence of this mechanism in mammals may be linked to the central vein’s role as the final exit point for blood flowing through the liver, representing a key metabolic checkpoint before the blood re-enters systemic circulation. It would be advantageous for the central vein to act as a signaling hub, instructing neighbouring hepatocytes to perform a final clearance of potentially harmful metabolites. This concept is well-established for ammonia detoxification^69–72^, where precise spatial compartmentalization ensures blood leaving the liver is low in ammonia. Mammals use two complementary pathways for ammonia clearance: the urea cycle provides a high-capacity, low-sensitivity mechanism detoxifying ∼30% of ammonia^44^ and glutamine synthesis via GLUL offers a high-affinity, low-capacity pathway tightly restricted to central hepatocytes^73,74^. This spatial partitioning positions central hepatocytes as “ammonia-scavengers”, efficiently clearing nitrogenous waste and preventing hyperammonemia^70^. Consistent with this model, many highly conserved zonated genes in our study are involved in amino acid catabolism and glutamine synthesis (Fig. 4j, Extended Data Fig. 8d-e).

The absence of this architecture in non-mammalian vertebrates could reflect different nitrogen excretion strategies. Fish (with few exceptions^75^) excrete ammonia directly via the gills, reducing the need for hepatic detoxification^76^. Amphibians can excrete ammonia directly and/or convert it to urea depending on their lifestyle and access to water^77,78^. Combined with a greater tolerance to ammonia and urea^79^, these mechanisms may reduce the need for specialized hepatic zonation. Birds and reptiles convert ammonia into uric acid for excretion, advantageous for water conservation and adapted to arid environments^80^. In birds, GLUL, an early-acting enzyme in the uric acid synthesis pathway, is both broadly expressed (our data) and highly active across liver tissue^81^, potentially obviating the need for zonated expression. Altogether, our findings support the notion that hepatocyte zonation evolved in mammals as a specialized solution to terrestrial ammonia detoxification.

Our study has limitations. Liver gene expression is influenced by temporal rhythms, hormonal signals, and sex^82–86^, factors we could only partially control. We included both sexes in all species to ensure that major observations were not sex-biased. Another potential confounder is temporal regulation, which is known to affect ∼20% of hepatocyte genes and interact with liver zonation^41^. In the human–mouse zonation comparison, 27 of 127 species-specific genes are also rhythmic in mouse liver^41^, raising the possibility that some differences may reflect temporal states rather than spatial patterning. However, only ∼4% of rhythmic genes alter the slope of zonation gradients over time, with most only oscillating in absolute expression levels rather than spatial distribution^41^. Therefore, we reason that while temporal effects cannot be excluded, their impact is likely limited in the cross-species zonation comparison. Consistently, our biological replicates cluster strongly by species, indicating interspecies differences dominate over sampling variation.

Leveraging dedicated single-nucleus and spatial transcriptomics data, we identified human-specific shifts in zonation patterns and cell-type specificity, undetectable by bulk transcriptomics. For example, one candidate gene, *CD36*, plays complex, cell-type-dependent roles in lipid uptake and lipogenesis^87,88^, and has been strongly implicated in the development of fatty liver diseases^89^. In mice, hepatocyte-specific *Cd36* deletion protects against diet-induced fatty liver^87^, whereas endothelial-specific deletion increases liver triglycerides^90^. We identified a human-specific shift in *CD36* expression from hepatocytes to endothelial cells after the human–chimpanzee split. This shift, which reduced *CD36* expression in hepatocytes and elevated it in endothelial cells, may limit hepatic lipid uptake while enabling non-harmful fat accumulation elsewhere in the body. Such molecular changes align with the hypothesis that the emergence of efficient fat storage conferred survival advantages during human evolution^91^.

## Methods

### Data reporting

No statistical methods were used to predetermine sample size. The experiments were not randomized and investigators were not blinded to allocation during experiments and outcome assessment.

### Sample collection and ethics statements

We used flash frozen samples from the following species for generating all data in this study: from human (Homo sapiens; abbreviation: HUM), chimpanzee (*Pan troglodytes*; abbreviation: CHM), bonobo (*Pan paniscus*; abbreviation: BON), gorilla (*Gorilla gorilla*; abbreviation: GOR), orangutan (*Pongo abelii*; abbreviation: ORA), rabbit (*Oryctolagus cuniculus*; abbreviation: RAB), mouse (*Mus musculus*; abbreviation: MOU), guinea pig (*Cavia porcellus*; abbreviation: GUI), cat (*Felis catus*; abbreviation: CAT), dog (*Canis lupus familiaris*; abbreviation: DOG), sheep (*Ovis aries*; abbreviation: SHE), pig (*Sus scrofa*; abbreviation: PIG), nine-banded armadillo (*Dasypus novemcinctus*; abbreviation: ARM), lesser hedgehog tenrec (*Echinops telfairi*; abbreviation: TEN), grey short-tailed opossum (*Monodelphis domestica*; abbreviation: OPO), platypus (*Ornithorhynchus anatinus*; abbreviation: PLA) and chicken (*Gallus gallus*; abbreviation: CHK) (Supplementary Table 1).

Our study complies with both local and international ethical regulations regarding samples from human and other species. Human liver samples were obtained from the Tissue Bank for Developmental Disorders at the University of Maryland (USA). Human post-mortem frozen tissue samples of the prefrontal cortex from healthy individuals were provided by the Human Brain Tissue Bank at the Semmelweis University. Informed consent for the use of tissues for research was obtained in writing from donors or their families. All primates suffered from deaths for reasons other than their participation in this study and without any relation to the organ sampled. The lesser hedgehog tenrec samples were collected under the experimentation permit GE/82/14 issued by the Geneva cantonal veterinary authority. All sample sources are provided in detail in Supplementary Table 1. The use of all samples for this type of work described in this study was approved by an ERC Ethics Screening panel (associated with H.K.’s ERC Consolidator Grant 615253, OntoTransEvol), Swiss National Science Foundation (associated with Sinergia grant 189970, joint with Clauss, Salzburger and Tschopp labs in Basel and Zürich), local ethics committees in Lausanne (authorization 504/12), Heidelberg (authorization S-220/2017), and Semmelweis University (No.32/1992/TUKEB).

### Sample quality control and nuclei prep

#### Liver

Liver samples were cut into pieces of suitable sizes (5–10 mg for nuclei preps, 2–5 mg for RNA extractions) on a pre-cleaned, pre-cooled metal stage (wrapped in aluminum foil) that was placed on dry ice. Everything used for cutting (forceps, razor, tubes, etc.) was also pre-cooled on dry ice so the tissue blocks remained frozen during the whole cutting process. For quality control, RNA was extracted directly from tissue pieces using RNeasy Micro kits (Qiagen) following manufacturer’s protocol. RNA quality was assessed on the Fragment Analyzer (Agilent). All extracted RNAs had an RQN value above 7.6 except for one orangutan sample (Supplementary Table 1).

Liver nuclei were extracted following a published protocol^92^ with small modifications. We started with 5–10 mg of frozen liver samples and homogenized using a micropestle in 400 μl ice-cold homogenization buffer. The homogenization buffer always contained 250 mM sucrose, 25 mM KCl, 5 mM MgCl2, 10 mM Tris-HCl (pH 8), 0.1% IGEPAL, and 1 μM DTT. For separate snRNA-seq and snATAC-seq experiments, we added 0.4 U/μl Murine RNase Inhibitor (New England BioLabs, cat# M0314L), 0.2 U/μl SUPERas-In (Ambion, cat# AM2694), and cOmplete Protease Inhibitor Cocktail (Roche, cat# 11 836 145 001). For single-nucleus Multiome experiments, we added the 1U/μl Sigma Protector RNase inhibitor (cat# 3335402001) recommended by 10X Genomics. To homogenize the samples, we triturated gently with a P1000 tip for 10 times, incubated on ice for 5 minutes, and then centrifuged at 100g for 1 minute at 4 °C to pellet any unlysed tissue chunks. The supernatant was transferred into another 1.5 mL Eppendorf tube and centrifuged at 400g for 4 minutes at 4 °C to collect nuclei. The nuclei were washed twice in 400 μl homogenization buffer and strained by a 40 μm Flowmi strainer (Sigma, BAH136800040) during the second wash step to remove nuclear aggregates. We always resuspended the final nuclei pellet in 20–50 μl nuclei buffer (10X Genomics, PN-2000207) but added different inhibitors to the nuclei buffer depending on the experiments (multiome: 1mM DTT and 1U/μl Sigma Protector RNase inhibitor; snRNA-seq: 0.2 U/μl SUPERas-In (Ambion) and 0.4 U/μl Murine RNase Inhibitor (New England BioLabs)). To estimate the nucleus concentration, nucleus aliquots were diluted in PBS with Hoechst and PI DNA dyes and counted on Countess II FL Automated Cell Counter (Thermo Fisher Scientific). 15,000–20,000 nuclei were loaded into the 10X Chromium machine as input for each experiment.

#### Cortex

Human cortex nuclei were prepared following a similar workflow with the following modifications: (1) The volume of homogenization buffer was adjusted to match the size of the tissue. (2) The supernatant after the first centrifuge (100g, 1min), which contained rich cytoplasmic fractions, was collected for quality control later. (3) The nuclei pellet was only washed once and resuspended in 1xPBS with Hoechst, 0.4 U/μl Murine RNase Inhibitor (New England BioLabs, cat# M0314L), and 0.2 U/μl SUPERas-In (Ambion, cat# AM2694) to reach a concentration of 2–10 x 107 cells/ml. (4) High-purity single nuclei were further isolated by flow cytometry using BD FACS Aria III U with a 70 µm nozzle. Nuclei were gated by their size and scatter properties and secondarily by their Hoechst signals. Doublet discrimination gates were used to exclude nuclear aggregates. Hoechst signal was detected using a 407 nm laser. Usually, 60,000–100,000 events were collected, and concentration was determined after sorting using the Countess II FL Automated Cell Counter (Thermo Fisher Scientific).

### Bulk RNA-sequencing

We have generated bulk liver RNA-seq data for the following species: guinea pig, gorilla, bonobo, chimpanzee, orangutan, cat, dog, pig, sheep, and armadillo (Supplementary Table 2). RNAs were extracted directly from tissue pieces using RNeasy Micro or Mini kits (Qiagen). All extracted RNAs had an RQN value above 8.1. Bulk RNA-sequencing libraries were prepared with the NEBNext Ultra II RNA kit (New England Biolabs) at the Deep Sequencing Core Facility of Heidelberg University. Qubit Fluorometer (Thermo Fisher Scientific) was used to estimate DNA concentrations, and the average fragment size was determined on a Bioanalyzer 2100 (Agilent). Libraries were sequenced on Illumina NextSeq 550 using High Output Kit v2.5 with 150 cycles (Illumina) and the following setting: 159 cycles for Read 1 (cDNA), 8 cycles for i7 index (sample index), to an average depth of 30 million reads per library.

### Single-nucleus library generation and sequencing

The transcriptome and accessibility data for platypus and chicken were jointly profiled from the same nuclei using Chromium Next GEM Single Cell Multiome ATAC + Gene Expression Reagent kit (10X Genomics, PN-1000283). All the other snRNA-seq datasets produced in this study were generated using the Chromium Next GEM Single Cell 3ʹ Reagent Kits v3.1 (10X Genomics, PN-1000121). The other snATAC-seq datasets were generated using the Chromium Next GEM Single Cell ATAC kit v1.1 (10X Genomics, PN-1000175). We always generated the separately-profiled snRNA-seq and snATAC-seq datasets from the same nuclei preps for the same biological replicates when possible (Supplementary Table 1). All libraries were prepared following manufacturer’s instructions. Library concentration was quantified on a Qubit Fluorometer (Thermo Fisher Scientific) and the average fragment size was determined on a Fragment Analyzer (Agilent).

Multiome libraries were sequenced on NextSeq 500/550 using the High Output Kit v2.5 with 150 cycles (Illumina) or on NextSeq2000 P2 flowcell with 2 x 50 bp reagent kit (Illumina). The following setting was used for Multiome snRNA libraries: 28 cycles for Read 1 (cell barcode), 10 cycles for both i7 and i5 indices (sample index), 90 cycles for Read 2 (cDNA). The following setting was used for Multiome snATAC-seq libraries: 50 cycles for both Read 1 and 2 (gDNA), 8 cycles for i7 index (sample index), 16 cycles for i5 index (cell barcode). A custom recipe that includes 8 dark cycles on i5 was provided by Illumina for sequencing multiome snATAC-seq libraries on NextSeq 500/550.

Separately profiled snRNA-seq and snATAC-seq libraries were sequenced on NextSeq 500/550 (Illumina) using the High Output Kit v2.5 with 75 Cycles (Illumina). The following setting was used for separately profiled snRNA-seq libraries: 28 cycles for Read 1 (cell barcode), 8 cycles for i7 index (sample index), 56 cycles for Read 2 (cDNA). The following setting was used for separately profiled snATAC-seq libraries: 34 cycles for both Read 1 and 2 (gDNA), 8 cycles for i7 index (sample index), 16 cycles for i5 index (cell barcode).

snRNA-seq libraries, including those generated in the Multiome experiments, were sequenced to a depth of 200 – 540 million reads (usually in multiple batches of sequencing), depending on the number of nuclei captured in the libraries (Supplementary Table 3). We experienced a wider range of nucleus numbers (253 – 13173) captured in different snATAC-seq experiments, so the libraries were sequenced to a wider range of depths (70 – 600 million reads) accordingly (Supplementary Table 4).

Our mouse liver snRNA-seq data, generated with the same protocol, were previously published in Rodríguez-Montes et al^85^. For cat, we obtained the data for another cat sample from Chen et al^93^, which corresponded to CAT_F2 in our figures and tables, besides the two cat snRNA-seq datasets that we generated ourselves (CAT_F1 and CAT_M1). We downloaded the fastq raw sequencing files of the published data and followed the same analysis pipeline as described below.

### Cryosection

We used cryosectioning to prepare samples for regular Visium (bin size: 50 μm), Visum HD (smallest bin size: 2 μm), and HCR RNA-FISH. For all these experiments, flash frozen samples were embedded in optimal cutting temperature (OCT) mounting medium on dry ice and cryosectioned into 10 μm sections (environmental temperature: between –21 and –19 °C; sample temperature: between –20 and –18 °C). For Visium HD and HCR experiments, sections were collected on Epredia Superfrost Plus Adhesion Slides (Thermo Fisher Scientific 10149870). For the regular Visium experiment, sections were collected on Visium Spatial Gene Expression slides (10X Genomics). All slides were stored in sealed containers at –70 °C according to manufacturer’s recommendation until further processing.

### Spatial transcriptome library generation and sequencing

As a validation experiment, the regular Visium spatial transcriptomic experiment were performed on one sample from mouse, opossum, platypus, and chicken (Supplementary Table 1) with the Visium Spatial Gene Expression Slide & Reagent Kit (10X Genomics, PN-1000184). Optimized permeabilization time for optimal tissue digestion and RNA release was determined with Visium Spatial Tissue Optimization Kit (10X Genomics, CG000238_RevF). The permeabilization time for platypus liver sections was 12 minutes; for opossum, chicken, and mouse, the permeabilization time was determined to be 24 minutes. Liver tissue sections were placed directly onto the capture areas of Visium Spatial Gene Expression slides and kept at –70°C until processing. Libraries were constructed following the manufacturer’s protocol (10X Genomics, CG000239_RevF). Briefly, sections melted onto the Visium slide were fixed in chilled Methanol, stained with Hematoxylin and Eosin, then imaged with an Olympus VS200 slide scanner (20X magnification). Tissues were permeabilized, the RNA was captured in situ, reverse-transcribed, and amplified. Barcoded libraries were pooled and sequenced with an Illumina NextSeq 2000 P3 kit with the following setting: 28 cycles for Read 1 (spatial barcode and UMI), 10 cycles for both i7 and i5 indices (sample index), 90 cycles for Read 2 (cDNA), to reach a depth of 210 – 490 million reads per library.

The mouse Visium HD spatial data were generated from two mouse samples with the Visium HD Mouse Transcriptome kit (10X Genomics, PN-1000676), while the human and chimpanzee Visium HD spatial data were generated from one sample from each species with the Visium HD Human Transcriptome kit (10X Genomics, PN-1000675). These Visium HD kits utilize probe-based approaches for catching mRNA. The high protein-coding sequence identity (99.1%) between human and chimpanzee^94^ allows the robust capture of chimpanzee mRNA with the same set of human probes. We collected the sections on Epredia Superfrost Plus Adhesion Slides (Thermo Fisher Scientific 10149870), which were compatible according to Visium HD Fresh Frozen Tissue Preparation Handbook (10X Genomics, CG000763_RevB). Sections were paraformaldehyde fixed and stained with Hematoxylin and Eosin. Imaging of sections was carried out with an Olympus VS200 slide scanner (20X magnification). Probe hybridization, ligation, and probe transfer to Visium HD capture slide with CytAssist was performed according to the Visium HD Spatial Gene Expression Reagent Kits User Guide (CG000685_RevB). The constructed Visium HD libraries were pooled and sequenced with an Illumina NextSeq 2000 P3 kit with the following setting: 43 cycles for Read 1 (UMI and spatial barcode), 10 cycles for both i7 and i5 indices (sample index), 50 cycles for Read 2 (probe insert), to reach a depth of 610 – 680 million reads per library.

### HCR RNA-FISH

HCR RNA-FISH reagents, including target-specific probe sets, amplifiers, and buffers, were ordered from Molecular Instruments. Frozen tissue sections were prepared as described in *Cryosection*. The sections were fixed with 4% PFA for 15 min at room temperature before being dehydrated in 70% EtOH overnight at 4 °C. Then sections were processed according to manufacturer’s protocols (MI Protocol-RNAFISH-FreshFixedFrozenTissue, Revision Number 4) with optimized probe concentration (mouse *Slc1a2*: 2 pmol per 100 μl, mouse *cyp2f2*: 0.4 pmol per 100 μl, guinea pig *SLC1A2*: 2 pmol per 100 μl, guinea pig *CYP2E1*: 2 pmol per 100 μl). Briefly, sections were incubated in 200 μl pre-hybridization buffers at 37 °C for 15 minutes within a humidified chamber. Then sections were incubated with probe sets of optimized concentration at 37 °C in a humidified chamber overnight. Coverslips were gently placed on top of the tissues to prevent evaporation. On the second day, consecutive washes at 37 °C with probe wash buffers and 5xSSCT were conducted to remove excess probes before preamplification at room temperature. Hairpin solutions were prepared by snap-cooling and added on top of the tissues for overnight incubation at room temperature. Sections were mounted within ProLong Diamond Antifade Mountant with DAPI (Thermo Fisher Scientific P36962) and imaged on an Olympus IX81 CellSens microscope. Staining for two biological replicates was conducted for both species.

### Genome and transcriptome annotation

#### For liver datasets

We downloaded the reference genomes from Ensembl release 103 for the following species: rabbit (OryCun2.0, GCA_000003625.1), guinea pig (Cavpor3.0, GCA_000151735.1), sheep (Oar_rambouillet_v1.0, GCA_002742125.1), pig (Sscrofa11.1, GCA_000003025.6); from Ensembl release 104 for the following species: human (GRCh38.p13, GCA_000001405.28), chimpanzee (Pan_tro_3.0, GCA_000001515.5), bonobo (panpan1.1, GCA_000258655.2), gorilla (gorGor4, GCA_000151905.3), mouse (GRCm39, GCA_000001635.9), cat (Felis_catus_9.0, GCA_000181335.4), opossum (ASM229v1, GCA_000002295.1); from Ensembl release 106 for the following species: orangutan (Susie_PABv2, GCA_002880775.3), dog (ROS_Cfam_1.0, GCA_014441545.1), armadillo (Dasnov3.0, GCA_000208655.2); from Ensembl release 110 for the following species: platypus (mOrnAna1.p.v1, GCA_004115215.2) and chicken (bGalGal1.mat.broiler.GRCg7b, GCA_016699485.1). For tenrec, we downloaded the ASM31398v2 (GCF_000313985.2) assembly from NCBI.

Since chromosomes 1 and 2 in the opossum genome are giant, which causes issues in many bioinformatic tools, we split both of them at position 530130000 with no gene annotation disrupted. The second segments of chromosomes 1 and 2 were named 1b and 2b, respectively. We used the genome and transcriptome after splitting for building references and carrying out most analyses. We restored the coordinates in chromosome 1b and 2b back to the original ones for evolutionary analyses in Extended Data Fig. 7.

We used the Ensembl annotation for human and mouse, the two species with the most extensive research and annotation. For almost all other species, the Ensembl annotation was extended using our previously published liver RNA-seq data^8,95^ or newly generated bulk RNA-seq data based on a previously established pipeline^96^ (Supplementary Table 2). Bulk RNA-seq reads were trimmed with Trimmomatic^97^ (v0.39) (LEADING:20 TRAILING:20 SLIDINGWINDOW:5:25 MINLEN:36) and aligned to the respective genome and transcriptome with STAR^98^ (v2.7.9a). Bam files of biological replicates were merged after alignment. We then assembled models of transcripts expressed in each tissue using StringTie (v2.2.1)^99^ (parameters: –f 0.2 –m 200 –a 10 –j 3 –c 2.5 –g 10 –M 0.5). We compared the assembled transcript models to the corresponding reference Ensembl annotations using the cuffcompare program from the cufflinks package (v2.2.1)^100^. We then combined the newly identified transcripts with the respective Ensembl gene annotation into a single gtf file.

Manual inspection of the platypus gtf file (after bulk RNA-seq-based extension) revealed poor annotation of some WNT ligands. Specifically, we observed extended transcript coverage beyond the annotated 3′UTRs of the corresponding genes, which suggests that the reads mapping to those regions were not being assigned to the corresponding genes. To ensure accurate quantification of these genes, we further extended the platypus gene annotation file using GeneExt^101^, with the following parameters: –-clip_5prime –m 5000 –j 30.

We generated Cellranger references, STARsolo^98^ (v2.7.9a) references, and ArchR^102^ (v1.0.2) annotation for each species based on the genome assemblies and custom annotations described above. For ArchR annotation, we only considered protein-coding genes, as we observed that the inclusion of non-coding genes led to noisy proximity-based gene score estimates.

#### For human cortex data

We used the genome assembly GRCh38 and another custom transcriptome annotation. This custom annotation was based on the transcriptome reference from Ensembl release 87, extended with human brain RNA-seq data^8,95^ using the same pipeline^96^.

### Processing and quality control of snRNA-seq data

We used STARsolo^98^ (v.2.7.9.a) aligner to map the raw reads to reference genomes and transcriptomes (--soloType CB_UMI_Simple clipAdapterType CellRanger4; –-outFilterScoreMin 20; –-soloCBmatchWLtype 1MM_multi_Nbase_pseudocounts; –-soloUMIfiltering MultiGeneUMI_CR; –-soloUMIdedup 1MM_CR; –-soloMultiMappers EM). We distributed multi-mapping reads based on the expectation-maximisation algorithm (--soloMultiMappers EM) and counted reads in two modes: exon-only reads and reads that mapped to the full transcripts. We calculated the fraction of intronic reads based on the read count from these two modes. Akin to previous studies^103,104^, we used both total UMI counts (knee-point) and the fraction of intronic reads to distinguish nucleus-containing barcodes from the empty barcodes, based on the idea that nuclear RNA contains a much larger fraction of pre-mRNA (with introns) than ambient RNA from the cytoplasm. We used scrublet^105^ (v0.2.3) to calculate doublet scores and imported the scores into Seurat^106^ (v4) for further examination. To remove low-quality and doublet barcodes, we first filtered out outlier barcodes on the basis of very low or very high UMI counts or high mitochondrial reads. We then applied an initial round of clustering (approaches described in detail in *Dimensional reduction, clustering, and cell type annotation*) on the dataset using Seurat^106^ (v4) and examined each cluster carefully. Low-quality clusters (low UMI counts, low number of genes detected, high mitochondrial reads) without unique marker genes or clusters high in doublets (high doublet scores, detection of mutually exclusive cell-type markers) were removed.

### Processing and quality control of snATAC-seq data

Cellranger-atac (v2.0.0, 10x Genomics) was used with default settings for demultiplexing and aligning reads to the reference genomes. We then calculated the fraction of reads in promoters and fraction of reads in peaks based on the output from cellranger-atac count. We used these metrics together with the number of fragments detected to identify nuclei-containing barcodes from the empty barcodes, keeping only barcodes that showed high values in all three metrics for downstream analysis in ArchR (v1.0.2)^102^. Doublet scores were estimated based on in-silico doublet simulation in ArchR and a filtering ratio of 1 was applied in the initial double removal (nuclei with the top N doublet scores were removed. N = filterRatio* cellnumber^2 / (100000)). Then, we applied an initial round of clustering (approaches described in detail in *Dimensional reduction, clustering, and cell type annotation*) on the datasets. Low-quality clusters (low number of fragments, low TSS enrichment, high mitochondrial reads) without clear unique marker gene signals or clusters high in doublets (high doublet scores, detection of mutually exclusive cell-type markers) were removed.

### Processing and quality control of snMultiome data

We processed the sequencing data and identified nuclei-containing barcodes from the two modalities (snRNA-seq and snATAC-seq) separately. The demultiplexing and genomic mapping of the snATAC-seq modality were carried out with cellranger-arc (v2.0.0, 10x Genomics) with default settings. The demultiplexing and mapping of the snRNA-seq modality were performed with STARsolo^98^ (v.2.7.9.a) as described above in the *Processing and quality control of snRNA-seq data* section. Then we conducted quality control on the RNA-seq and ATAC-seq modalities separately, following the same approaches as described in *Processing and quality control of snRNA-seq/snATAC-seq data* sections. Overall, the RNA-seq quality correlates well with the ATAC-seq quality. We kept only barcodes that passed quality control in both modalities for most analyses, except for the transcriptome-only analysis in platypus, for which we included all nuclei passing quality control of the RNA modality to retain enough nuclei to robustly resolve subtypes of endothelial cells (e.g., central veins, liver sinusoid endothelial cells, portal veins).

### Dimensional reduction, clustering and cell type annotation

#### snRNA-seq data modality

After filtering for the nucleus-containing barcodes from STARsolo raw outputs, we imported the filtered count matrices (from full-transcript counting mode) into Seurat^106^ (v4). Each library was individually normalized with the *SCTransform* function. Variances from percent of mitochondrial reads and UMI numbers were regressed out during the normalization, and the top 3000 variable genes were identified. We applied PCA with *RunPCA* and determined the appropriate number of principal components via the elbow plots. We identified cell clusters using the *FindNeighbors* and *FindClusters* functions with a resolution between 0.5 – 1. This initial clustering was aimed at identifying doublet and low-quality cell groups (as described in *Processing and quality control of snRNA-seq data* section).

After finishing filtering on each library, we integrated the processed data from the same species using the CCA approach provided by Seurat^106^ (v4). Dimensional reduction (*RunPCA* and *RunUMAP*) and clustering (*FindNeighbors* and *FindClusters*) were performed again on the integrated data. Liver cell types were annotated based on the following marker genes: peri-central hepatocytes (*SLC1A2, GLUL, LGR5, CYP2A5, CYP2E1, RGN, HRG, SERPIN2, SLCO1B3, SULT2A8, ABCC2*), peri-portal hepatocytes (*HAL, SLC7A2, GLDC, GLS2, PCK1, NRG4, ASS1, DPYD, HSD17B13, AOX3, AHSG*), endothelial cells (*LDB2*, *PLPP1, FYN, STAB2, PTPRB, FGD5*), hepatic stellate cells (*NRXN1, RELN, ANK3, GPC6, SOX5, COLEC10, VIPR1, CCL21, SEMA3A*), cholangiocytes (*PKHD1, BICC1, GPM6A, CDON, SEMA3C, DCDC2, CHST9, WNK2*), general macrophages (*FYB, CD163, HDAC9, SLC8A1, MARCO, MYO9A*), activated macrophages (*LYZ, VCAN, FCN1, CD74, S100A6, FLT3, PTPRE*), B cells (*PAX5, EBF1, BANK1, BACH2, MEF2C*), plasma cells (*IRF4, PIM2, FCRL5, RALGPS2, CREB3L2, TXNDC11, XBP1*), T cells (*SKAP1, CAMK4, DOCK2, RUNX1, TNIK*), NK cells (*CD7, CMC1, RUNX2, KLRF1,*

*IL2RB, TXK*). These marker genes were extracted from previous snRNA-seq or single-cell RNA-seq of human and mouse livers^34,39,107^. In different species, we commonly observed combinatory expression of different subsets of the marker genes for a given cell-type in different species and annotate according. This annotation is reported as the “uni_broad.CellTypes” in the shiny app and the Seurat objects provided. A more general annotation (hepatocytes, endothelial cells, hepatic stellate cells, cholangiocytes, macrophages, B_cell_lineage, NKT_cells) is also reported as the “general.CellTypes” in the shiny app and the Seurat objects.

#### snATAC-seq data modality

Following quality control and doublet removal, we imported the filtered fragment data into ArchR^102^ for downstream analysis. We used ArchR to summarize the chromatin accessibility profiles within tiled windows of 500 bp across the whole genome and perform dimensional reduction using iterative Latent Semantic Indexing (LSI) algorithms (*addIterativeLSI*). We inspected the correlation between sequencing depth and the first dimension and removed the first dimension from downstream analysis if it was strongly correlated with sequencing depth. Samples from the same species were further integrated with Harmony^108^ to remove batch effect and facilitate clustering (*addHarmony*). We estimated gene scores based on the overall accessibility signals across its regulatory domains and identified the marker genes (genes with enriched gene scores) for each cell cluster. Following the initial clustering, low-quality clusters (low number of fragments, low TSS enrichment, lack of marker genes) or doublet clusters (high doublet scores, detection of mutually exclusive cell-type markers) were removed from further analysis. After filtering, we reran the dimensional reduction (*addIterativeLSI*), integration (*addHarmony*), and clustering (*addClusters*) pipeline and proceeded with cell type annotation.

For species for which transcriptome and chromatin accessibility datasets were generated separately, we integrated snRNA-seq and snATAC-seq data by comparing gene expression (snRNA-seq) with gene scores (snATAC-seq).

We used the *addGeneIntegrationMatrix* function to find the nuclei in the snRNA-seq data most similar to the nuclei in the snATAC-seq data and assigned the cell type label and gene expression of the snRNA-seq cell to the snATAC-seq cell. Integration showed high confidence scores at the general cell type level (atac_general_CellType), allowing us to primarily transfer cell type labels from snRNA-seq to annotate snATAC-seq datasets. For the few snATAC-seq clusters with lower confidence scores, we manually inspected and annotated them based on gene score-derived marker genes. For species where transcriptome and chromatin accessibility were jointly profiled from the same nuclei, we directly assigned cell type labels to the snATAC-seq modality using the corresponding snRNA-seq annotations based on cell barcode identity.

### Identification of putative CREs

We followed the ArchR (v1.0.2)^102^ pipeline for calling regions with enriched chromatin accessibility signals, i.e., ATAC-seq peaks, as a proxy for putative CREs. Briefly, we used the *addGroupCoverages* function to generate pseudobulk accessibility coverage profiles for each biological replicate and cell type based on the cell grouping of “atac_general_CellType” (minCells = 100, maxCells = 1000, maxFragments = 50*1e6, minReplicates = 2, maxReplicates = 10, sampleRatio = 0.8). If a cluster did not have 100 cells (minCells = 100) from at least two samples, we allowed up to 80% of the cells from a group to be resampled with replacement to reach the required number (sampleRatio = 0.8). Then we used the *addReproduciblePeakSet* function that invoked MACS2^109^ to identify reproducible peaks that were detected in at least 2 replicates of the cell groupings (peaksPerCell = 1000, maxPeaks = 200000, minCells = 40). This procedure, which called peaks per cell type and replicate, resulted in multiple overlapping peaks from each call. To facilitate downstream analyses, we followed the iterative overlap peak merging procedure implemented by ArchR to curate a union peak set with a fixed width (500 bp). This procedure allowed the overlapping peaks being represented by the most significant ones, while avoiding chain merging of adjacent peaks or removal of peaks that were proximal but non-overlapping to the most significant peaks^110^. We then annotated if each peak in the union peak set was reproducibly detected in each cell grouping by asking for at least 20% (100 bp) overlap between the original reproducible peak and the union peak. This allowed us to obtain a reproducible peak set for each cell grouping with unified coordinates and width, which we used for all downstream analyses. The reproducible peaks were annotated as promoter (between –2000 bp and 100 bp around the TSS), intronic, exonic, and distal (intergenic) in the ArchR pipeline.

### Marker peaks and motif enrichment analysis

We identified marker peaks, i.e., ATAC-seq peaks that showed unique accessibility signals in each cell type, using the *getMarkerFeatures* function in ArchR (v1.0.2)^102^, which accounted for technical differences (TSS enrichment and fragment numbers between cell groups) and performed Wilcox tests within the union peak set. We used a cut-off of FDR <= 0.05 & Log2FC >= 1 to call marker peaks. We annotated the JASPAR2020^111^ vertebrate non-redundant motifs in the marker peaks and identified the enriched motifs from marker peaks of each cell type using the *peakAnnoEnrichment* function. The motifs were ranked by the adjusted *p*-value and those with an adjusted *p*-value < 0.001 (mlog10Padj > 3) were kept for downstream analysis. Only the cell types with more than 5000 marker peaks identified were kept for motif enrichment comparison and plotting in Extended Data Fig. 5b-d.

### Orthologous gene sets

We limit our comparative analyses to 1:1 orthologs. For most of our comparative analyses, we used human as the anchor species and mapped genes from other species to orthologs in human. We used Ensembl BioMart R package (v.2.56.1) to obtain the 1:1 orthologs between human and the other 15 species (except tenrec). Since we used transcriptome annotation from NCBI instead of Ensembl for tenrec, we built orthologous gene trees using Orthofinder^112^ (v.2.5.4) for tenrec genes. Specifically, we selected 19 species for building the orthologous gene trees, including the 15 profiled eutherian species, plus two other afrotheria (southern two-toed sloth and asian elephant), and two marsupials (gray short-tailed opossum and common wombat). We downloaded the protein fasta sequences from NCBI (tenrec) or Ensembl (the rest of the species) and used them as the input for OrthoFinder. We ran OrthoFinder with default settings and extracted the 1:1 orthologous genes between tenrec and human for our analyses. Depending on the species, we obtained 13507–21220 1:1 orthologs between human and each of the respective species. Collectively, we obtained 7250 1:1 orthologs across all 17 species (amniote ortholog set).

### Phylogenetic trees and dietary type classification

Phylogenetic distances and indicated divergence times (Fig. 1a, 2a, 4a and 4f) were obtained from TimeTree 5^113^ (https://timetree.org/). Dietary types of mammalian species (except for human) were obtained from the EltonTraits 1.0 dataset^114^ (https://opentraits.org/datasets/elton-traits.html). Specifically, we summed up the percent of diet from all plant-based sources (Diet-Fruit, Diet-Nect, Diet-Seed, and Diet-PlantO), vertebrate sources (Diet-Vend, Diet-Vect, Diet-Vfish, and Diet-Vunk), and invertebrate sources (Diet-Inv). We then classified the dietary types (Fig. 1a) based on the following criteria. Herbivore: 100% plant-based diet; Omnivore I (plant-biased): 70% <= plant-based diet < 100%; Carnivore I (insect-biased): invertebrate-based diet >= 70%; Carnivore II (vertebrate-biased): vertebrate-based diet >= 70%; Omnivore II: other cases that do not fall into the previous classifications.

### Analysis of global patterns of gene expression

#### Pseudobulk gene expression summarization

We summed up raw pseudobulk counts by each cell type and biological replicates in each species. Only the pseudobulks with a minimum of 100 nuclei were kept for downstream normalization and analysis. We filtered out non-protein-coding genes before normalization to avoid uneven dilution of normalized counts due to more extensive annotation of non-coding transcripts in some species, such as human and mouse.

To evaluate the global transcriptome patterns across liver cell types from all 17 species, we used the 3399 1:1 common orthologs that were detected in all species. Akin to our previous work, we performed a trimmed mean of M-values (TMM) normalization, implemented by the edgeR package^115^ (v.3.43.4) to achieve similar distributions of gene expression across the pseudobulks in all species. We used the TMM-normalized and log-transformed expression values for downstream analyses.

#### Principle component analysis (PCA)

Akin to previous work^116^, we used genes that were both robustly expressed and highly variable, as these genes tend to contain rich biological information but less technical noise. We considered a gene as robustly expressed within a species if it was detected in at least 5% of the cells within at least one pseudobulk. To extract highly variable genes, we used the vst standardized variance ranking calculated in the Seurat^106^ pipeline, which ranked the genes based on their cellular expression variance among all genes. We selected the 1:1 orthologous genes that also belonged to the top 5000 highly variable genes in each species and obtained the union of the highly variable genes across species. Finally, we extracted the intersection of the genes robustly expressed across all species and highly variable in at least one species (n = 959) to conduct the PCA. We used the *prcomp* function from the R stats package (v.4.3.1)^117^.

#### Gene expression tree analysis

We built nearest-neighbour joining trees for the whole liver transcriptome and different liver cell types across all species based on the distances calculated from Spearman correlation. We limited this analysis to cell types that have good representation across all species (Hepatocytes, EC, Macrophages, and NKT cells). We included all 3399 expressed amniote 1:1 orthologous genes in this analysis to retain as much ancestral and lineage-specific information as possible. Specifically, we calculated pairwise Spearman’s correlation coefficient (ρ) between the pseudobulks of the same cell type and computed their distance as 1-ρ. Then we constructed neighbour-joining trees using the R ape package^118^ (v.5.7.1) (as.phylo(root(bionj(1-ρ))) with chicken samples as the outgroup. The robustness of the branch patterning was assessed with bootstrap analyses, in which we randomly sampled the 3399 amniote orthologs with replacement 1000 times and built neighbour-joining trees upon random sampling. We calculated the bootstrap values as the proportions of the bootstrap trees that shared the branching patterns of the majority-rule consensus trees. We found that biological replicates consistently clustered together within these gene expression trees, indicating high data quality and that evolutionary differences outweighed technical and sampling variations.

### Evolutionary rate analyses on snRNA-seq data

#### Pair-wise correlation analysis after subsampling

We first subsampled UMI counts to 1500 per cell using the *SampleUMI* function from Seurat^106^ R package across the whole dataset and then randomly sampled the number of cells to 500 with replacement for each cell type in each species. We then sum up the UMI counts across each pseudobulks (cell type + species) after subsampling, performed TMM normalization, and calculated the pairwise Spearman correlation with the similarly processed pseudobulks from human. We repeated the cell subsampling process 100 times to obtain a distribution of the correlation.

We chose 1500 as the maximal UMI number for subsampling, as the large majority (82/85) of the cell type pseudobulks that we examined had a median UMI larger than 1500. We subsampled to 500 cells per pseudobulks as most species had more than 500 cells captured for each major cell type (Hepatocytes, HSC, EC, Macrophages, NKT_cells). We also tried subsampling to 750 UMI counts (the smallest median UMI across all pseudobulks: 775) and 247 cells per pseudobulk (the smallest cell number across all pseudobulks: 247), and observed very similar patterns, but with bigger variation in the 100 random sampling. We decided on 1500 UMI counts and 500 cells for subsampling to strike a balance between fair comparisons across all pseudobulks and avoiding losing too much information.

#### Sources of the snRNA-seq datasets from the other tissues

We generated new human cortex snRNA-seq data (n = 12,779) in this study. The mouse cortex data were previously published in Zaremba et al^103^. All testis data were obtained from Murat et al^104^.

#### Functional and evolutionary constraint analyses including subsampling and method comparisons

In Fig. 2b and Extended Fig. 6b, we used the LOEUF scores from the gnomAD v2.1.1 database^25^ to assess the tolerance to loss-of-function mutations of the genes expressed in each liver cell type in human. The LOEUF scores are defined as the upper bound of the 90% confidence interval for the observed/expected ratio for predicted loss-of-function (pLoF) variants^25^. High LOEUF scores indicate high tolerance to gene inactivation, while low scores indicate strong selection against the pLoF variation within the genes. For each nucleus, we calculated an average LOEUF score weighted by the log-transformed expression of the genes expressed to represent the level of the whole transcriptome. We used this approach for most of the transcriptome-based evolutionary rate analysis (except for the percentage of essential or positively selected genes) to buffer the technical noise in single-cell gene expression measurements, as lowly expressed genes were more susceptible to technical variations. We also compared other average approaches (simple mean or average weighted by expression values before log transformation) and showed that they gave very similar patterns (Extended Fig. 6a).

To assess the robustness of our analyses across cell types with varying UMIs, cell numbers, and gene capture rates, we performed subsampling on the human liver data and calculated LOEUF scores after subsampling (Extended Fig. 6b). To control for differences in UMI counts, we used the *SampleUMI* function from Seurat^106^ to subsample all nuclei to a maximum of 1,500 UMIs. Since human liver cell types had median UMI counts well above this threshold (minimum median = 2,352), this subsampling provided a stringent test. To control for differences in the number of detected genes, we excluded nuclei with fewer than 1,848 genes (the lowest median across all human liver cell types; n = 3,273). For the remaining nuclei (n = 9,214), we randomly sampled up to 1,848 genes without replacement. To control for differences in cell numbers, we subsampled each cell type to 146 cells, except for cholangiocytes (90 cells). These subsampling procedures yielded patterns consistent with those observed in the full dataset (Extended Data Fig. 6b).

In Extended Fig. 6a, we used the gene constraint scores^26^ calculated from the Zoonomia alignment across 240 mammals to assess the evolutionary constraints of the genes used in each liver cell type. Specifically, we used the fraction of coding sequences under constraint (fracCdsCons) for our calculation, as this metric has been well benchmarked in the previous publication^26^.

In Extended Fig. 6c, we used the viable gene data provided by the International Mouse Phenotyping Consortium^24^ (IMPC) release 18 to assess the usage of essential genes in each cell type in the mouse liver. Specifically, we classify both lethal and subviable (less than half of the preweaning pups survive in homozygous knockout, as described in Dickinson et al.^119^) genes as essential genes and calculate the percentage of expressed essential genes within all expressed genes. For each nucleus, the denominator is the number of the expressed genes that were accessed in IMPC, and the numerator is the number of expressed genes that were classified as essential.

We used the two-sided Mann-Whitney *U* test for testing statistical significance. We compared hepatocytes with the liver cell type with the second highest value in each species (Fig. 2b and Extended Fig. 6a-c). We also compared all liver cells with all cortex cells in Fig. 2b and Extended Fig. 6a. Results for the statistical tests were indicated in the relevant figures and provided in Supplementary Table 5.

#### dN/dS ratio

To obtain a comprehensive view of the substitute rates across different evolutionary distances and phylogenetic groups, we calculated d*N*/d*S* ratios across three lineages: mammals (armadillo, cat, pig, orangutan, chimpanzee, human, mouse, rabbit, opossum), primates (bonobo, chimpanzee, gorilla, human, macaque, marmoset), and great apes (bonobo, chimpanzee, gorilla, human, orangutan). Specifically, pairwise d*N* and d*S* values for 1:1 orthologs were downloaded from Ensembl (v99) and used to compute d*N*/d*S* ratios. For each species and each phylogenetic group, we averaged d*N*/d*S* values across all 1:1 orthologs within the group. For each nucleus, we then calculated the average d*N*/d*S* based on all genes expressed in the nucleus.

We used the two-sided Mann-Whitney *U* test for testing statistical significance. We compared hepatocytes with the liver cell type with the second-highest value in each species. Except for the rabbit analysis in Extended Data Fig. 6e, in which hepatocytes and neutrophils had the same median value, so we compared the hepatocytes with NKT cells, the third-highest cell type. We also compared all liver cells with all cortex cells in Fig. 2c and Extended Fig. 6d. Results for the statistical tests were indicated in the relevant figures and provided in Supplementary Table 5.

#### Positively selected genes

We used two sets of genes that were previously identified as carrying evidence for positive selection in primates^27^ and mammals^28^, respectively. Human gene IDs/names were obtained from the previous studies and converted to orthologous genes in all other relevant species based on our orthologous gene list (described in *Orthologous gene sets*). For great ape species, we analyzed the proportions of both the primate and mammalian positively selected gene sets (Fig. 2d and Extended Data Fig. 6g-i); for other non-primate species, we only analyzed the proportion of the mammalian positively selected gene set (Extended Data Fig. 6i,j). For each nucleus, the denominator is the number of expressed genes that were tested for signatures of positive selection, and the numerator is the number of expressed genes with evidence for positive selection.

We used the two-sided Mann-Whitney *U* test for testing statistical significance. For all species, we compared all immune cells to all non-immune cells in the livers. For all great ape species, we also compared hepatocytes to the cell type with the second-highest proportion within the non-immune cells. For the species with both liver and testis data available (human, bonobo, chimpanzee, gorilla, mouse, and opossum), we compared all liver cells to all testis cells. For the species with liver and cortex data available (human and mouse), we also compared all liver cells to all cortex cells. Results for all statistical tests were indicated in Fig. 2d and Extended Data Fig. 6g-j and provided in Supplementary Table 5.

#### Transcriptome phylogenetic age

The phylogenetic age index of genes was derived by dating their evolutionary origin, with greater weight assigned to younger genes (described in ref^8, 120^). A lower index reflects ancient origins, while species-specific genes receive the maximal score. Because the score range depends on the number of outgroup lineages available, indices are only comparable within, not across, species. Gene age indices were obtained from (http://gentree.ioz.ac.cn/) using Ensembl release 95 for human and mouse and release 69 for opossum. For each nucleus, we calculated the average transcriptome age index based on all genes expressed in the nucleus.

We used the two-sided Mann-Whitney *U* test for testing statistical significance. For each species, we compared (1) hepatocytes with all other non-immune cells (EC, HSC, cholangiocytes), and (2) immune cells with non-immune cells excluding hepatocytes. For human and mouse, we also compared all liver cells with all cortex or all testis cells. Statistical results are reported in the relevant figures and Supplementary Table 5.

#### Tissue or cell-type specificity

We used specificity or pleiotropy indices from three complementary sources: the time and tissue specificity indices derived from gene expression across seven developing organs and the pleiotropy indices that combining specificity from both developmental time and tissue dimensions^8^, tissue specificity based on bulk transcriptome data from 50 human tissues from GTEx (v8)^121,122^, and cell type specificity (cell-type tau) calculated from single-cell mRNA-seq data covering 81 major human cell types from the Human Protein Atlas^123^ (https://www.proteinatlas.org/). For the cell-type specificity indices, we downloaded the cell-type-gene expression values from the Human Protein Atlas and calculated indices based on the tau formula^124^. For all the other indices, we obtained them directly from the previous publications^8,122^.

We used two-sided Mann-Whitney *U* test for testing statistical significance. For all species and measurements (except for the time specificity in opossum, in which hepatocytes did not rank the first), hepatocytes were compared with the cell type exhibiting the second-highest specificity score, or, for pleiotropy, the second-lowest score. Results of all statistical tests are reported in the relevant figures and provided in Supplementary Table 5.

### Evolutionary rate analysis on snATAC-seq data

#### Constraint score calculation

To assess the evolutionary constraints on putative CREs, we used constraint scores derived from two different types of approaches: generative model-derived Phastcon scores^125^, which explicitly model probability of constraint within an alignment, and bottom-up approach-derived GERP++ scores^126^, which estimate constraint at individual bases and then search for clusters of highly constrained positions. We downloaded GERP++ scores from Ensembl for all the species with snATAC-seq data available. The GERP scores for the following species are derived from multiple alignment across 91 eutherian mammals from Ensembl release 106: human, chimpanzee, bonobo, gorilla, orangutan, mouse, rabbit, sheep, pig, cat, dog. For opossum, we used the GERP scores based on the multiple alignment across 58 amniota vertebrates from Ensembl release 106. For platypus and chicken, we used GERP scores based on the multiple alignment across 65 amniote vertebrates from Ensembl release 110. In addition, we also downloaded Phastcon scores from UCSC: mm10_phastCons60way (multiple alignment of 59 vertebrate genomes to the mouse genome) for mouse and hg38_phastCons100way (multiple alignment of 99 vertebrate genomes to the human genome) for human. We also calculated alignment depths (the number of genomes that targeted sequences can be aligned to) from the Zoonomia^127^ alignment across 240 mammals using the *halAlignmentDepth* function from the HAL toolkit^128^ for mouse, rabbit, sheep, dog, pig, and bonobo.

We used the *computeMatrix* function from deeptools^129^ to calculate constraint or conservation scores mentioned above across each putative CRE. Akin to the previous study^130^, we employed a sliding window approach to identify the most conserved 100 bp region of each putative CRE and used the mean score within this 100 bp region to represent the constraint level of each putative CRE. For each nucleus, we calculated the mean score of all putative CREs whose fragments were detected in the nucleus and plotted them in Fig. 2h and Extended Data Fig. 7a-r.

We used the two-sided Mann-Whitney *U* test for testing statistical significance. For all species, we compared (1) hepatocytes with all the other non-immune cells (EC, HSC, and cholangiocytes) and (2) immune cells with all non-immune cells except for hepatocytes (EC, HSC, and cholangiocytes). Results for all statistical tests were provided in Supplementary Table 5.

#### Minimal evolutionary age inference

To infer the minimal evolutionary age of each putative CRE in mouse, we traced the alignability of each region in other 18 vertebrate species spanning various evolutionary distances based on syntenic alignment, similar to the procedure employed in a previous study^130^ (Extended Data Fig. 7s). We downloaded chain alignments between mouse and each species from UCSC and used LiftOver^131^ (-minMatch=0.2 –multiple –minSizeQ=100 – minSizeT=100) to assess if at least 20% of the region could be aligned to each of the 18 species. We assess the alignability in a hierarchical manner. We first classified regions as likely originating in the common ancestor of vertebrates if it was alignable to lamprey (*Petromyzon marinus*), zebrafish (*Danio rerio*), fugu (*Takifugu rubripes*), or medaka (*Oryzias latipes*). For the rest of the regions, we then classified the ones as likely originated in the common ancestor of tetrapods if they could be aligned to one of the non-mamalian tetrapod species (western clawed frog (*Xenopus Tropicalis*), African clawed frog (*Xenopus laevis*), common garter snake (*Thamnophis sirtalis*), green anole (*Anolis carolinensis*), painted turtle (*Chrysemys picta*), zebra finch (*Taeniopygia guttata*), chicken (*Gallus gallus*)). For the rest of the regions, we then repeated this process and classified the ones as likely originating in the common ancestor of mammals if they could be aligned to one of the non-rodent mammals (platypus (*Ornithorhynchus anatinus*), opossum (*Monodelphis domestica*), African bush elephant (*Loxodonta africana*), tenrec (*Echinops telfairi*)). Lastly, for the rest of the regions, we classified them as likely originating in the common ancestor of rodents if they could be aligned to one of the rodents (guinea pig (*Cavia porcellus*), kangaroo rat (*Dipodomys ordii*), rat (*Rattus norvegicus*)). For each nucleus, we focused on the putative CREs whose fragments were detected in the nucleus, calculated the proportion of CREs with different evolutionary ages based on these detected CREs, and plotted them in the Fig. 2g,h and Extended Data Fig. 7t.

We used the two-sided Mann-Whitney *U* test for testing statistical significance. For both distal and intronic elements, we compared (1) the fraction of tetrapod and mammal-origin CREs in hepatic stellate cells and endothelial cells with those in the other cell types; (2) the fraction of rodent-origin CREs in hepatocytes and immune cells with that in the other cell types. Results for all statistical tests were provided in Supplementary Table 5.

### Analysis of the Visium spatial transcriptome data

We processed the raw sequencing data by Spaceranger (10x Genomics, v3.1.1) with default settings for demultiplexing, alignment, tissue detection, fiducial detection, and barcode/UMI counting. Then we used Seurat^106^ (v4.1.0) for further filtering (based on UMI counts, number of genes detected, percent of mitochondrial reads), normalization (*SCTransform*), dimensional reduction and clustering. We annotated the clusters using known markers from snRNA-seq data. As expected, most bins from all the liver sections displayed strong expression signatures of hepatocytes, while a small percentage of bins showed mixed signatures of non-hepatocytes. The clusters with dominating hepatocyte signatures were annotated as “Hepatocytes portal”, “Hepatocytes central”, or “Hepatocytes mid” based on the marker gene expression in mouse, opossum, and platypus samples. The clusters with non-hepatocyte signatures were annotated as “Others” in these three species in Fig. 3c-e. To search for zonation signatures in the chicken data, clustering was conducted with multiple different resolutions (0.5, 0.7, 1), but no zonation-like signatures were identified with any of the resolutions. We showed the clustering results with a resolution of 0.7 in Fig. 3f, in which most clusters had relatively clear marker gene signatures.

### Annotation of sn-RNA-seq Senegal bichir data

snRNAseq data from outgroup species was obtained from Wu et al.^9^. In brief, starting from the raw FASTQ files, we applied a processing and annotation pipeline similar to the one described above. However, the markers used for cell type annotation differed from those applied to the mammalian data. For this dataset, we used the marker genes recommended by the authors of the original study^9^.

After thorough processing of all samples, we retained only those with quality control metrics comparable to our dataset—specifically, similar average numbers of UMIs (∼2500) and genes detected (∼1500) per cell, to ensure that observed differences in downstream analyses represented true biological variation rather than differences in data quality.

As a result, we kept the Senegal bichir samples with Run Accessions SRR26534023 and SRR26534024 from NCBI project PRJNA1025373.

### Zonation marker gene comparison between mammalian and non-mammalian datasets

One-to-one orthologous genes between bichir, chicken, platypus, opossum and mouse were obtained by running Orthofinder (v2)^112^ on the proteomes of these species with parameters “-M msa” and “opt=blast”.

To investigate the presence of potential portal and central hepatocyte subtypes in the non-mammalian datasets based on marker gene expression, we began by identifying the two most transcriptionally distinct hepatocyte populations. We first subsetted the hepatocyte cluster and performed re-clustering to generate several smaller hepatocyte subclusters. Using the phylogenetic tree generated by the *BuildClusterTree* function from the Seurat package (v4)^106^, we progressively merged the most similar subclusters until only two remained. These final two subclusters were considered to represent the most transcriptionally divergent hepatocyte populations within the dataset and served as candidate portal and central subtypes. We then examined the expression of a panel of well-established zonation marker genes^33,34,39,40^ across both mammalian and non-mammalian species.

### Spatial autocorrelation metric

For assessing differences in spatial autocorrelation across species, we used the Geary’s C index. We first selected all spots annotated as hepatocytes per species in the Visium 10X data and downsampled each spot to a median of 10000 total counts with the function downsampleMatrix from DropletUtils^132^ (v1.18.0), to account for differences in sequencing depth across samples. We then took the top 200 highly variable genes per species and computed the Geary’s C index per gene with the function *runUnivariate* with type=”geary”, from the voyager package (v1.0.3)^133^. Finally, we compared the distributions of Geary’s C values across species, and assessed statistical significance using a Mann-Whitney *U* test.

### Hepatocyte heterogeneity metric

To assess hepatocyte heterogeneity within each species, we calculated pairwise hepatocyte Euclidean distances within the hepatocyte cluster. For each species, we randomly selected a single replicate to avoid confounding effects from inter-replicate variability. We then subsetted the hepatocyte cluster, identified the top 1,000 highly variable genes, and performed PCA with Seurat (v4)^106^. From each species, 500 hepatocytes were randomly sampled, and pairwise Euclidean distances were computed with the function *dist* from the stats package^117^ (v4.2.2) using the first 10 principal components. Lastly, we compared the distribution of these distances across species, and assessed statistical significance using a Mann-Whitney *U* test.

### Zonation scores

We calculated gene expression scores similarly to Sepp et al.^116^. First, data were normalized by calculating counts per million (CPM) and subsetted for the gene set of interest. Then, we scaled the genes’ expression vectors to have a mean of 0 and a variance of 1. We averaged the scaled expression of all genes of interest to compute the score and calculate its 0.01 and 0.99 percentile. Finally, we used the percentiles for capping the score to remove outliers. Values that fell out of these ranges were assigned to the nearest accepted value. We used this approach to create a “Portal score” (using portally expressed genes as input), a “Central score” (using centrally expressed genes as input), and a “Zonation score”, defined as follows:

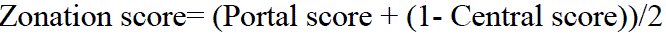

This “Zonation score” ranges from 0 (cell located close to the central vein) to 1 (cell located close to the portal vessels). The set of central and portal genes used to calculate the zonation scores were identified independently for each species using the function *FindConservedMarkers* from Seurat^106^ (v4) between the central and portal population previously annotated manually based on reference marker genes (see above). We also set the parameter grouping.var as the replicate identifier to ensure that these genes showed consistent zonation patterns across all replicates. Only the genes that met the following thresholds were kept: p_val_adj<0.05 & (avg_log2FC > 0.7 | avg_log2FC < –0.7). These can be found in Supplementary Tables 47–62.

### Ordering cells along the porto-central axis

We tried two different approaches to order cells along the porto-central axis:

● An unsupervised approach: based on pseudotime inference. Specifically, we integrated hepatocyte data from human, mouse, opossum, and platypus—representing all three major mammalian lineages—using canonical correlation analysis (CCA) as implemented in Seurat^106^ (v4). A steady-state transcriptomic zonation reference was then established by identifying a latent ordering axis using DiffusionMaps from the destiny (v3.12.0) package^134^. The first 12 CCA components were used as input for the diffusion map calculation. We subsequently applied Diffusion Pseudotime (DPT) to infer a continuous pseudotime ordering—here referred to as “pseudozonation.” The resulting pseudozonation values were normalized to range between 0 and 1 and served as our pseudozonation trajectory. This approach, however, was not applicable across all 16 mammalian species, due to its reliance on a sufficient number of 1:1 orthologs and conserved zonated genes—both of which become limiting as more distantly related species are included. Consequently, we implemented an alternative strategy described below.
● A supervised approach: based on the zonation scores calculated as stated above. Each cell was assigned a score ranging from 0 (indicative of a position near the central vein) to 1 (indicative of a position near the portal vein), reflecting its inferred location along the porto-central axis.

To assess the agreement between the supervised (zonation score–based) and unsupervised (pseudotime-based) approaches, we computed the Spearman correlation between the ordering vectors generated by both approaches for four representative species: human, mouse, opossum, and platypus. The two metrics showed substantial concordance, with correlation coefficients around 0.7 (Extended Data Fig. 8f), indicating that both approaches capture similar spatial organization along the porto-central axis.

Given this agreement, we used the supervised zonation score–based ordering for all downstream analyses.

### Detecting zonated gene expression in hepatocytes

For each species, hepatocytes were ranked by zonation score as explained above, and divided into four bins corresponding to the quartiles of the zonation score distribution. Cells in the lowest quartile (central; ≤25th percentile) and highest quartile (portal; ≥75th percentile) were aggregated into pseudobulks per replicate by summing counts. Differential expression between portal and central pseudobulks was then performed with edgeR^115^ (v3.40.0). We called genes with FDR < 0.05 as significantly zonated genes. We used the log2 fold change calculated in the edgeR pipeline as a representation of the degree of zonation, with positive values indicating portally enriched genes and negative values indicating centrally enriched genes.

### Comparison of zonated gene sets across species

For each 1:1 ortholog, we first determined the number of species in which the gene showed significant zonation (FDR < 0.05). Genes that were zonated in at least two species were then further analyzed: we identified their zonation status (zonated vs. not zonated) as well as the direction of zonation (portally-enriched vs. centrally-enriched) across species and mapped this information onto the species phylogeny. This enabled us to quantify zonation changes such as (i) gains and losses of zonation status and (ii) radical switches in zonation direction (i.e., transitions from portal to central enrichment and vice versa).

We excluded dog from the cross-species zonated gene analyses (Fig. 2a–e) because far fewer hepatocytes were captured from dog samples (mean = 578) than from other species (1,447–6,992). Such under-sampling can lead to significant bias in detecting zonated genes, which we aim to avoid. Zonated genes identified in dog are provided in Supplementary Table 26, and including dog did not alter the overall patterns observed in Fig. 2.

### Analysis of Visium HD data

#### Quality control, normalization, clustering, and annotation

We first processed the raw sequencing data with spaceranger (10x Genomics, v3.1.2) with default settings for demultiplexing, alignment, tissue detection, fiducial detection, and UMI counting. All data were aligned to transcriptomes (mouse: refdata-gex-mm10-2020-A; human and chimpanzee: refdata-gex-GRCh38-2020-A) and summarized based on the probe sets (mouse: v2.0_mm10-2020-A; human and chimpanzee: v2.0_GRCh38-2020-A.csv) provided by 10x Genomics. We imported the 8 μm binned output from spaceranger into Seurat (v5)^135^ and filtered out bins with low UMI counts captured (< 200 UMI). Since the chimpanzee and human samples (both of which were processed on the same chip in the same experimental batch) experienced pool RNA capture surrounding the center of the tissues (Extended Data. Fig. 13), 53.6% (255231/485237), 62.3 % (310796/499033) of the bins were removed after the UMI based filtering in the human and chimpanzee samples, respectively. This allowed us to achieve reasonable and comparable UMI counts for the two samples (human median UMI counts: 388, chimpanzee median UMI: 320, Extended Data. Fig. 13). We performed normalization (*NormalizeData*), sketch-based dimensional reduction, and clustering on the 8 μm bins using Seurat (v5)^135^. We annotated the clusters based on known marker genes from snRNA-seq. For zonation analysis, we focused on clusters that showed high expression of hepatocyte markers but no marker genes from non-hepatocytes to avoid signal contamination from other cell types. We then annotated the portal and central hepatocyte clusters based on the expression of zonated markers. We further visualized the spatial patterns of these clusters on the tissue sections to confirm that they fit expectation (Extended Data Fig. 10 and 13).

#### Zonated gene analysis and calling of high-confidence genes

We measured the zonation biases of all genes robustly detected in Visium HD (detected in at least 1% of the bins in at least one hepatocyte cluster) by combining two approaches. First, we aggregated all central or portal bins to calculate the normalized expression of each gene in the corresponding hepatocyte clusters. Second, we computed the percentage of bins in which each gene was detected in central or portal clusters. This binary measure was introduced to mitigate sparsity issues inherent to spatial transcriptomics at 8 μm high resolution. For each gene, we tested whether the percentage of positively detected bins differed significantly between central and portal clusters using the chi-square test. A gene was classified as centrally or portally enriched only if it exhibited the same bias in both normalized expression and bin detection percentage, and passed the chi-square test (adjusted *p*-value < 0.05). Log₂ fold changes between portal and central bins were calculated, with positive values indicating portal enrichment and negative values indicating central enrichment; these values are shown in Fig. 4g,h and 6j, and Extended Data Figs. 10 and 13. For human and chimpanzee, we had one sample for each (as a validation experiment), so we processed the data directly as described above. For mouse, Visium HD data from two samples were combined by extracting and merging central and portal bins to determine gene zonation biases.

We used the edgeR pipeline to identify zonated genes from the snRNA-seq data, as described in the section *Detecting zonated gene expression in hepatocytes*. We then obtained the genes that retained the same classifications (portal, central, and not_zonated) across both Visium HD data and snRNA-seq data, which we termed high-confidence genes. Because this analysis required agreement across multiple assays, we applied a slightly more lenient threshold in edgeR (FDR < 0.1) to reduce false negatives.

Combining snRNA-seq and Visium HD data, we detected 5,044, 4,644, and 4,443 genes robustly in both assays for mouse, human, and chimpanzee, respectively. Of these, 78.4% (3,953/5,044) in mouse, 66.3% (3,077/4,644) in human, and 88.9% (3,953/4,443) in chimpanzee maintained consistent zonation classifications across the two assays, i.e., were considered high-confidence genes.

#### Zonated gene comparison between species

For cross-species comparisons, we limited our analysis to the high-confidence genes that are also 1:1 orthologs between the species being compared. This allowed us to assess the conservation and divergence of 1,523 genes between human and mouse, and 2,107 genes between human and chimpanzee (Supplementary Tables 39 and 45). Genes were defined as conserved if they had the same zonation classification in both species, species-specific if they had different classifications and the fold-change differences between species met thresholds in both Visium HD and snRNA-seq data (abs(VisiumHD_posi_p.logFC.diff) >= 0.25 & abs(edgeR_logFC.diff) >= 0.4). Genes with differing zonation classification but insufficient fold-change differences were classified as intermediate.

We used orangutan snRNA-seq data to polarize the direction of changes for species-specific genes identified between human and chimpanzee. We used gprofiler2^136^ to perform gene set enrichment analysis on genes changed in the human or chimpanzee lineage (default settings with adjusted *p*-value < 0.1). Enriched terms from “GO:BP”, “GO:CC”, “GO:MF”, or “KEGG” categories were plotted in Fig. 6m. We chose orangutan over gorilla as the outgroup due to the relatively small number of hepatocytes captured in the gorilla snRNA-seq experiments. Nonetheless, 30 of 43 species-specific genes showed consistent polarization results using either gorilla or orangutan, which we designated as stringent genes (Supplementary Table 45). Using this stringent gene set, we still observed enrichment for lipid homeostasis pathways for genes with human-specific zonation changes.

To plot the spatial patterns of candidate genes, we used the 16 μm binned output from spaceranger (10x Genomics, v3.1.2) to mitigate the sparsity issue associated with higher resolution outputs. Data were normalized with the *NormalizeData* function in Seurat and gene expression patterns for the same cropped areas (mouse coordinates: 6000 <= x <= 13000, 3000 <= y <= 10000; human coordinates: 2000 <= x <= 10000, 2000 <= y <= 10000; chimpanzee coordinates: 16665 <= x <= 23665, 14500 <= y <= 21500) were plotted for each species in Fig. 4j and 6k.

### Quantification of metabolic activity in single cells

To assess differential metabolic activity between portal and central hepatocyte subtypes we used scCellfie^43^ (v1), which utilizes genome-scale metabolic networks to quantify metabolic activity from omics data.

In brief, for every species we used as input raw counts per cell for 1:1 orthologous genes between the species of interest and human. run_sccellfie_pipeline was run with default parameters, using the human metabolic model Recon2.2^137^ as a reference genome-scale model. For differential analysis between portal and central hepatocytes, sccellfie.stats.scanpy_differential_analysis was run between the two hepatocyte subtypes. We considered statistically significant metabolic tasks with Cohen’s D > 0.5, logFC>0.5 and adj.p-val<0.05.

### Endothelial cell subtype annotation and integration

Initial annotation of endothelial cell subtypes was done manually using known marker genes^47–49^. Endothelial cells from human, mouse, cat, pig, opossum, and platypus were then integrated using reciprocal PCA (RPCA) as implemented in Seurat^135^ (v5). We focused on species that represent distinct mammalian lineages and had sufficiently large numbers of endothelial cells (>2,500 cells) to support robust analysis. The only exception was the platypus, which, despite having fewer cells (1,460), was included due to its key phylogenetic position for inferring ancestral mammalian traits. The integrated dataset was subsequently re-clustered, and cell subtypes were re-annotated according to the expression profiles of canonical marker genes^47–49^.

### Ligand-receptor interactions between endothelial cells and hepatocytes

Ligand-receptor pairs mediating cell-cell communication events were detected between different subtypes of hepatocytes and endothelial cells along the liver lobule with CellPhoneDB (v5)^50^ using method 2 (statistical analysis). We chose this method over method 3 (differentially expression analysis) because certain hepatocyte receptors are broadly expressed across the liver lobule rather than being restricted to specific hepatocyte subtypes (i.e.: *LGR4*). As input, we provided a list with the expressed TFs for each cell type, along with matrices of normalized counts for the different subtypes of hepatocytes and endothelial cells from 6 mammalian species (human, mouse, cat, pig, opossum, platypus). Given the low recovery of central endothelial cells for some species (from 5 to 50 cells per species), we combined these cells with the central liver sinusoidal endothelial cells (LSEC_central). This analysis included genes that met at least one of the following criteria:

● Expressed in ≥5% of cells within a broad cell type (hepatocytes or endothelial cells).
● Expressed in ≥10% of cells within a specific subtype (distinct hepatocyte or endothelial cell subtypes, i.e.: VE_portal).

This approach allowed us to retain both genes with widespread expression and those that are highly expressed in a specific cell subtype, such as portal vein marker genes.

As we were interested in detecting conserved ligand-receptor interactions, only 1:1 orthologous genes between all 6 species were considered, and the default human CellPhoneDB database was used.

We then combined the ligand–receptor interaction results across species and retained those interactions that were only found either in the central or in the portal region, making them more likely to be involved in zonation, and that showed strong evolutionary conservation, defined as being detected in at least 4 species.

When assessing the expression of WNT ligands relevant for zonation, all WNT ligands were considered (including those that are non-1:1 orthologs), as we know that *WNT9B* is not present in the platypus genome but is relevant for zonation in mouse^48^.

### Gene regulatory network inference

#### Species selection

Gene regulatory network inference was performed using data from mouse, sheep, and platypus. These species were selected based on two criteria: (1) sufficient ATAC-seq resolution to robustly distinguish portal and central hepatocyte subpopulations based on gene scores for zonation marker genes, calculated from accessibility signals across their regulatory domains, and (2) representation of a broad phylogenetic span within mammals. In mouse and sheep, our single-nucleus ATAC-seq data enabled clear annotation of the two hepatocyte subtypes (central and portal), while in platypus, multiomic profiling (RNA + ATAC) facilitated their identification. Together, these species provided representatives of two distantly related eutherian lineages—rodents and even-toed ungulates— as well as a monotreme, allowing us to explore ancestral mammalian regulatory programs.

#### Pre-processing of the data

For mouse and sheep, we used Seurat (v5)^135^ to integrate the snRNA-seq and snATAC-seq modalities. We identified the 3000 most highly variable genes in the snRNA-seq dataset. These were used to construct transfer anchors between the reference set (snRNA-seq data) and the query (snATAC-seq data). We used the function *TransferData* in the snATAC-seq LSI embedding to weigh the predictions. We transferred labels for cell types from the annotated snRNAseq dataset to the snATACseq dataset.

Given the higher sparsity of our snATACseq data in comparison to the snRNA-seq data, we simplified our hepatocyte annotation of four subtypes to only two subtypes (central and portal hepatocytes). We did this by merging the most similar hepatocyte subtypes together based on the phylogenetic tree generated by *BuildClusterTree* in the snRNAseq reference dataset before label transfer.

#### pycisTopic analysis

The labelled snATACseq and multiome cells (annotated based on the transcriptome labels) and the snATACseq fragments were used as input for pycisTopic^57^, which was run with default settings except where specified below.

For each species, we generated sets of co-accessible CREs using two complementary approaches: differential accessibility and topic modeling.

To identify CREs specific to different major cell types (hepatocytes, endothelial cells, macrophages and hepatic stellate cells) we performed differential accessibility analysis across major cell types using the pycisTopic function *find_diff_features*, with adjusted *p*-value threshold of 0.05 and a log fold-change threshold of log2(1.5). We then performed differential accessibility analysis using the same parameters between hepatocyte subtypes (portal vs central), to identify CREs specific to different zonation subtypes.

To capture more complex patterns of co-accessibility, we also used a cell annotation-free approach, topic modeling. For each species, we tested models with topic numbers ranging from 15 to 60 in increments of 5. The optimal topic number, determined using multiple built-in metrics, was 40 for mouse, 40 for sheep, and 50 for platypus. We identified CREs associated with each topic by binarizing the topic-region matrix with the Otsu method and by selecting the top 3,000 regions per topic.

#### Motif enrichment analysis

To identify potential transcription factor binding sites within our CREs, we used TF motif enrichment analysis as implemented in pycisTarget^57^. For each species and each set of co-accessible CREs (DARs and binarized topics), we assessed the enrichment of over 49,000 TF motifs, grouped into more than 8,000 clusters (https://resources.aertslab.org/cistarget/motif2tf/). We used the v10 motif annotations for mouse and identified homologous motifs for sheep and platypus based on the mouse motif annotation. To ensure that all CREs were considered for TF motif enrichment, we built custom cisTarget databases for each species using create_cistarget_databases. Enrichment analysis was performed using both the cisTarget and differential enrichment of motifs (DEM) methodologies with default parameters. Motifs that obtained a normalized enrichment score (NES) >3.0 were kept.

#### SCENIC+ analysis

The gene expression matrix, the imputed accessibility from pycisTopic and the TF cistromes previously identified by motif enrichment analysis on DARs and topics with pycisTarget were used as input for SCENIC+^57^. SCENIC+ was run with the default parameters using only hepatocytes subtypes (central to portal). For our multiome dataset (platypus), we set “is_multiome” to “True” and for the paired snRNAseq and snATACseq data (mouse and sheep), we set “is_multiome” to “False”. In brief, a search space of a maximum between either the boundary of the closest gene or 150 kb and a minimum of 1 kb upstream of the TSS or downstream of the end of the gene was considered for calculating region-to-gene relationships using gradient boosting machine regression.

Region-to-gene importance scores were binarized using the 90th quantile and the top 10 regions per gene. Only region-to-gene links with correlation coefficients (rho) > 0.1 were kept. TF–region–gene triplets were generated by taking all regions that are enriched for a motif annotated to the TF and all genes linked to these regions, based on the binarized region-to-gene links.

The last step of the SCENIC+ pipeline, where Gene Set Enrichment Analysis (GSEA) is performed to rank genes based on their TF-to-gene importance score and assess enrichment of genes within TF–region–gene triplets, was deliberately skipped, as we know that for some TFs (particularly those involved in WNT signaling) expression between TF and target genes does not necessarily show a high correlation. eRegulons with at least 30 target genes were kept, obtaining 337, 323 and 259 eRegulons.

We reasoned that for a TF to be important for zonation it should have a considerable proportion of zonated target genes, so for every species we ranked the TFs expressed in hepatocytes by the proportion of portal and central targets in its eRegulon, and compared these ranking across species to select the TFs with conserved high ranks across species (rank>90th quantile across species).

Gene-based eRegulons were scored in the relevant datasets using AUCell (v1.20.1)^138^.

#### Analysis of TCF7L2 ChIPseq data

For validating the TF-CRE links in the TCF7L2 eGRN, we relied on previously published ChIP-seq data for this TF^58^. In brief, high-quality ChIP-seq peaks were converted from mm10 to mm39 using LiftOver^139^. We then intersected these peaks with the differentially accessible regions (DARs) identified between central and portal hepatocytes with the function *findOverlaps* from the GenomicRanges^140^ (v1.50.2) package to assess the extent to which TCF7L2 directly binds regions showing zonation-dependent chromatin accessibility.

### Relative gene expression changes: shifts of cell-type specific expression

We focused on the great ape phylogeny and included mouse as the outgroup. We aggregated pseudobulk UMI counts across each cell type and each species, calculated counts per million (CPM) values, and performed log transformation. We only kept the major cell types common to all species (Hepatocytes, EC, HSC, Macrophages, and NKT cells) and conducted z-score scaling within each species. We used the scaled expression values as input for the downstream analysis.

#### Ancestral inference and calling species-specific changes

We used the fuzzy-c-means package^141^ (v1.7.0) to apply a soft-clustering approach on the 1:1 orthologous genes that were highly variable across the liver cell types in all six species (n = 1986). This approach assigned genes into different clusters with certain probabilities based on their relative expression in each cell type. Similar to previous studies^8,104^, we inferred within a phylogenetic framework the probability that there were changes in the cluster assignments of orthologous genes in specific branches (Extended Data Fig. 13a). Specifically, we first inferred the assignment probabilities of the internal ancestral nodes based on the child nodes, calculated as the average of the child nodes inversely weighted by the tree branch lengths, so that the closer child node has more weight. To achieve high-confidence ancestral inference, we excluded genes with highly divergent expression patterns in the child nodes by implementing two criteria. First, the pairwise correlation of the cluster assignment probabilities of the two child nodes should be higher than 0.4. This ensures that the z-score distributions (relative expression patterns) of the child nodes are similar enough. Second, for each cell type, the mean expression values of the two child nodes are not very different, i.e., within all species involved, the standard deviation (SD) across the species of each cell type should all be lower than 1.5.

Then we compared neighbouring branches to identify genes switching patterns. We summed up the probabilities across all clusters in the neighbouring nodes (node A and node B) as the fuzzy co-clustering probability score based on the following formula, akin to the idea of dot product.

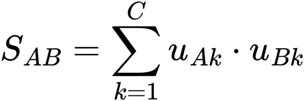

μ*_Ak_* is the membership probability of node *A* in cluster *k,* μ*_Bk_* is the membership probability of node *B* in cluster *k*, *C* corresponds to the total number of clusters. A larger co-clustering score S_AB_ indicates that both nodes tend to belong to the same clusters with high probabilities. We called genes with a co-clustering probability score smaller than 0.1 (less than 10% probability of sharing the same cluster assignment) as the ones with switched expression patterns. Lastly, we used the most closely related outgroup to polarize the direction of gene expression changes. We also made sure that the expression patterns in the outgroup and the unchanged node were similar enough by using the same two criteria that we implemented in the ancestral inference step.

To call conserved genes between two neighbouring branches, we required the following criteria: pairwise correlation of the clustering probabilities of the two branches > 0.6, co-clustering probability score (S_AB_) > 0.4, and the total variation distance of the clustering probabilities < 0.3. If one of the branches was an internal node, we also required the conserved genes to be conserved across all species used for ancestral inference.

#### Determining the most affected cell types

To determine the cell types that were mostly affected in the relative gene expression switches, we calculated the Δ z-score, z-score differences between the changed node and the ancestral node (Δ z-score = z-score_changing_node_ – z-score_ancestral_node_). For internal nodes, we inferred the z-score based on the child nodes as described above. We referred to the cell type with the biggest absolute z-score differences (abs(Δ z-score)) as the maximally affected cell type. We then calculated the non-scaled expression differences of the maximally affected cell type between the changed node and the ancestral node (Δ normExp = normExp_changing_node_ – normExp_ancestral_node_) and classified the genes with a positive Δ normExp as up-regulated genes, a negative Δ z-score as down-regulated genes.

#### Cross-reference with the snATAC-seq data

To validate the conserved and divergent genes with the snATAC-seq gene scores, we calculated pseudobulks and performed normalization and z-score scaling of the snATAC-seq gene scores for each gene in each species, similar to the pipeline that we applied to the snRNA-seq data. We then calculated the gene score-based Δ z-score for each candidate gene and plotted them together with the gene expression-based Δ z-scores (Fig. 6d). We also checked if the cell types with the extreme z-scores (abs(z-score) being the largest) in the changing node were the same in the snRNA-seq gene expression and snATAC-seq gene score data. We found 31/41 of the candidate genes fulfilled this criterion, indicating overall concordant patterns in snRNA-seq and snATAC-seq data.

#### Linking putative CREs to target genes

Putative CREs identified in the great ape species were associated with their potential target genes based on correlation between peak accessibility and gene expression from the snRNA-seq data. Specifically, we run the *addPeak2GeneLinks* function from ArchR^102^ to search for positively correlated peaks within 500 bp of the transcription start site (k = 50, knnIteration = 500, overlapCutoff = 0.5, maxDist = 5e5). We exported significantly associated peaks using the *getPeak2GeneLinks* function with default settings (corCutOff = 0.45, FDRCutOff = 1e-04).

#### Identifying orthologous sequences of putative CREs

We illustrate the pipeline using human putative CREs as an example. We used the Zoonomia^127^ alignment to identify the orthologous regions of human putative CREs in bonobo, chimpanzee, and orangutan. Specifically, for each CRE, we first performed base-by-base mapping between human and the other species using the *halLiftover* function from the HAL tool kit^128^ and then constructed contiguous orthologs using HALPER^142^. For bonobo and orangutan, the HALPER output was directly compatible with our snATAC-seq analysis, so these regions were used as orthologs. For chimpanzee, we further converted PanTro6 coordinates (used in Zoonomia) to PanTro5 coordinates (used in our ATAC-seq analysis) via UCSC LiftOver^131^ (default settings). Reciprocally, we applied the same workflow to map putative bonobo, chimpanzee, and orangutan CREs to the other great ape species, including human. For gorilla, we directly used UCSC LiftOver^131^ (default settings) to identify orthologous regions of human putative CREs in the gorilla gorGor4 genome (and vice versa) due to compatibility issues between the assembly used in our analysis and in Zoonomia.

### Putative CREs with human-specific activities

To call human-specific putative CREs, we first assessed if the orthologous regions of human putative CREs in species A overlapped the putative CREs identified in the same cell type in species A. We used BEDtools^143^ (v2.31) intersect to determine if the overlap existed. We limited our analysis to robustly detected ATAC-seq peaks (accessible in at least 2% of the nuclei in at least one cluster) in all species to exclude weakly accessible regions with high noise. We focused on common liver cell types detected in snATAC-seq data in all great ape species. For each cell type, we considered human putative CREs with orthologous regions in all the other great apes but not overlapping accessible regions in any of the species as human-specific putative CREs.

We used GREAT^144^ (v4) to predict the functional enrichment of human-specific CREs with default settings and plotted the top 5 GO biological process terms passing the default threshold (BinomFdrQ < 0.05 & HyperFdrQ < 0.05 & RegionFoldEnrich >=2) in Fig. 6i.

## Acknowledgements

We thank Walter Salzburger, Patrick Tschopp, Marcus Clauss, Antoine Fages, Marcin Falis, Sophie Kraunsoe, Simon Anders, Ioannis Sarropoulos, Mari Sepp, and all members from the Kaessmann Lab for discussion and suggestion; K. Hall for assistance; Deep Sequencing Core Facility at Heidelberg University and the MULTI-SPACE Platform from health-life science alliance Heidelberg Mannheim for providing support and funding for the spatial transcriptomic experiments; Genomics Core Facility (GeneCore), EMBL, Heidelberg for providing sequencing service for multiome libraries; Michel Gex and Slaughterhouse Abattoir Vevey, Switzerland for providing the sheep and pig liver samples; Philipp Khaitovich and Tissue Bank for Developmental Disorders at the University of Maryland (USA) for providing human samples; Isa Lindgren and Linköping University for providing chicken samples; Miguel Carneiro and CIBIO-InBIO research center in biodiversity and genetic resources for providing rabbit samples; Ulrich Zeller and Humboldt University of Berlin, John L. VandeBerg and The University of Texas Rio Grande Valley for providing opossum samples. We acknowledge support by the state of Baden-Württemberg through bwHPC and the German 789 Research Foundation (DFG) through grant INST 35/1597-1 FUGG. We acknowledge the data storage 790 service SDS@hd supported by the Ministry of Science, Research and the Arts Baden-Württemberg 791 (MWK) and the German Research Foundation (DFG) through grant INST 35/1503-1 FUGG. The purchase of the NextSeq 550 instrument was supported by the Klaus Tschira Foundation. This research was funded by a Sinergia grant from the Swiss National Science Foundation (grant 189970) to H.K; by an add-on fellowship of the Joachim Herz Stiftung to L.R.-M.; A.F. was supported by the Swedish Research Council for Sustainable Development (Formas) (2021-00513); M.C.-M was supported by The Francis Crick Institute, which receives its core funding from Cancer Research UK (grant CC2185 to M.C.-M.), the UK Medical Research Council (grant CC2185 to M.C.-M.), and the Wellcome Trust (grant CC2185 to M.C.-M; M.P. was supported by a grant from the Hungarian Brain Research Program (NAP2022-I-4/2022) of the Hungarian Academy of Sciences and the Thematic Excellence Program of the Semmelweis University.

## Author contributions

X.Y., L.R.-M., and H.K. conceived and organized the study;

X.Y. performed most experiments with support from J.S., C.S., B.B., D.I.; B.Z. generated and analyzed the snRNA-seq data from human prefrontal cortex.

X.Y. and L.R.-M performed all analyses with support from E.L. and N.T.;

M.C.-M. and H.K. provide key feedback and discussions;

B.N., M.P., R.T., U.Z., J.V., G.B., J.L., A.F., F.G., A.T., M.M., S.P. provided the samples.

X.Y., and L.R.-M. wrote the original manuscript, with critical review by B.Z, M.C.-M., and H.-K.

## Data Availability

snRNA-seq data produced in this study are available on ArrayExpress under the following accession number: E-MTAB-15509 (rabbit and guinea pig, reviewer access: https://www.ebi.ac.uk/biostudies/arrayexpress/studies/E-MTAB-15509?key=c93047f4-7b2a-43f6-adfe-8de2d7ea59c9), E-MTAB-15545 (human, reviewer access: https://www.ebi.ac.uk/biostudies/arrayexpress/studies/E-MTAB-15545?key=2f7244eb-3410-4f37-82e4-e57a28f5ff50), E-MTAB-15548 (chimpanzee and bonobo, reviewer access: https://www.ebi.ac.uk/biostudies/arrayexpress/studies/E-MTAB-15548?key=35b111f3-8273-4c43-b4e0-9eef5ab4cfb2), E-MTAB-15549 (gorilla and orangutan, reviewer access: https://www.ebi.ac.uk/biostudies/arrayexpress/studies/E-MTAB-15549?key=6e3a9fad-6545-4bf7-ba11-20ce3e815ca9), E-MTAB-15543 (sheep and pig, reviewer access: https://www.ebi.ac.uk/biostudies/arrayexpress/studies/E-MTAB-15543?key=0caec76d-4968-4278-af99-8e588eeb6b21), E-MTAB-15544 (cat and dog, reviewer access: https://www.ebi.ac.uk/biostudies/arrayexpress/studies/E-MTAB-15544?key=42db14da-468f-4601-a009-41dc99e62128), and E-MTAB-15547 (tenrec, armadillo, and opossum, reviewer access: https://www.ebi.ac.uk/biostudies/arrayexpress/studies/E-MTAB-15547?key=ed1dd691-2a5c-4b5b-8f19-1aea0857a6f9), E-MTAB-15550 (chicken and platypus, snMultiome, reviewer access: https://www.ebi.ac.uk/biostudies/arrayexpress/studies/E-MTAB-15550?key=f436ca6b-9ff5-4617-ae65-748167d1d2bb). Previously published mouse snRNA-seq data are available under E-MTAB-12180.

snATAC-seq data produced in this study are available on ArrayExpress under the following accession number: E-MTAB-15541 (human, reviewer access: https://www.ebi.ac.uk/biostudies/arrayexpress/studies/E-MTAB-15541?key=c7a543f1-1f38-4343-9ced-73a3e57f6af1), E-MTAB-15546 (chimpanzee, bonobo, gorilla, and orangutan, reviewer access: https://www.ebi.ac.uk/biostudies/arrayexpress/studies/E-MTAB-15546?key=5409adb4-dae3-456b-b5e1-17a9c409d5dd), E-MTAB-15540 (rabbit, mouse, and opossum, reviewer access: https://www.ebi.ac.uk/biostudies/arrayexpress/studies/E-MTAB-15540?key=a6c91210-6813-495a-9a2f-ebbfa7c2571c), E-MTAB-15542 (sheep, pig, cat, and dog, reviewer access: https://www.ebi.ac.uk/biostudies/arrayexpress/studies/E-MTAB-15542?key=9121ec86-0f8c-4280-a99f-e8cdedef44c0), and E-MTAB-15539 (platypus and chicken, snMultiome, reviewer access: https://www.ebi.ac.uk/biostudies/arrayexpress/studies/E-MTAB-15539?key=80662f29-0bf6-442d-84a7-48de0eb81ef2).

Bulk mRNA-seq data produced in this study are available on ArrayExpress under E-MTAB-15538 (reviewer access: https://www.ebi.ac.uk/biostudies/arrayexpress/studies/E-MTAB-15538?key=3e58028f-81e0-405d-825f-df3eabb510c3). Spatial transcriptomics data produced in this study are available on ArrayExpress under the following accession number: E-MTAB-15553 (10x Visium regular, mouse, opossum, platypus, and chicken, reviewer access: https://www.ebi.ac.uk/biostudies/arrayexpress/studies/E-MTAB-15553?key=e5d1101a-9877-44ce-bc5a-6e851d52fd66), E-MTAB-15552 (10x Visium HD, mouse and chimpanzee, reviewer access: https://www.ebi.ac.uk/biostudies/arrayexpress/studies/E-MTAB-15552?key=c8af7567-44cf-4ab7-94c9-905b05f3fba7), and E-MTAB-15551 (10x Visium HD, human, reviewer access: https://www.ebi.ac.uk/biostudies/arrayexpress/studies/E-MTAB-15551?key=aed2ca00-1345-4bdf-9a01-5005eb2e4f4c).

Genome-wide chromatin accessibility profiles for the 13 species are available on UCSC genome browser:

● human (https://genome.ucsc.edu/s/Xuefei/hg38_liver_full)
● chimpanzee (https://genome.ucsc.edu/s/Xuefei/panTro5_liver_full)
● bonobo (https://genome.ucsc.edu/s/Xuefei/panPan2_liver_full)
● gorilla (https://genome.ucsc.edu/s/Xuefei/gorGor4_liver_full)
● orangutan (https://genome.ucsc.edu/s/Xuefei/orangutan_liver_full)
● mouse (https://genome.ucsc.edu/s/Xuefei/mm39_liver_full)
● rabbit (https://genome.ucsc.edu/s/Xuefei/oryCun2_liver_full)
● cat (https://genome.ucsc.edu/s/Xuefei/felCat9_liver_full)
● dog (https://genome.ucsc.edu/s/Xuefei/dog_liver_full)
● pig (https://genome.ucsc.edu/s/Xuefei/susScr11_liver_full)
● sheep (https://genome.ucsc.edu/s/Xuefei/sheep_liver_full)
● opossum (https://genome.ucsc.edu/s/Xuefei/monDom5_liver_full)
● platypus (https://genome.ucsc.edu/s/Xuefei/platypus_liver_full)
● chicken (https://genome.ucsc.edu/s/Xuefei/chicken_liver_full)

Other processed data can be interactively explored and downloaded at a web application (https://apps.kaessmannlab.org/liver_app/).

## Code Availability

Custom scripts for analyses reported in this manuscript are available at https://gitlab.com/kaessmannlab/liver-paper.

---

**Extended Data Fig. 1.**
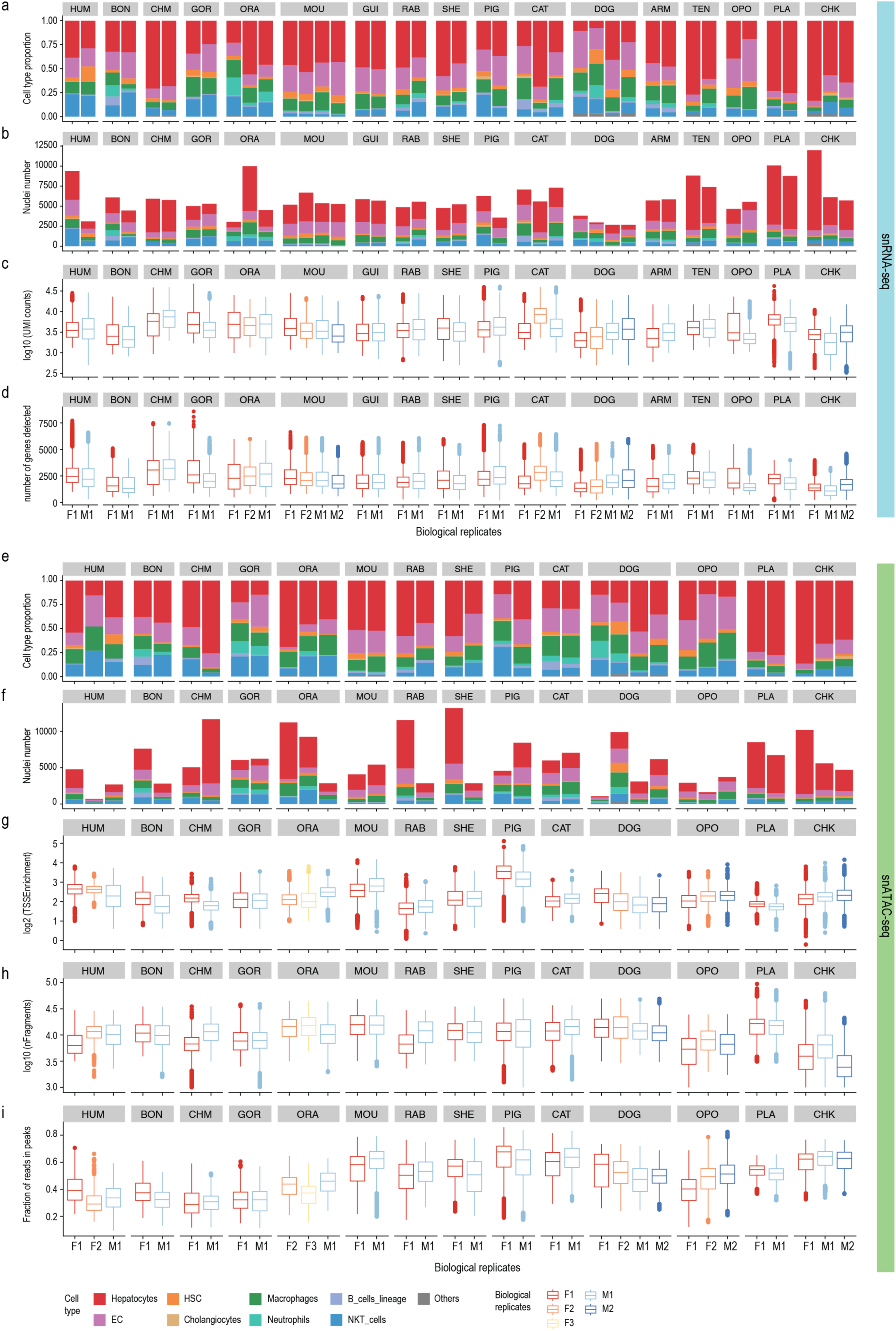
| Quality control of snRNA-seq and snATAC-seq datasets. **a-d**, Quality control of snRNA-seq data. Cell type proportion **(a**), number of nuclei (**b**), UMI counts (**c**), and number of genes detected (**d**) of each biological replicate. **e-i**, Quality control of snATAC-seq data. Cell type proportion (**e**), number of nuclei (**f**), TSS enrichment (**g**), number of fragments detected (**h**), and fraction of reads in peaks of each biological replicate. F: female; M: male.

**Extended Data Fig. 2.**
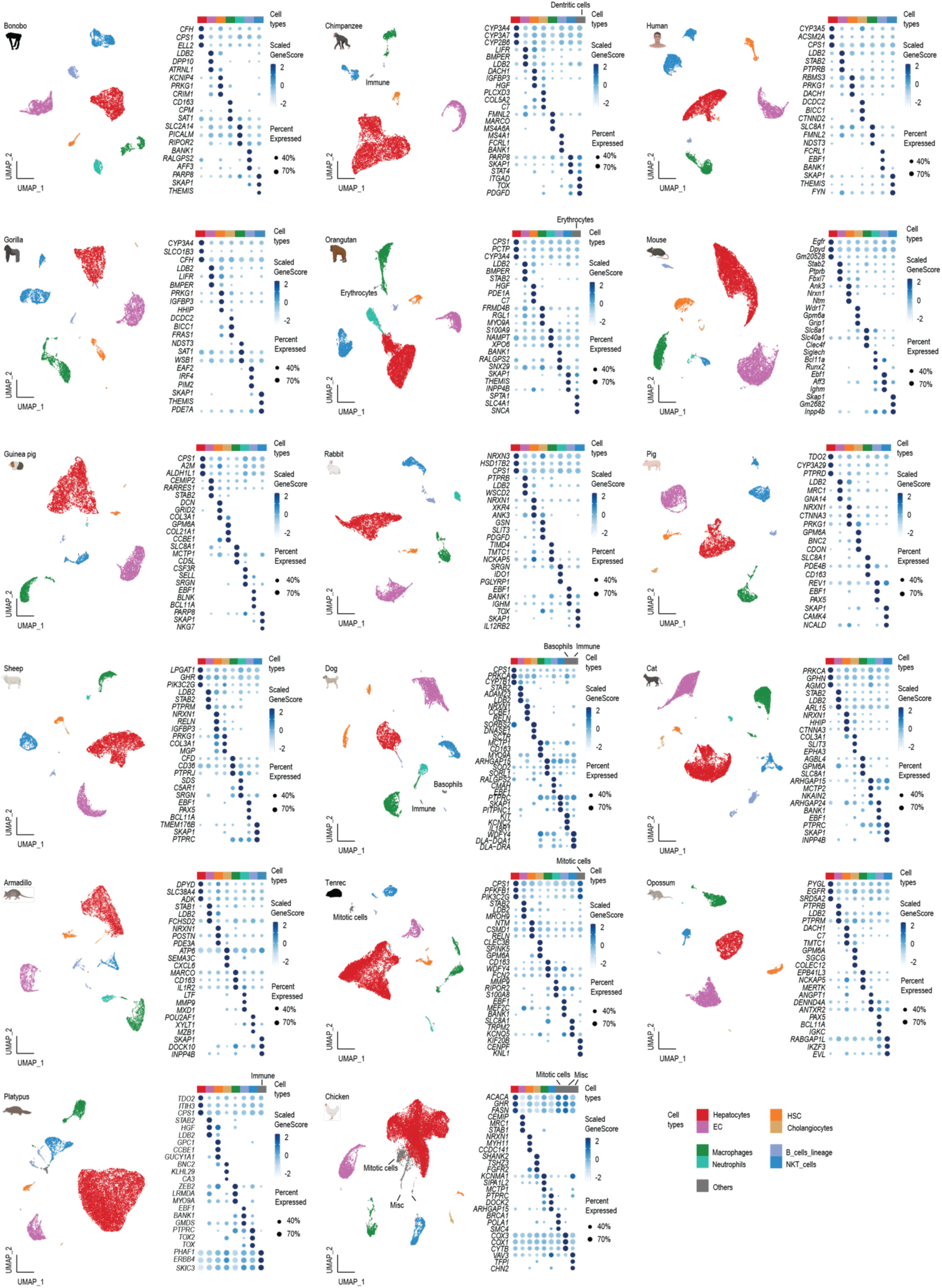
| Overview of the snRNA-seq dataset for each species. For each species, two plots are shown: UMAP visualization of the cell-type clusters (left) and dotplot showing the scaled gene expression of the top 3 marker genes for each cluster (right).

**Extended Data Fig. 3.**
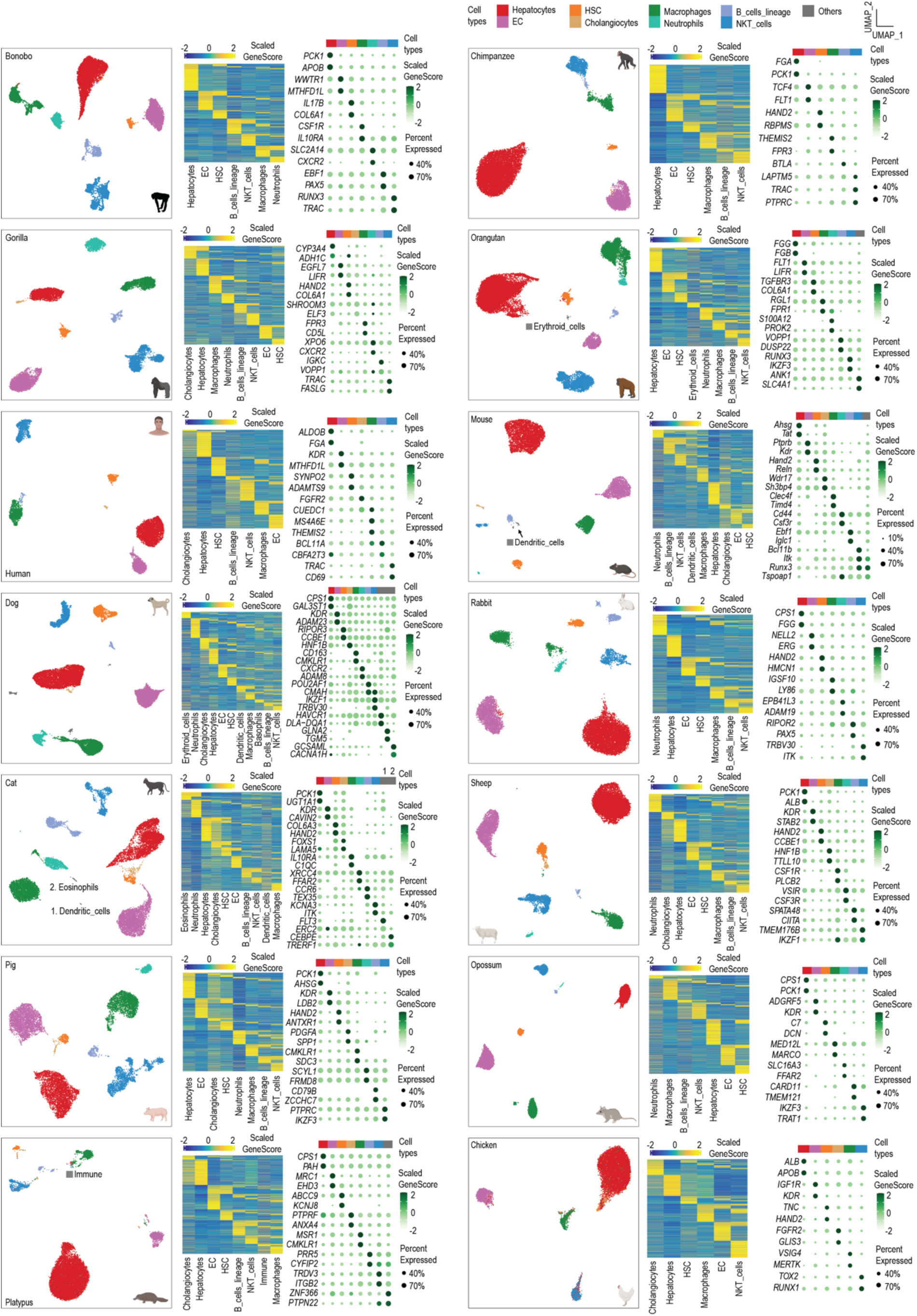
| Overview of the snATAC-seq dataset for each species. For each species, three plots are shown: UMAP visualization of the cell-type clusters (left), heatmap showing chromatin accessibility-inferred gene activity scores (gene scores) of all marker genes (middle), and dotplot showing the gene scores of the top 2 marker genes for each cluster (right). In the heatmaps, each row represents a gene

**Extended Data Fig. 4.**
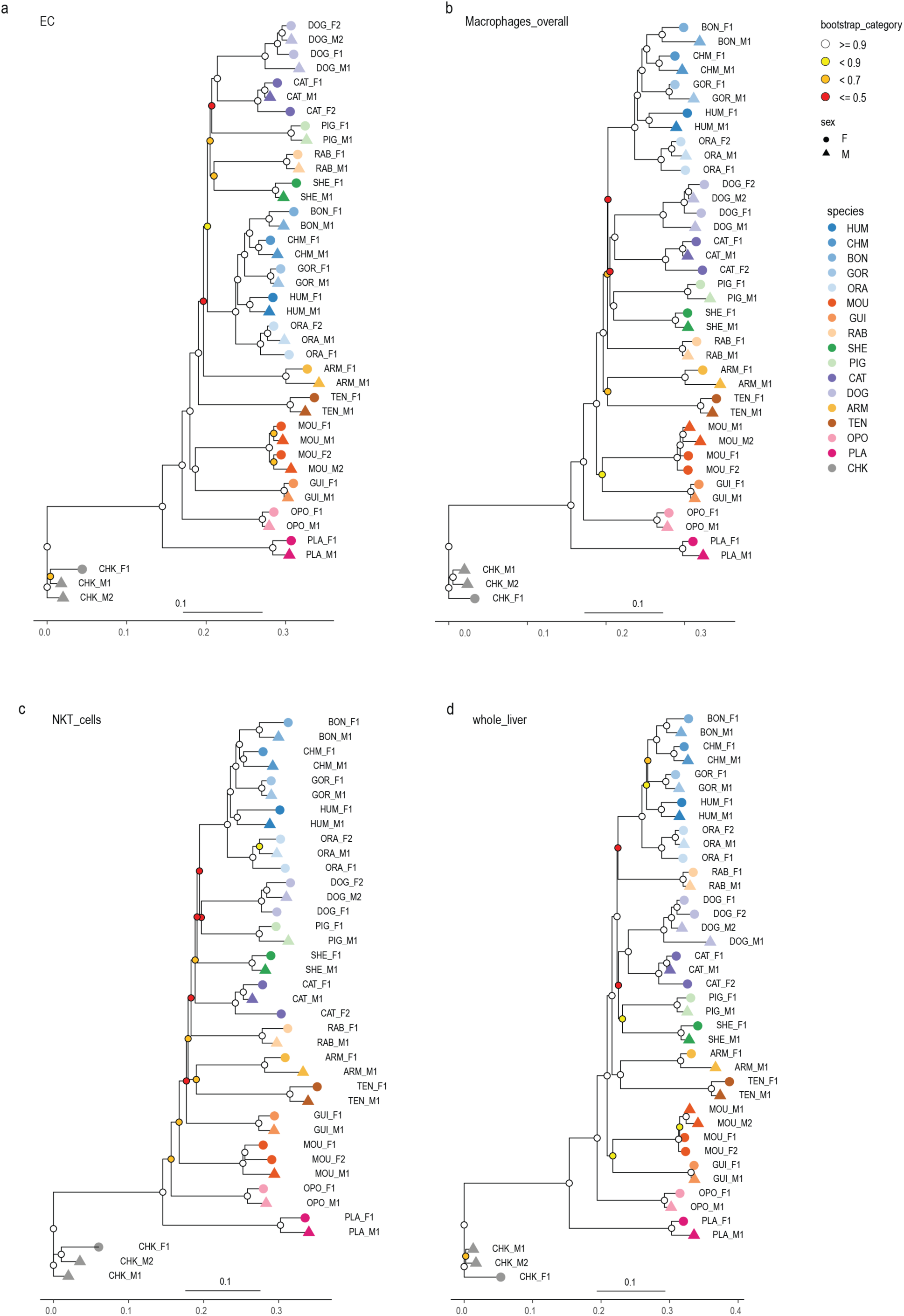
| Gene expression trees of hepatic cells. Gene expression tree based on pseudobulk transcriptomes for endothelial cells (**a**), macrophages (**b**), NKT cells (**c**), and whole liver (**d**) transcriptomes. Bootstrap values (3,399 1:1 orthologous genes were randomly sampled with replacement 1,000 times) are indicated by circles.

**Extended Data Fig. 5.**
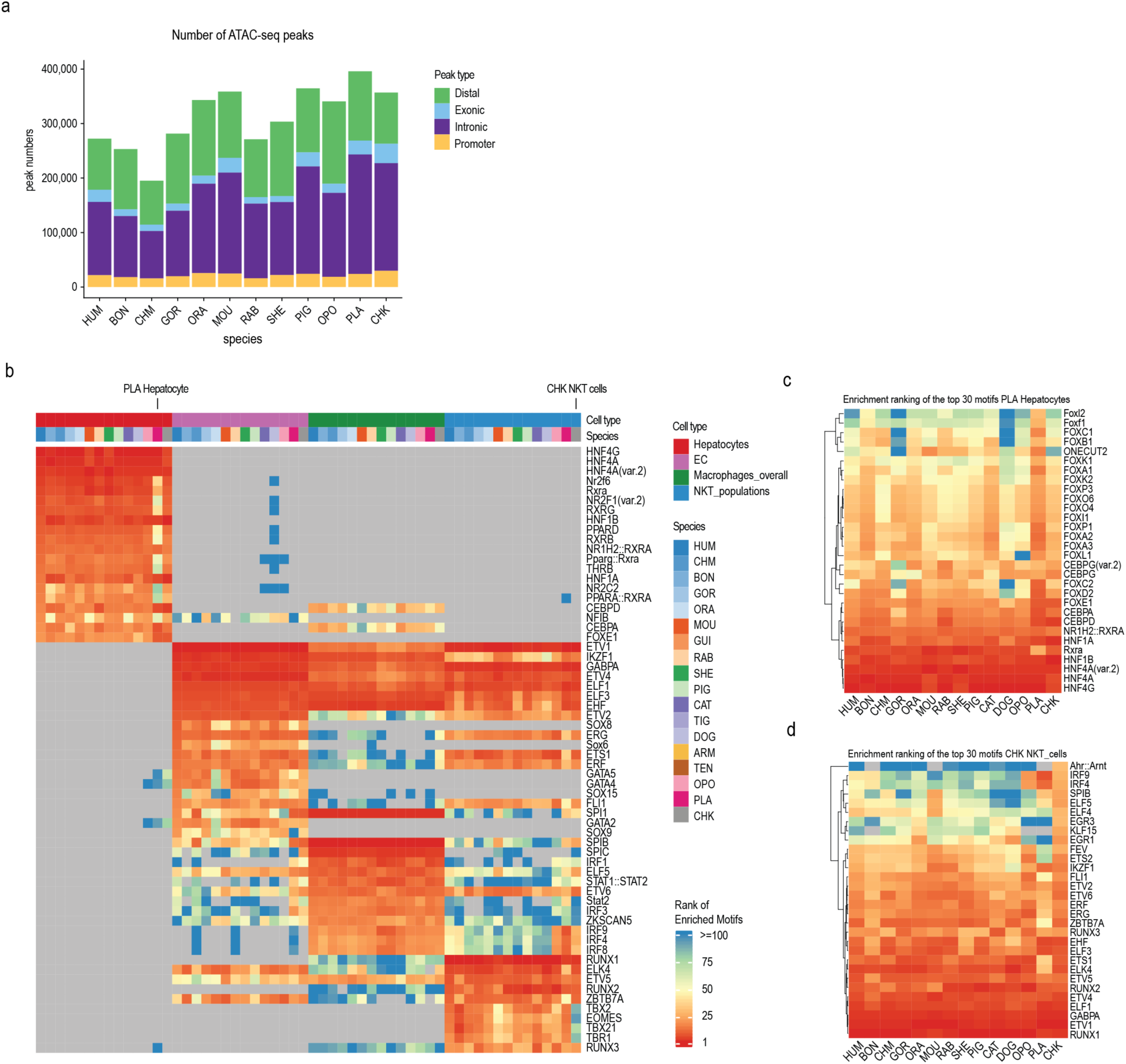
| Overview of putative CREs and motif enrichment in mammalian liver cells. **a**, Numbers of ATAC-seq peaks identified in each species based on the iterative procedure in ArchR. **b,** Ranking of top 20 enriched motifs (adjusted *p*-value < 0.001) identified from the marker peaks of hepatocytes, endothelial cells, macrophages, and NKT cells in human, plotted together with the enrichment ranking of the same motifs in the other species. Grey indicates no significant enrichment. **c**, Ranking of the top 30 enriched motifs identified from the hepatocyte marker peaks in platypus, plotted together with the enrichment ranking in the hepatocytes from other species. **d**, Ranking of the top 30 enriched motifs identified from the NKT cell marker peaks in chicken, plotted together with the enrichment ranking in the NKT cells from other species.

**Extended Data Fig. 6.**
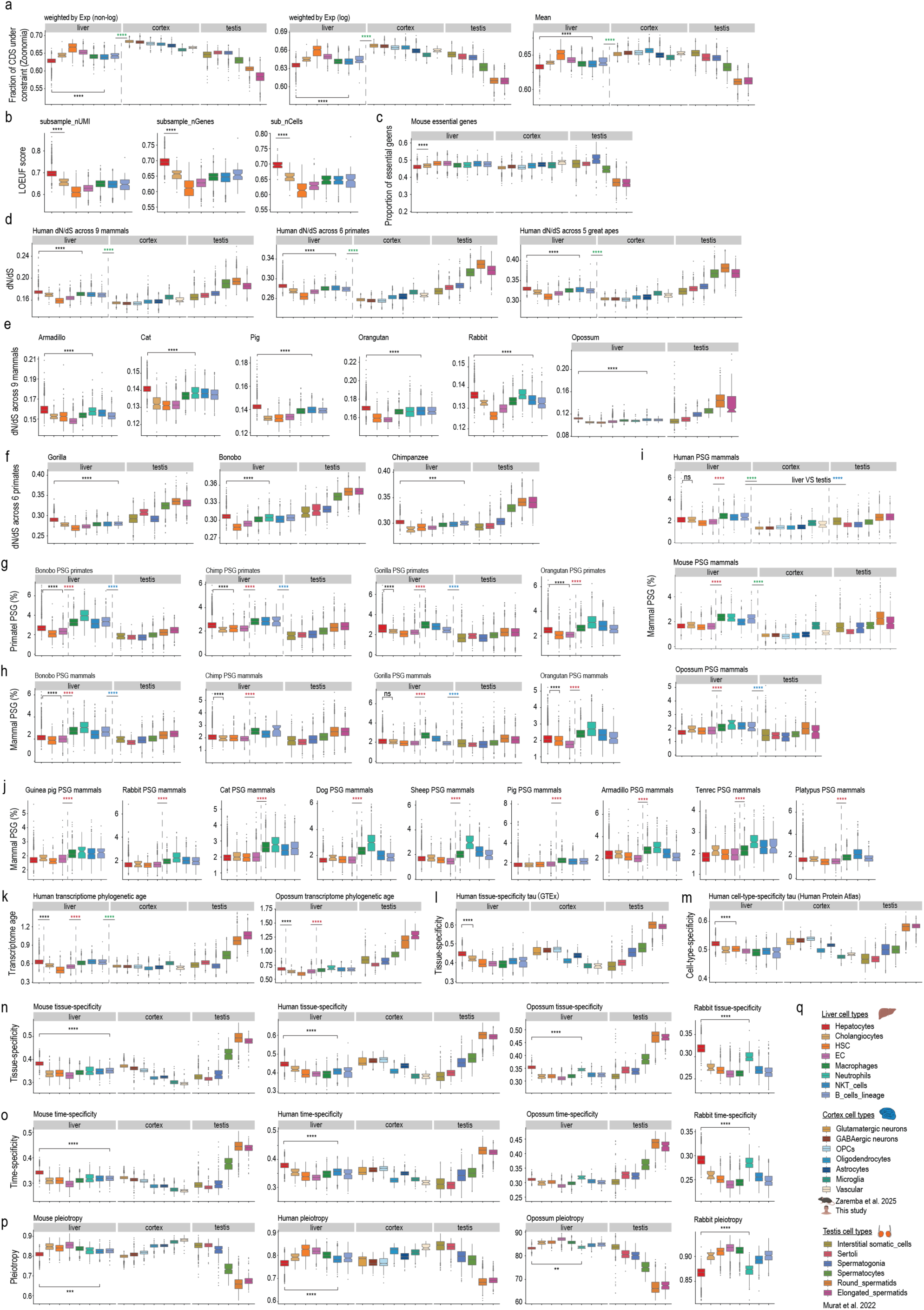
| Transcriptome-based evolutionary rates. **a**, Comparisons of three different approaches for calculating the average evolutionary rate value (here using the fraction of CDS under constraint from Zoonomia alignment as an example) for each cell: Simple mean of the genes expressed in a given cell (right), average weighted by the log-transformed expression of the genes in a given cell (middle), average weighted by the non log-transformed expression of the genes in a given cell (left). All approaches showed a similar trend. The approach shown in the middle, average weighted by the log-transformed expression of the genes in a given cell, was used for all relevant analyses. **b**, Evolutionary rate analysis upon different types of subsampling, using the LOEUF score as an example. Left: all nuclei were subsampled to the same UMI count (n = 1500); middle: all nuclei were subsampled to the same number of genes detected (n = 1848); right: all cell types were subsampled to the same number of cells (n = 146) except for cholangiocytes (n = 90). All subsampling showed results similar to those from the full data (Fig. 2b). **c**, Percentage of expressed genes leading to a lethal or sub-viable phenotype when knocked out in mouse embryos. **d**, Comparison of d*N*/d*S* ratio analysis based on d*N*/d*S* values from different evolutionary lineages, with human cell type as an example. **e**, d*N*/d*S* ratio (across 8 mammals) of genes expressed in each cell type in six different species. **f**, d*N*/d*S* ratio (across 6 primates) of genes expressed in each cell type in three great ape species. **g**, Percentage of expressed genes with signs of being positively selected in the primate lineage. **h-j**, Percentage of expressed genes with signs of being positively selected in the mammalian lineage. **k**, Transcriptome phylogenetic ages of genes expressed in each cell type. **l**, Tissue-specificity index (based on GTEx data) of genes expressed in each cell type in human. **m**, Cell-type-specificity index (based on single-cell data from Human Protein Atlas) of genes expressed in each cell type in human. **n**, Tissue-specificity index (based on mammalian transcriptome data across seven organs) in each cell type. **o**, Time-specificity index (based on mammalian liver developmental data) in each cell type. **p**, Pleiotropy index (based on mammalian organ developmental data) of the genes expressed in each cell type. **q**, Cell type color code used in this figure. **a-p**, Boxplots display the median (center value) and upper and lower quantiles (box limits) with whiskers at 1.5 times the interquartile ranges. ****: *P* < 0.0001, ***: *P* < 0.001, **: *P* < 0.01, ns: *P* > 0.05. Results for all statistical tests are shown in Supplementary Table 5. Green stars: all liver cells compared with all cortical cells; blue stars: all liver cells compared to all testicular cells; red stars: immune cells compared to non-immune cells in livers.

**Extended Data Fig. 7.**
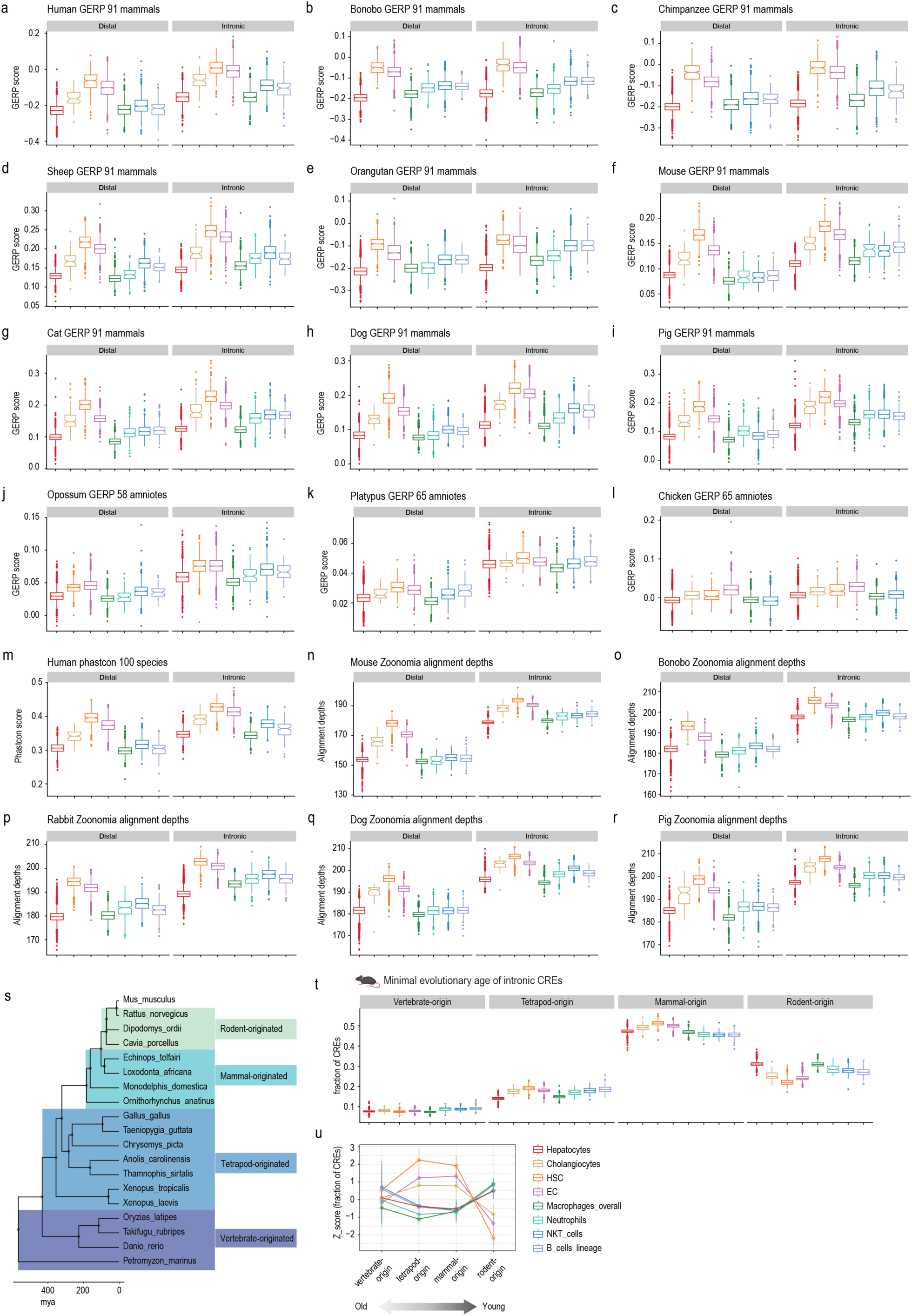
| Evolutionary rates of putative CREs. **a-i**, GERP scores (across 91 mammals) of the putative distal or intronic CREs active in each cell type in each species. **j**, GERP scores (across 58 amniotes) of the putative distal or intronic CREs active in each cell type. **k, l**, GERP scores (across 65 amniotes) of the putative distal or intronic CREs active in each cell type. **m**, Phastcon scores (across 100 vertebrates) of the putative distal or intronic CREs active in each cell type in human. **n-r**, Zoonomia alignment depths of the putative distal or intronic CREs active in each cell type. **s**, Phylogenetic tree of the species used for minimal evolutionary age inference for mouse CREs. **t**, Fraction of intronic CREs with different minimal evolutionary ages in mouse liver cells. **u**, Z-score of the fraction of intronic CREs with different minimal evolutionary ages scaled across all mouse nuclei. **a-r**, **t**, Results for all statistical tests are shown in Supplementary Table 5.

**Extended Data Fig. 8.**
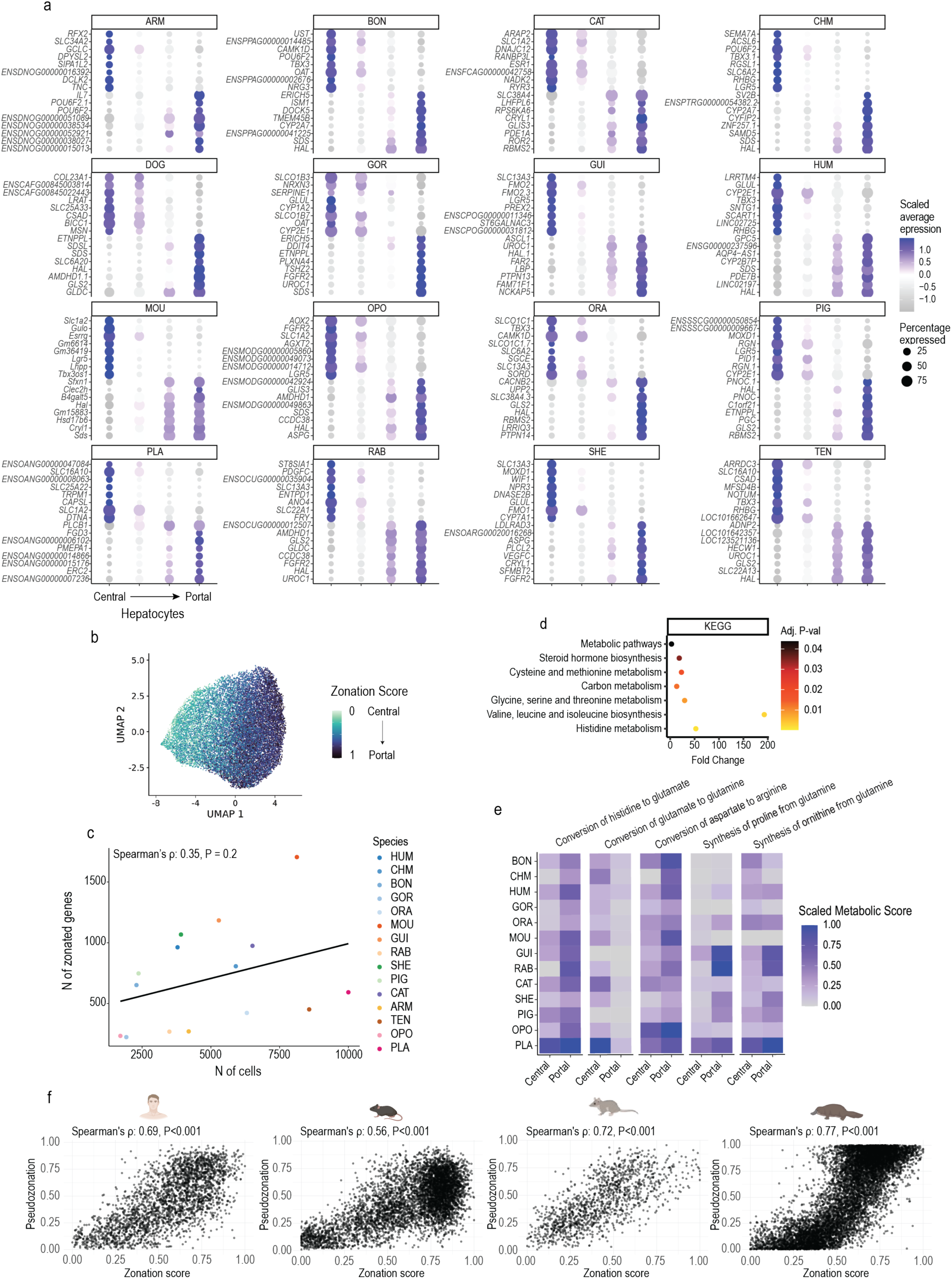
| Detection of zonated genes and metabolic functions. **a**, Top 16 zonated genes per species. **b**, UMAP of integrated hepatocytes across human, mouse, opossum, and platypus. Cells are colored by zonation score. **c**, Correlation between the number of hepatocytes and the number of zonated genes detected per species. Pearson’s correlation coefficient and *p*-value are indicated in the top left corner. **d**, KEGG pathway enrichment analysis of highly conserved zonated genes (zonated in > 8 species) (Benjamini–Hochberg-adjusted *P* < 0.05, hypergeometric test). **e**, Scaled metabolic scores in portal and central hepatocytes for the five most conserved and differentially active metabolic tasks. **f**, Scatter plots of zonation scores and pseudotime-based pseudozonation in human, mouse, opossum, and platypus. The Spearman correlation coefficient (ρ) and associated *p*-value are shown at the top.

**Extended Data Fig. 9.**
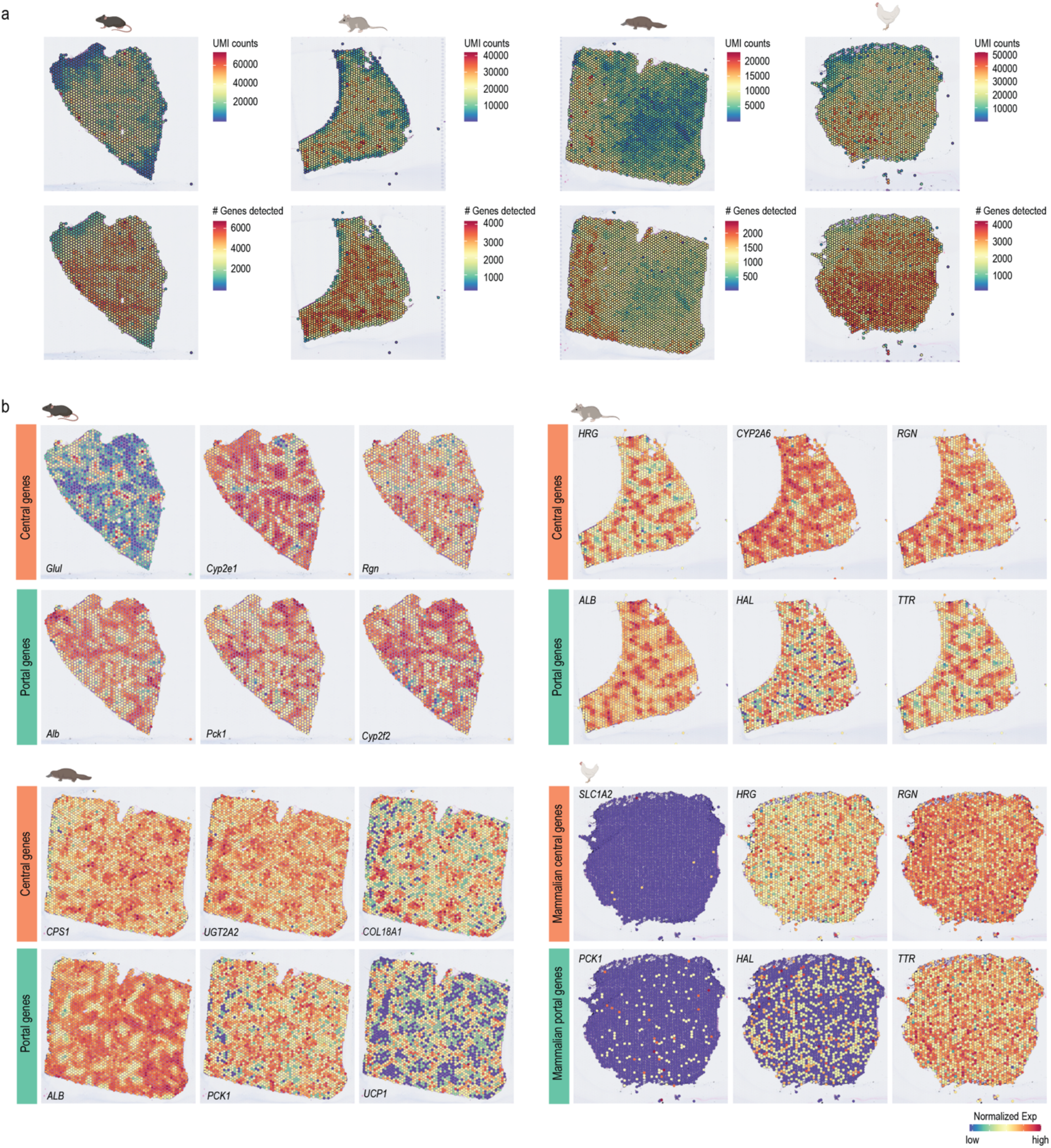
| Visium data quality control. **a**, UMI counts and number of genes detected in mouse, opossum, platypus, and chicken samples in the Visium experiments. **b**, Spatial feature plots showing the expression of the central and portal marker genes across the whole tissue section in mouse, opossum, and platypus, or the orthologs of mammalian central or portal markers in the chicken section.

**Extended Data Fig. 10.**
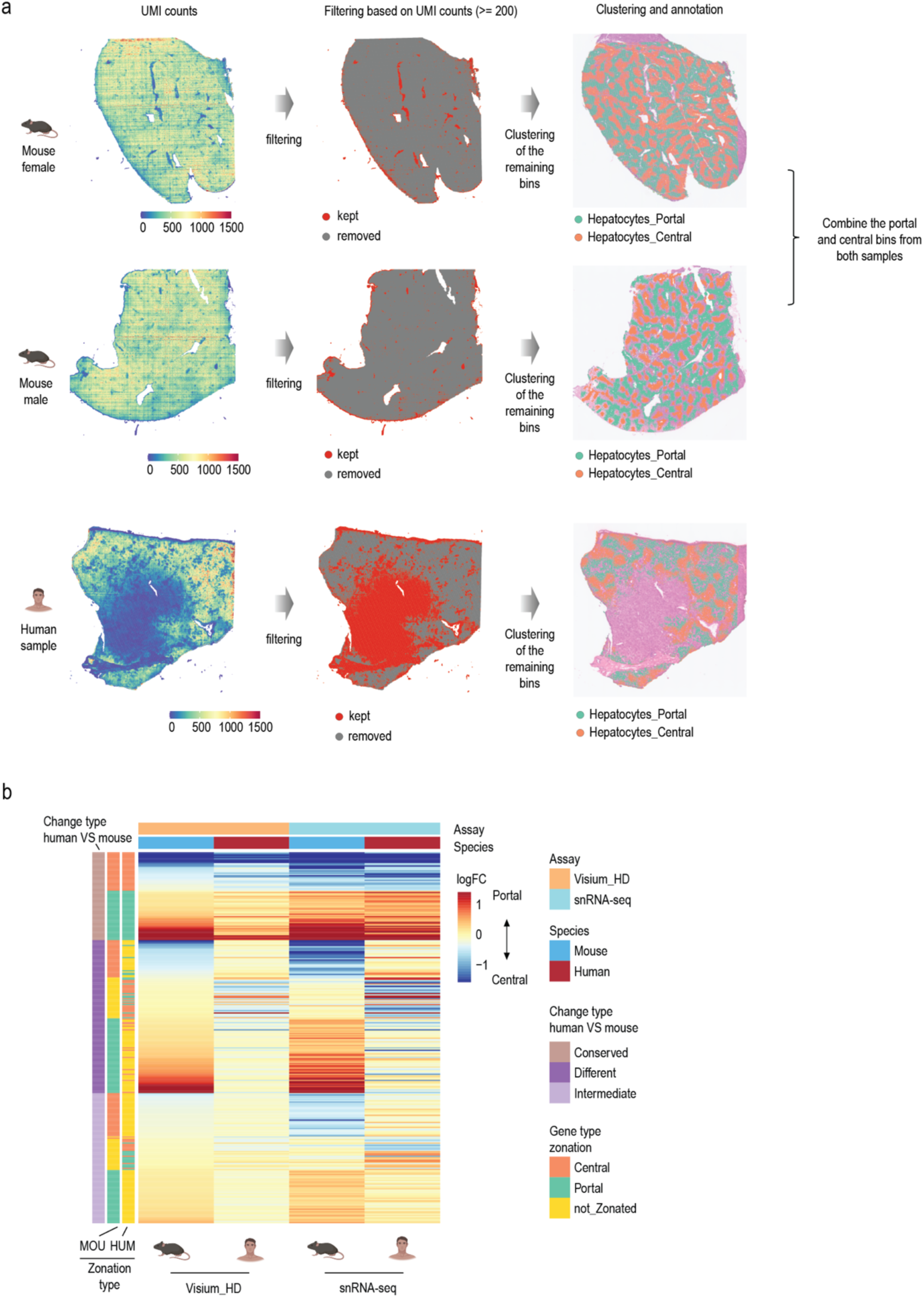
| Comparative analysis of zonated genes between human and mouse. **a**, Spatial map showing the number of UMIs detected in each 8 μm bin (left), the bins kept after UMI-based filtering (UMI counts >= 200 kept, middle), and spatial patterns of the portal and central hepatocyte clusters (right) in mouse and human Visium HD data. **b**, Heatmap showing the zonation patterns of all 1523 1:1 orthologous genes that are high-confidence genes in both human and mouse.

**Extended Data Fig. 11.**
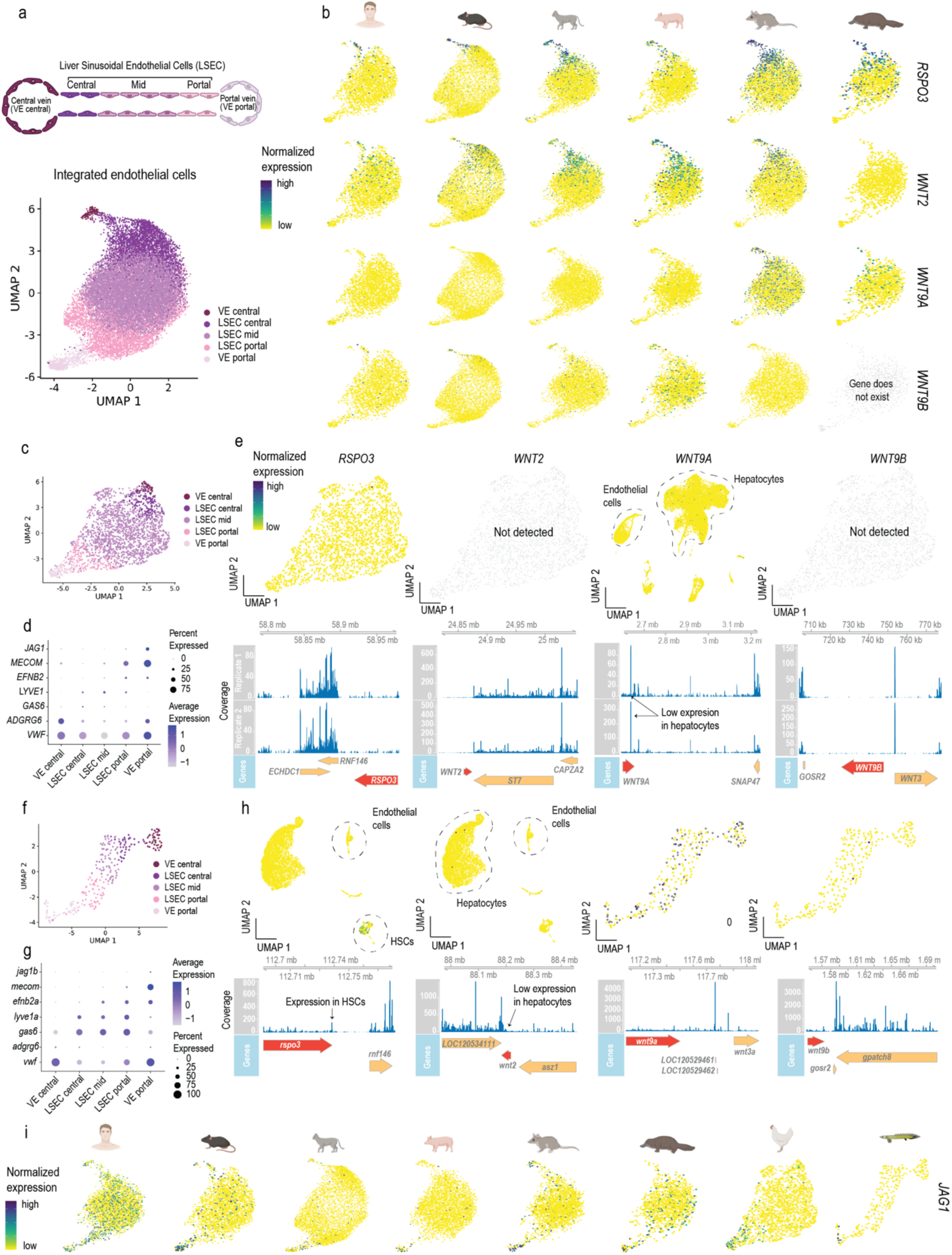
| Endothelial cell annotation and morphogen expression. **a**, UMAP of integrated endothelial cells across human, mouse, cat, pig, opossum and platypus with cells colored by endothelial cell subtype (VE_central: central vein, LSEC_central: central liver sinusoidal endothelial cells, LSEC_mid: mid-lobular liver sinusoidal endothelial cells, LSEC_portal: portal liver sinusoidal endothelial cells, VE_portal: portal vein). **b**, Feature plots showing the expression of *RSPO3, WNT2, WNT9A and WNT9B* in human, mouse, cat, pig, opossum and platypus. **c**, UMAP of chicken endothelial cells subtypes. **d**, Marker genes for endothelial cell subtypes in chicken. **e**, Feature plots showing the expression of *RSPO3, WNT2, WNT9A and WNT9B* in chicken (top) and coverage plots for those same genes in each replicate (bottom). Depending on the gene, either the full dataset embedding or a focused endothelial cell embedding is shown to best illustrate expression patterns. **f**, UMAP of bichir endothelial cells subtypes. **g**, Marker genes for endothelial cell subtypes in bichir. **h**, Feature plots showing the expression of *RSPO3, WNT2, WNT9A, and WNT9B* in bichir (top) and coverage plots for those same genes (bottom). Depending on the gene, either the full dataset embedding or a focused endothelial cell embedding is shown to best illustrate expression patterns. **i**, Feature plots showing the expression of *JAG1* in human, mouse, cat, pig, opossum, platypus, chicken, and bichir.

**Extended Data Fig. 12.**
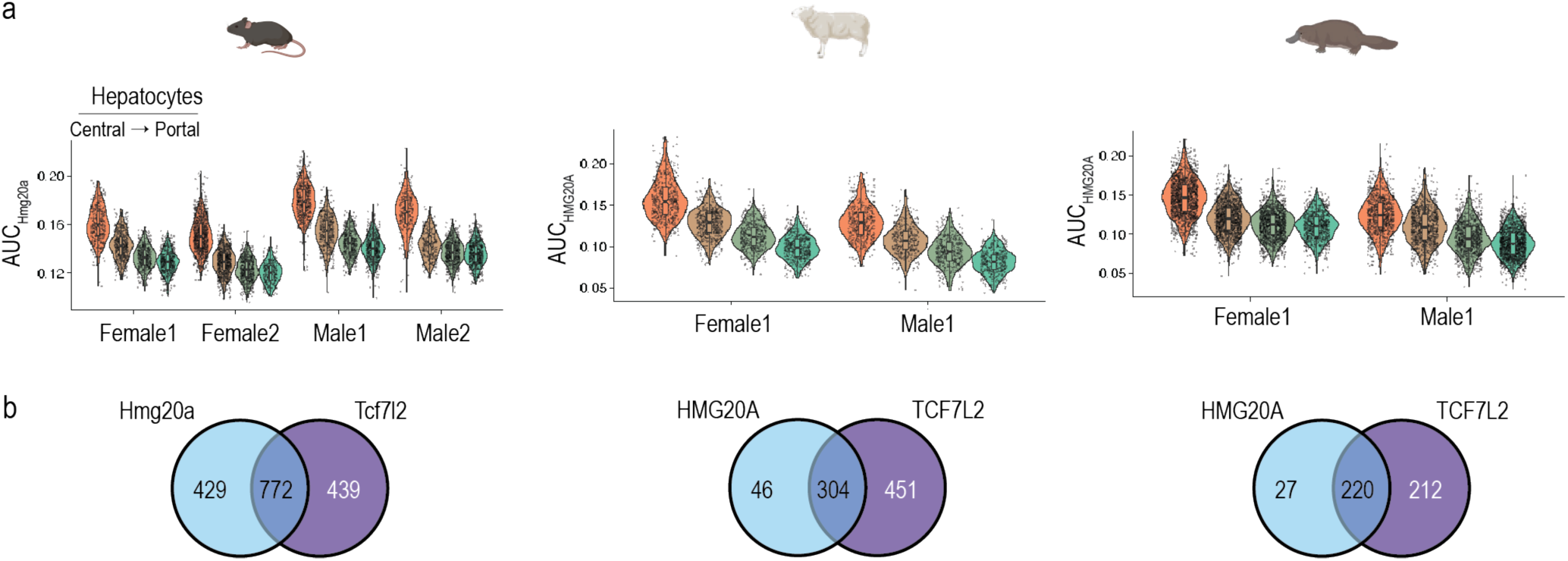
| HMG20A eRegulon. **a**, Activity of HMG20A eRegulon (as AUC: area under the curve) along the different subtypes of hepatocytes ordered from central to portal in mouse, sheep, and platypus. **b**, Intersect of HMG20A target genes with TCF7L2 target genes.

**Extended Data Fig. 13.**
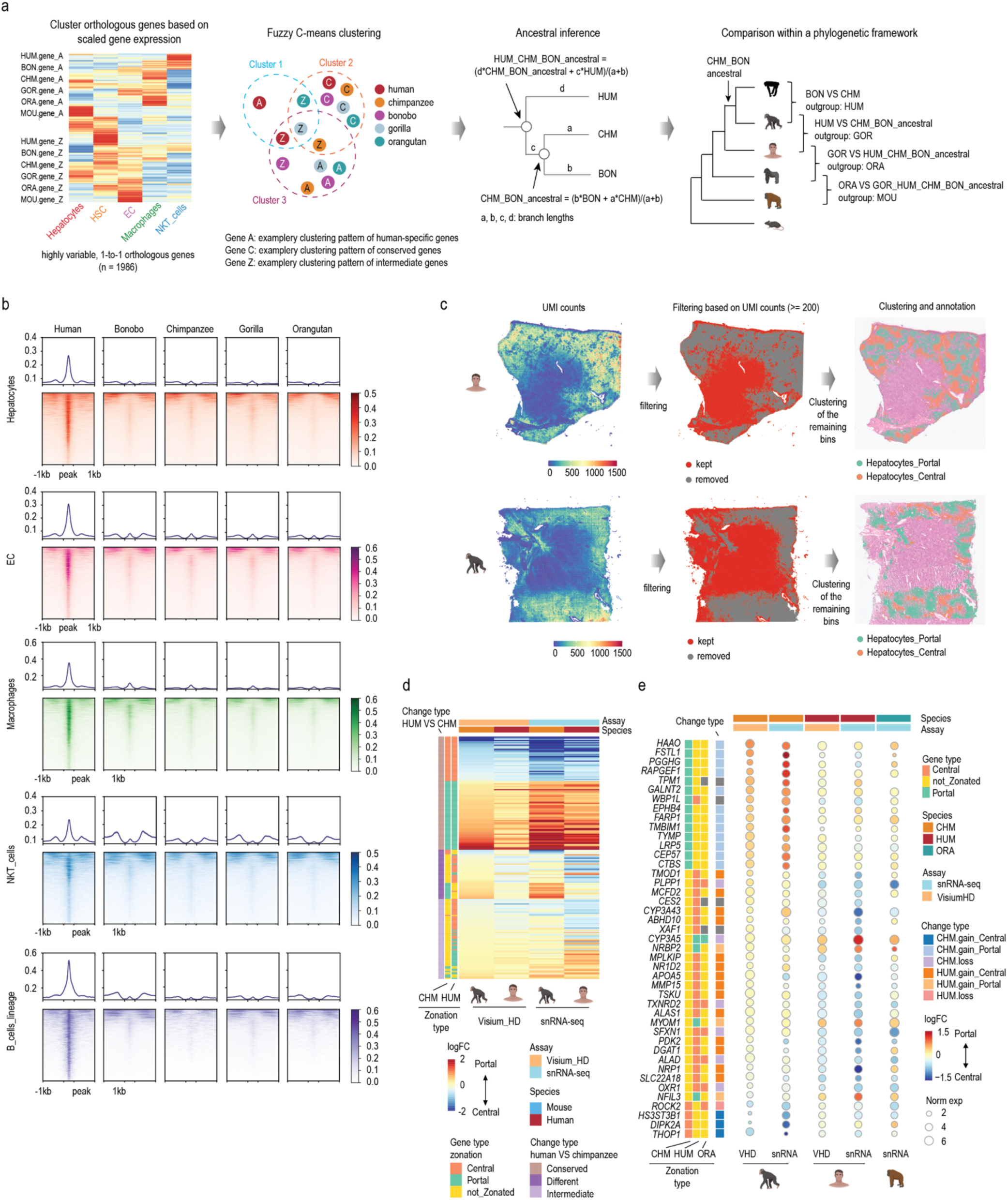
| human-specific molecular changes. **a**, Schematics of the fuzzy C-means clustering approach for identifying genes that switched cell-type specificity patterns within the great ape phylogeny. **b**, Average profiles and heatmaps showing the chromatin accessibility signals in each cell type centered around human-specific putative CREs identified in the given cell type, and the chromatin accessibility signals at the orthologous regions in other great ape species. For each locus, signals in the neighboring region (1 kb extended upstream and downstream) were also plotted for better visualization. **c**, Spatial map showing the number of UMIs detected in each 8 μm bin (left), the bins kept after UMI-based filtering (UMI counts >= 200 kept, middle), and spatial patterns of the portal and central hepatocyte clusters (right) in human (top) and chimpanzee (bottom) Visium HD data. **d**, Heatmap showing the zonation patterns of all 1:1 orthologous genes that are high-confidence genes (n = 2107) in both human and chimpanzee. **e**, Dotplot showing the expression patterns of all species-specific zonated genes identified in chimpanzee-human comparison measured in Visium HD (chimpanzee, human, and orangutan) or snRNA-seq (chimpanzee and human) experiments. The grey color bars indicate that the genes were not robustly detected in orangutan hepatocytes based on the snRNA-seq data (*XAF1*), shared no clear 1-to-1 orthologs with orangutan (*TPM1, CES2*) (Ensembl release 106), or the zonation pattern in orangutan does not match either that in human or chimpanzee (*WBP1L*).

## Supplementary Tables

Supplementary Table 1: Sample information

Supplementary Table 2: Liver-specific transcriptome annotation

Supplementary Table 3: Quality metrics of single-nucleus RNA-seq libraries

Supplementary Table 4: Quality metrics of single-nucleus ATAC-seq libraries

Supplementary Table 5: Statistical tests and *p*-values in evolutionary rate analyses.

Supplementary Tables 6–21: Zonation scores per hepatocyte cell for armadillo, bonobo, cat, chimpanzee, dog, gorilla, guinea pig, human, mouse, opossum, orangutan, pig, platypus, rabbit, sheep, and tenrec, respectively.

Supplementary Tables 22–37: EdgeR results for assessing zonated gene expression in armadillo, bonobo, cat, chimpanzee, dog, gorilla, guinea pig, human, mouse, opossum, orangutan, pig, platypus, rabbit, sheep, and tenrec, respectively.

Supplementary Table 38: List of 1:1 orthologs that show zonation changes (from portally-enriched to centrally-enriched and vice versa).

Supplementary Table 39: Zonation patterns of the 1:1 orthologous, high-confidence genes (n = 1,523) in the human and mouse comparisons.

Supplementary Table 40: CellPhoneDB results for human, mouse, cat, pig, opossum, and platypus.

Supplementary Table 41–43: Targets of TCF7L2 (from the eRegulons) in mouse, sheep, and platypus, respectively.

Supplementary Table 44: Genes with cell-type specific expression changes within the great ape phylogeny.

Supplementary Table 45: Genes with conserved cell-type specific expression patterns within the great ape lineage.

Supplementary Table 46: Zonation patterns of the 1:1 orthologous high-confidence genes (n = 2,107) in human and chimpanzee comparisons

Supplementary Tables 47–62: Set of marker genes used to calculate zonation scores in armadillo, bonobo, cat, chimpanzee, dog, gorilla, guinea pig, human, mouse, opossum, orangutan, pig, platypus, rabbit, sheep, and tenrec, respectively.

## References

1. Trefts, E., Gannon, M. & Wasserman, D. H. The liver. Current Biology 27, R1147–R1151 (2017).

2. Kalra, A., Yetiskul, E., Wehrle, C. J. & Tuma, F. Physiology, Liver. in StatPearls (StatPearls Publishing, Treasure Island (FL), 2025).

3. Ronca, V. et al. The liver as a central “hub” of the immune system: pathophysiological implications. Physiological Reviews 105, 493–539 (2025).

4. Rui, L. Energy Metabolism in the Liver. Comprehensive Physiology 4, 177–197 (2014).

5. Abumrad, N. A. The Liver as a Hub in Thermogenesis. Cell Metabolism 26, 454–455 (2017).

6. Lerapetritou, M. G., Georgopoulos, P. G., Roth, C. M. & Androulakis, L. P. Tissue-Level Modeling of Xenobiotic Metabolism in Liver: An Emerging Tool for Enabling Clinical Translational Research. Clinical and Translational Science 2, 228–237 (2009).

7. Guschanski, K., Warnefors, M. & Kaessmann, H. The evolution of duplicate gene expression in mammalian organs. Genome Res. 27, 1461–1474 (2017).

8. Cardoso-Moreira, M. et al. Gene expression across mammalian organ development. Nature 571, 505–509 (2019).

9. Wu, B. et al. Single-cell analysis of the amphioxus hepatic caecum and vertebrate liver reveals genetic mechanisms of vertebrate liver evolution. Nat Ecol Evol 8, 1972–1990 (2024).

10. Delgado-Coello, B. Liver regeneration observed across the different classes of vertebrates from an evolutionary perspective. Heliyon 7, e06449 (2021).

11. Zhang, S., Shen, Z. & Li, H. Hepatic caecum of amphioxus and origin of vertebrate liver. Acta Oceanol. Sin. 42, 1–8 (2023).

12. Albadry, M. et al. Cross-species variability in lobular geometry and cytochrome P450 hepatic zonation: insights into CYP1A2, CYP2D6, CYP2E1 and CYP3A4. Front. Pharmacol. 15, (2024).

13. Teutsch, H. F. The modular microarchitecture of human liver. Hepatology 42, 317–325 (2005).

14. Ben-Moshe, S. & Itzkovitz, S. Spatial heterogeneity in the mammalian liver. Nat Rev Gastroenterol Hepatol 16, 395–410 (2019).

15. Ota, N. & Shiojiri, N. Comparative study on a novel lobule structure of the zebrafish liver and that of the mammalian liver. Cell Tissue Res 388, 287–299 (2022).

16. Ota, N. et al. Comparative study on a unique architecture of the brook lamprey liver and that of the hagfish and banded houndshark liver. Cell Tissue Res 398, 93–110 (2024).

17. Shiojiri, N. et al. Phylogenetic analyses of the hepatic architecture in vertebrates. Journal of Anatomy 232, 200–213 (2018).

18. Shiojiri, N. et al. Changes of biliary cilia, smooth muscle tissue distribution, innervation and extracellular matrices during morphological evolution of hepatic architectures in vertebrates. Annals of Anatomy – Anatomischer Anzeiger 250, 152148 (2023).

19. Schmidt, D. et al. Five-Vertebrate ChIP-seq Reveals the Evolutionary Dynamics of Transcription Factor Binding. Science 328, 1036–1040 (2010).

20. Wilson, M. D. et al. Species-Specific Transcription in Mice Carrying Human Chromosome 21. Science 322, 434–438 (2008).

21. Ballester, B. et al. Multi-species, multi-transcription factor binding highlights conserved control of tissue-specific biological pathways. eLife 3, e02626 (2014).

22. Brawand, D. et al. The evolution of gene expression levels in mammalian organs. Nature 478, 343–348 (2011).

23. Villar, D. et al. Enhancer Evolution across 20 Mammalian Species. Cell 160, 554–566 (2015).

24. Groza, T. et al. The International Mouse Phenotyping Consortium: comprehensive knockout phenotyping underpinning the study of human disease. Nucleic Acids Research 51, D1038–D1045 (2023).

25. Karczewski, K. J. et al. The mutational constraint spectrum quantified from variation in 141,456 humans. Nature 581, 434–443 (2020).

26. Sullivan, P. F. et al. Leveraging base-pair mammalian constraint to understand genetic variation and human disease. Science 380, eabn2937 (2023).

27. van der Lee, R., Wiel, L., van Dam, T. J. P. & Huynen, M. A. Genome-scale detection of positive selection in nine primates predicts human-virus evolutionary conflicts. Nucleic Acids Research 45, 10634– 10648 (2017).

28. Kosiol, C. et al. Patterns of Positive Selection in Six Mammalian Genomes. PLOS Genetics 4, e1000144 (2008).

29. Barreiro, L. B. & Quintana-Murci, L. From evolutionary genetics to human immunology: how selection shapes host defence genes. Nat Rev Genet 11, 17–30 (2010).

30. Shultz, A. J. & Sackton, T. B. Immune genes are hotspots of shared positive selection across birds and mammals. eLife 8, e41815 (2019).

31. Stern, D. L. Perspective: Evolutionary developmental Biology and the problem of variation. Evol 54, 1079–1091 (2000).

32. Carroll, S. B. Evolution at Two Levels: On Genes and Form. PLOS Biology 3, e245 (2005).

33. Halpern, K. B. et al. Single-cell spatial reconstruction reveals global division of labour in the mammalian liver. Nature 542, 352–356 (2017).

34. Aizarani, N. et al. A human liver cell atlas reveals heterogeneity and epithelial progenitors. Nature 572, 199–204 (2019).

35. Wagenaar, G. T. et al. Vascular branching pattern and zonation of gene expression in the mammalian liver. A comparative study in rat, mouse, cynomolgus monkey, and pig. Anat Rec 239, 441–452 (1994).

36. Smith, D. D. & Campbell, J. W. Distribution of glutamine synthetase and carbamoyl-phosphate synthetase I in vertebrate liver. Proceedings of the National Academy of Sciences 85, 160–164 (1988).

37. Ota, N., Kato, H. & Shiojiri, N. Gene expression in the liver of the hagfish (Eptatretus burgeri) belonging to the Cyclostomata is ancestral to that of mammals. The Anatomical Record 307, 690–700 (2024).

38. Schär, M., Maly, I. P. & Sasse, D. Histochemical studies on metabolic zonation of the liver in the trout (Salmo gairdneri). Histochemistry 83, 147–151 (1985).

39. MacParland, S. A. et al. Single cell RNA sequencing of human liver reveals distinct intrahepatic macrophage populations. Nat Commun 9, 4383 (2018).

40. Ang, C. H. et al. Self-maintenance of zonal hepatocytes during adult homeostasis and their complex plasticity upon distinct liver injuries. Cell Reports 44, (2025).

41. Droin, C. et al. Space-time logic of liver gene expression at sub-lobular scale. Nat Metab 3, 43–58 (2021).

42. Squair, J. W. et al. Confronting false discoveries in single-cell differential expression. Nat Commun 12, 5692 (2021).

43. Armingol, E. et al. Atlas-scale metabolic activities inferred from single-cell and spatial transcriptomics. 2025.05.09.653038 Preprint at 10.1101/2025.05.09.653038 (2025).

44. Hakvoort, T. B. M. et al. Pivotal role of glutamine synthetase in ammonia detoxification. Hepatology 65, 281 (2017).

45. Planas-Paz, L. et al. The RSPO–LGR4/5–ZNRF3/RNF43 module controls liver zonation and size. Nat Cell Biol 18, 467–479 (2016).

46. Benhamouche, S., et al. *Apc* Tumor Suppressor Gene Is the “Zonation-Keeper” of Mouse Liver. Developmental Cell 10, 759–770 (2006).

47. Halpern, K. B. et al. Paired-cell sequencing enables spatial gene expression mapping of liver endothelial cells. Nat Biotechnol 36, 962–970 (2018).

48. Hu, S. et al. Single-cell spatial transcriptomics reveals a dynamic control of metabolic zonation and liver regeneration by endothelial cell Wnt2 and Wnt9b. Cell Reports Medicine 3, 100754 (2022).

49. Inverso, D. et al. A spatial vascular transcriptomic, proteomic, and phosphoproteomic atlas unveils an angiocrine Tie–Wnt signaling axis in the liver. Dev Cell 56, 1677–1693.e10 (2021).

50. Troul’e, K. et al. CellPhoneDB v5: inferring cell-cell communication from single-cell multiomics data. in (2023).

51. Hofmann, J. J. et al. Jagged1 in the portal vein mesenchyme regulates intrahepatic bile duct development: insights into Alagille syndrome. Development 137, 4061–4072 (2010).

52. Alshamy, Z. et al. Structure and age-dependent growth of the chicken liver together with liver fat quantification: A comparison between a dual-purpose and a broiler chicken line. PLOS ONE 14, e0226903 (2019).

53. Bravo González-Blas, C., et al. Single-cell spatial multi-omics and deep learning dissect enhancer-driven gene regulatory networks in liver zonation. Nat Cell Biol 26, 153–167 (2024).

54. Ayala, I. et al. The Spatial Transcriptional Activity of Hepatic TCF7L2 Regulates Zonated Metabolic Pathways that Contribute to Liver Fibrosis. Nat Commun 16, 3408 (2025).

55. Krawczyk, J. et al. The Diabetes Gene Tcf7l2 Organizes Gene Expression in the Liver and Regulates Amino Acid Metabolism. 2025.04.03.647067 Preprint at 10.1101/2025.04.03.647067 (2025).

56. Brosch, M. et al. Epigenomic map of human liver reveals principles of zonated morphogenic and metabolic control. Nat Commun 9, 4150 (2018).

57. Bravo González-Blas, C., et al. SCENIC+: single-cell multiomic inference of enhancers and gene regulatory networks. Nat Methods 20, 1355–1367 (2023).

58. Mononen, J. et al. Genetic variation is a key determinant of chromatin accessibility and drives differences in the regulatory landscape of C57BL/6J and 129S1/SvImJ mice. Nucleic Acids Res 52, 2904– 2923 (2024).

59. Pontzer, H. et al. Metabolic acceleration and the evolution of human brain size and life history. Nature 533, 390–392 (2016).

60. Zihlman, A. L. & Bolter, D. R. Body composition in Pan paniscus compared with Homo sapiens has implications for changes during human evolution. Proceedings of the National Academy of Sciences 112, 7466–7471 (2015).

61. Watkins, P. A. et al. Identification of differences in human and great ape phytanic acid metabolism that could influence gene expression profiles and physiological functions. BMC Physiology 10, 19 (2010).

62. Rada, P., González-Rodríguez, Á., García-Monzón, C. & Valverde, Á. M. Understanding lipotoxicity in NAFLD pathogenesis: is CD36 a key driver? Cell Death Dis 11, 802 (2020).

63. Xiao, X. et al. Liver ACSM3 deficiency mediates metabolic syndrome via a lauric acid-HNF4α-p38 MAPK axis. The EMBO Journal 43, 507–532 (2024).

64. Villanueva, C. J. et al. Specific role for acyl CoA: Diacylglycerol acyltransferase 1 (: Dgat1:) in hepatic steatosis due to exogenous fatty acids†. Hepatology 50, 434 (2009).

65. Forte, T. M. & Ryan, R. O. Apolipoprotein A5: Extracellular and Intracellular Roles in Triglyceride Metabolism. Current Drug Targets 16, 1274–1280.

66. Cao, Y. et al. ABHD10 is an S-depalmitoylase affecting redox homeostasis through peroxiredoxin-5. Nat Chem Biol 15, 1232–1240 (2019).

67. Go, Y. et al. Inhibition of Pyruvate Dehydrogenase Kinase 2 Protects Against Hepatic Steatosis Through Modulation of Tricarboxylic Acid Cycle Anaplerosis and Ketogenesis. Diabetes 65, 2876–2887 (2016).

68. Hardman, R. C., Volz, D. C., Kullman, S. W. & Hinton, D. E. An in vivo Look at Vertebrate Liver Architecture: Three-Dimensional Reconstructions from Medaka (Oryzias latipes). The Anatomical Record 290, 770–782 (2007).

69. Zhou, Y., Eid, T., Hassel, B. & Danbolt, N. C. Novel aspects of glutamine synthetase in ammonia homeostasis. Neurochemistry International 140, 104809 (2020).

70. Qvartskhava, N. et al. Hyperammonemia in gene-targeted mice lacking functional hepatic glutamine synthetase. Proceedings of the National Academy of Sciences 112, 5521–5526 (2015).

71. Brosnan, M. E. & Brosnan, J. T. Hepatic glutamate metabolism: a tale of 2 hepatocytes. Am J Clin Nutr 90, 857S–861S (2009).

72. Bartl, M. et al. Optimality in the zonation of ammonia detoxification in rodent liver. Arch Toxicol 89, 2069–2078 (2015).

73. Häussinger, D. Hepatocyte Heterogeneity in Glutamine and Ammonia Metabolism and the Role of an Intercellular Glutamine Cycle during Ureogenesis in Perfused Rat Liver. European Journal of Biochemistry 133, 269–275 (1983).

74. Deuel, T. F., Louie, M. & Lerner, A. Glutamine synthetase from rat liver. Purification, properties, and preparation of specific antisera. J Biol Chem 253, 6111–6118 (1978).

75. Ip, A. Y. K. & Chew, S. F. Ammonia Production, Excretion, Toxicity, and Defense in Fish: A Review. Front. Physiol. 1, (2010).

76. Wilkie, M. P. Ammonia excretion and urea handling by fish gills: present understanding and future research challenges. J Exp Zool 293, 284–301 (2002).

77. Cragg, M. M., Balinsky, J. B. & Baldwin, E. A comparative study of nitrogen excretion in some Amphibia and reptiles. Comp Biochem Physiol 3, 227–235 (1961).

78. Wood, C. M., Munger, R. S. & Toews, D. P. Ammonia, Urea and H+ Distribution and the Evolution of Ureotelism in Amphibians. Journal of Experimental Biology 144, 215–233 (1989).

79. Méndez-Narváez, J. & Warkentin, K. M. Reproductive colonization of land by frogs: Embryos and larvae excrete urea to avoid ammonia toxicity. Ecol Evol 12, e8570 (2022).

80. Campbell, J. W., Vorhaben, J. E. & Smith Jr., D. D. Uricoteley: Its nature and origin during the evolution of tetrapod vertebrates. Journal of Experimental Zoology 243, 349–363 (1987).

81. Lenártová, V., Holovská, K., Rosival, I. & Havassy, I. Glutamate dehydrogenase and glutamine synthetase activity in some organs of ruminants and monogastric animals. Physiol Bohemoslov 26, 535–542 (1977).

82. Vollmers, C. et al. Time of feeding and the intrinsic circadian clock drive rhythms in hepatic gene expression. Proceedings of the National Academy of Sciences 106, 21453–21458 (2009).

83. Guan, D. et al. The hepatocyte clock and feeding control chronophysiology of multiple liver cell types. Science 369, 1388–1394 (2020).

84. Fernández-Pérez, L. et al. Control of Liver Gene Expression by Sex Steroids and Growth Hormone Interplay. in Chemistry and Biological Activity of Steroids (IntechOpen, 2019). doi:10.5772/intechopen.86611.

85. Rodríguez-Montes, L. et al. Sex-biased gene expression across mammalian organ development and evolution. Science 382, eadf1046 (2023).

86. Waxman, D. J. & O’Connor, C. Growth Hormone Regulation of Sex-Dependent Liver Gene Expression. Mol Endocrinol 20, 2613–2629 (2006).

87. Wilson, C. G. et al. Hepatocyte-Specific Disruption of CD36 Attenuates Fatty Liver and Improves Insulin Sensitivity in HFD-Fed Mice. Endocrinology 157, 570–585 (2016).

88. Zeng, H. et al. CD36 promotes de novo lipogenesis in hepatocytes through INSIG2-dependent SREBP1 processing. Molecular Metabolism 57, 101428 (2022).

89. Liu, Y. & Yin, W. CD36 in liver diseases. Hepatology Communications 9, e0623 (2025).

90. Son, N.-H. et al. Endothelial cell CD36 optimizes tissue fatty acid uptake. J Clin Invest 128, 4329–4342 (2018).

91. Neel, J. V. Diabetes Mellitus: A “Thrifty” Genotype Rendered Detrimental by “Progress”? Am J Hum Genet 14, 353–362 (1962).

92. Krishnaswami, S. R. et al. Using single nuclei for RNA-seq to capture the transcriptome of postmortem neurons. Nat Protoc 11, 499–524 (2016).

93. Chen, D. et al. Single cell atlas for 11 non-model mammals, reptiles and birds. Nat Commun 12, 7083 (2021).

94. Wildman, D. E., Uddin, M., Liu, G., Grossman, L. I. & Goodman, M. Implications of natural selection in shaping 99.4% nonsynonymous DNA identity between humans and chimpanzees: Enlarging genus Homo. Proceedings of the National Academy of Sciences 100, 7181–7188 (2003).

95. Marin, R. et al. Convergent origination of a Drosophila-like dosage compensation mechanism in a reptile lineage. Genome Res. 27, 1974–1987 (2017).

96. Wang, Z.-Y. et al. Transcriptome and translatome co-evolution in mammals. Nature 588, 642–647 (2020).

97. Bolger, A. M., Lohse, M. & Usadel, B. Trimmomatic: a flexible trimmer for Illumina sequence data. Bioinformatics 30, 2114–2120 (2014).

98. Kaminow, B., Yunusov, D. & Dobin, A. STARsolo: accurate, fast and versatile mapping/quantification of single-cell and single-nucleus RNA-seq data. 2021.05.05.442755 Preprint at 10.1101/2021.05.05.442755 (2021).

99. Kovaka, S. et al. Transcriptome assembly from long-read RNA-seq alignments with StringTie2. Genome Biology 20, 278 (2019).

100. Trapnell, C. et al. Transcript assembly and quantification by RNA-Seq reveals unannotated transcripts and isoform switching during cell differentiation. Nat Biotechnol 28, 511–515 (2010).

101. Zolotarov, G., Grau-Bové, X. & Sebé-Pedrós, A. GeneExt: a gene model extension tool for enhanced single-cell RNA-seq analysis. 2023.12.05.570120 Preprint at 10.1101/2023.12.05.570120 (2023).

102. Granja, J. M. et al. ArchR is a scalable software package for integrative single-cell chromatin accessibility analysis. Nat Genet 53, 403–411 (2021).

103. Zaremba, B. et al. Developmental origins and evolution of pallial cell types and structures in birds. Science 387, eadp5182 (2025).

104. Murat, F. et al. The molecular evolution of spermatogenesis across mammals. Nature 613, 308–316 (2023).

105. Wolock, S. L., Lopez, R. & Klein, A. M. Scrublet: Computational Identification of Cell Doublets in Single-Cell Transcriptomic Data. Cell Systems 8, 281–291.e9 (2019).

106. Hao, Y. et al. Integrated analysis of multimodal single-cell data. Cell 184, 3573–3587.e29 (2021).

107. Nault, R., Fader, K. A., Bhattacharya, S. & Zacharewski, T. R. Single-Nuclei RNA Sequencing Assessment of the Hepatic Effects of 2,3,7,8-Tetrachlorodibenzo-p-dioxin. Cellular and Molecular Gastroenterology and Hepatology 11, 147–159 (2021).

108. Korsunsky, I. et al. Fast, sensitive and accurate integration of single-cell data with Harmony. Nat Methods 16, 1289–1296 (2019).

109. Zhang, Y. et al. Model-based Analysis of ChIP-Seq (MACS). Genome Biology 9, R137 (2008).

110. Granja, J. M. et al. Single-cell multiomic analysis identifies regulatory programs in mixed-phenotype acute leukemia. Nat Biotechnol 37, 1458–1465 (2019).

111. Fornes, O. et al. JASPAR 2020: update of the open-access database of transcription factor binding profiles. Nucleic Acids Research 48, D87–D92 (2020).

112. Emms, D. M. & Kelly, S. OrthoFinder: phylogenetic orthology inference for comparative genomics. Genome Biology 20, 238 (2019).

113. Kumar, S. et al. TimeTree 5: An Expanded Resource for Species Divergence Times. Molecular Biology and Evolution 39, msac174 (2022).

114. Wilman, H. et al. EltonTraits 1.0: Species-level foraging attributes of the world’s birds and mammals. Ecology 95, 2027–2027 (2014).

115. Chen, Y., Chen, L., Lun, A. T. L., Baldoni, P. L. & Smyth, G. K. edgeR v4: powerful differential analysis of sequencing data with expanded functionality and improved support for small counts and larger datasets. Nucleic Acids Research 53, gkaf018 (2025).

116. Sepp, M. et al. Cellular development and evolution of the mammalian cerebellum. Nature 625, 788–796 (2024).

117. R Core Team. R: A Language and Environment for Statistical Computing. (2013).

118. Paradis, E., Claude, J. & Strimmer, K. APE: Analyses of Phylogenetics and Evolution in R language. Bioinformatics 20, 289–290 (2004).

119. Dickinson, M. E. et al. High-throughput discovery of novel developmental phenotypes. Nature 537, 508–514 (2016).

120. Shao, Y. et al. GenTree, an integrated resource for analyzing the evolution and function of primate-specific coding genes. Genome Res. 29, 682–696 (2019).

121. Lonsdale, J. et al. The Genotype-Tissue Expression (GTEx) project. Nat Genet 45, 580–585 (2013).

122. Palmer, D., Fabris, F., Doherty, A., Freitas, A. A. & Magalhães, J. P. de. Ageing transcriptome meta-analysis reveals similarities and differences between key mammalian tissues. Aging 13, 3313–3341 (2021).

123. Karlsson, M. et al. A single–cell type transcriptomics map of human tissues. Science Advances 7, eabh2169 (2021).

124. Yanai, I. et al. Genome-wide midrange transcription profiles reveal expression level relationships in human tissue specification. Bioinformatics 21, 650–659 (2005).

125. Siepel, A. et al. Evolutionarily conserved elements in vertebrate, insect, worm, and yeast genomes. Genome Res. 15, 1034–1050 (2005).

126. Davydov, E. V. et al. Identifying a High Fraction of the Human Genome to be under Selective Constraint Using GERP++. PLOS Computational Biology 6, e1001025 (2010).

127. Genereux, D. P. et al. A comparative genomics multitool for scientific discovery and conservation. Nature 587, 240–245 (2020).

128. Hickey, G., Paten, B., Earl, D., Zerbino, D. & Haussler, D. HAL: a hierarchical format for storing and analyzing multiple genome alignments. Bioinformatics 29, 1341–1342 (2013).

129. Ramírez, F. et al. deepTools2: a next generation web server for deep-sequencing data analysis. Nucleic Acids Research 44, W160–W165 (2016).

130. Sarropoulos, I. et al. Developmental and evolutionary dynamics of cis-regulatory elements in mouse cerebellar cells. Science 373, eabg4696 (2021).

131. Hinrichs, A. S. et al. The UCSC Genome Browser Database: update 2006. Nucleic Acids Research 34, D590–D598 (2006).

132. Griffiths, J. A., Richard, A. C., Bach, K., Lun, A. T. L. & Marioni, J. C. Detection and removal of barcode swapping in single-cell RNA-seq data. Nat Commun 9, 2667 (2018).

133. Moses, L. et al. Voyager: exploratory single-cell genomics data analysis with geospatial statistics. bioRxiv 2023.07.20.549945 (2023) doi:10.1101/2023.07.20.549945.

134. Angerer, P. et al. destiny: diffusion maps for large-scale single-cell data in R. Bioinformatics 32, 1241– 1243 (2016).

135. Hao, Y. et al. Dictionary learning for integrative, multimodal and scalable single-cell analysis. Nat Biotechnol 42, 293–304 (2024).

136. Kolberg, L. & Raudvere, U. gprofiler2: Interface to the ‘g:Profiler’ Toolset. (2021).

137. Swainston, N. et al. Recon 2.2: from reconstruction to model of human metabolism. Metabolomics 12, 109 (2016).

138. Van de Sande, B. et al. A scalable SCENIC workflow for single-cell gene regulatory network analysis. Nat Protoc 15, 2247–2276 (2020).

139. Navarro Gonzalez, J., et al. The UCSC Genome Browser database: 2021 update. Nucleic Acids Research 49, D1046–D1057 (2021).

140. Lawrence, M. et al. Software for Computing and Annotating Genomic Ranges. PLOS Computational Biology 9, e1003118 (2013).

141. Dias, M., Florêncio, A. & dirk. omadson/fuzzy-c-means: v1.2.2. Zenodo 10.5281/zenodo.4310027 (2020).

142. Zhang, X., Kaplow, I. M., Wirthlin, M., Park, T. Y. & Pfenning, A. R. HALPER facilitates the identification of regulatory element orthologs across species. Bioinformatics 36, 4339–4340 (2020).

143. Quinlan, A. R. & Hall, I. M. BEDTools: a flexible suite of utilities for comparing genomic features. Bioinformatics 26, 841–842 (2010).

144. Tanigawa, Y., Dyer, E. S. & Bejerano, G. WhichTF is functionally important in your open chromatin data? PLOS Computational Biology 18, e1010378 (2022).

